# Higher-order combinatorial chromatin perturbations by engineered CRISPR-Cas12a for functional genomics

**DOI:** 10.1101/2023.09.18.558350

**Authors:** CC Hsiung, CM Wilson, NA Sambold, R Dai, Q Chen, S Misiukiewicz, A Arab, N Teyssier, T O’Loughlin, JC Cofsky, J Shi, LA Gilbert

## Abstract

Multiplexed genetic perturbations are critical for testing functional interactions among coding or non-coding genetic elements. Compared to double-stranded DNA cutting, repressive chromatin formation using CRISPR interference (CRISPRi) avoids genotoxicity and is more effective for perturbing non-coding regulatory elements in pooled assays. However, current CRISPRi pooled screening approaches are limited to targeting 1-3 genomic sites per cell. To develop a tool for higher-order (≥3) combinatorial targeting of genomic sites with CRISPRi in functional genomics screens, we engineered an *Acidaminococcus* Cas12a variant -- referred to as <u>mul</u>tiplexed transcriptional interference AsCas12a (multiAsCas12a). multiAsCas12a incorporates a key mutation, R1226A, motivated by the hypothesis of nicking-induced stabilization of the ribonucleoprotein:DNA complex for improving CRISPRi activity. multiAsCas12a significantly outperforms prior state-of-the-art Cas12a variants in combinatorial CRISPRi targeting using high-order multiplexed arrays of lentivirally transduced CRISPR RNAs (crRNA), including in high-throughput pooled screens using 6-plex crRNA array libraries. Using multiAsCas12a CRISPRi, we discover new enhancer elements and dissect the combinatorial function of cis-regulatory elements. These results instantiate a group testing framework for efficiently surveying potentially numerous combinations of chromatin perturbations for biological discovery and engineering.

## Introduction

Functional interactions among combinations of genetic elements underlie many natural and engineered phenotypes ^1–3^, often involving higher-order (≥3-plex) combinations of coding ^4,5^ or non-coding elements ^6–9^. Experimentally testing higher-order combinations of genetic perturbations has been limited by throughput, with prior systematic analyses primarily performed in yeast ^10–14^. In mammalian functional genomics, Cas9-based pooled CRISPR screens ^15,16^ using sequencing readouts have been limited in multiplexing capability, with only a few studies targeting more than 2 genomic sites per cell ^17,18^. Further multiplexing in Cas9-based pooled screening is challenging due to 1) increasingly complex cloning schemes for large constructs encoding multiple guides, each expressed from a separate promoter ^17–19^, and 2) length-dependent high recombination frequencies of lentiviral guide libraries ^20–22^. Conceptually, it also remains unclear how to tractably survey the potentially vast combinatorial spaces for ≥3-plex perturbations.

Cas12a, a member of the type V CRISPR-Cas family, has been proposed as an alternative to Cas9 for genetic perturbations due to enhanced multiplexing capabilities. Cas12a harbors RNase activity, separable from its DNase activity, that can process a compact primary transcript expressed from a single promoter into multiple CRISPR RNAs (crRNA) ^23,24^. An array of multiple Cas12a crRNAs, each composed of 19nt direct repeat and 19-23nt spacer, can be encoded on a chemically synthesized oligo for single-step cloning into an expression vector ^25–29^. Cas12a has been engineered for mammalian cell applications using its DNase activity to disrupt coding gene function using single or multiplexed crRNA constructs in individual well-based assays ^24–27,30–32^ and in pooled sequencing screens ^28,29,32–36^. However, extended multiplexing with fully DNase-competent Cas12a is expected to be constrained by genotoxicity from double-stranded DNA breaks in many biological contexts ^28,37,38,39–42^. In principle, avoiding genotoxicity can be achieved by using DNase-dead Cas fusion proteins to control chromatin state, such as by dCas9-based transcriptional repression (CRISPRi) and activation (CRISPRa)^43–45^. Moreover, CRISPRi is more efficient than DNA cutting at perturbing enhancers in pooled screens ^46–48^, likely due to CRISPRi’s larger genomic window of activity via formation of repressive chromatin ^49^. Thus, a DNase-dead Cas12a (dCas12a) functional genomics platform for multi-site CRISPRi targeting would be highly desirable for testing the combinatorial functions of coding and non-coding genetic elements. However, no dCas12a-based pooled CRISPRi screening platform has been reported. Several studies have used dCas12a fusion proteins for CRISPRi in human cells in individual well-based assays, reporting either successful ^27,50–52^ or unsuccessful ^53^ repression of target genes. These dCas12a CRISPRi studies delivered crRNA plasmids by transient transfection, rather than lentiviral transduction. Transient plasmid transfections express synthetic components at 10 to 1000-fold higher than single-copy lentiviral integration of crRNA constructs, which is required in pooled screens to attribute cellular phenotypes to unique crRNA constructs by high-throughput sequencing ^15,16^. Whether prior dCas12a CRISPRi constructs are sufficiently potent for pooled screens remains unclear.

In this study, we show that existing dCas12a CRISPRi fusion constructs function poorly when used with limiting doses of lentivirally delivered components, thus precluding their application in pooled screens. We engineered a new Cas12a variant that incorporates a key mutation, R1226A, which enhances stability of the ribonucleoprotein:DNA complex in the form of a nicked DNA intermediate in vitro ^54,55^. We show that in human cells, R1226A significantly improves CRISPRi activity in the setting of lentivirally delivered crRNA constructs, enabling use of 6-plex crRNA arrays in high-throughput pooled screens using 6-plex crRNA arrays and up to 10-plex crRNA arrays in well-based assays. We use this combinatorial CRISPRi platform to efficiently discover new enhancer elements and to test higher-order combinatorial perturbations of cis-regulatory elements. These results instantiate a group testing framework that enables efficient searches of potentially large combinatorial spaces of chromatin perturbations.

## Results

### CRISPRi using state-of-the-art dAsCas12a fusion proteins is dose-limited and hypoactive in the setting of lentivirally delivered components

We focused on building a CRISPRi functional genomics platform using *Acidaminococcus* Cas12a (AsCas12a), the only Cas12a ortholog with demonstrated success in pooled screens in mammalian cells ^28,29,32–35,56,57^. A previous study reported using dAsCas12a for CRISPRi by plasmid transient transfection delivery of dAsCas12a-KRABx3 protein (harboring the E993A DNase-dead mutation) and crRNA in HEK 293T cells (Campa et al. 2019). To test this construct in the setting of lentivirally delivered crRNA, we introduced dAsCas12a-KRABx3 by piggyBac transposition in K562 cells, followed by lentiviral transduction of single crRNA constructs targeting canonical TTTV protospacer adjacent motifs (PAM) proximal to transcriptional start sites of 4 cell surface genes, 3 of which (CD55, CD81 and B2M) have been successfully knocked down by dCas9-KRAB CRISPRi ^58^. Throughout this study we encoded crRNAs in a previously optimized CROP-seq lentiviral vector ^29,59^. We observed no expression change in any of the targeted genes (Fig. 1A-B and Fig. S1-S2). We confirmed the expression of dAsCas12a-KRABx3 by western blot (Fig. S3) and by flow cytometry monitoring of the in-frame P2A-BFP (Fig. S4A). We also observed this lack of CRISPRi activity for dAsCas12a-KRABx3 using lentivirally transduced crRNAs in C4-2B prostate cancer cells (Fig. S4B). In contrast, transient co-transfection of dAsCas12a-KRABx3 and CD55-targeting crRNA plasmids shows modest CRISPRi knockdown in HEK 293T cells (Fig. S5), consistent with prior work ^27^. These findings indicate that the requirements for CRISPRi activity using dAsCas12a-KRABx3 with lentiviral crRNA constructs are distinct from those of plasmid transient transfection in HEK 293T cells ^27^.

**Figure 1.**
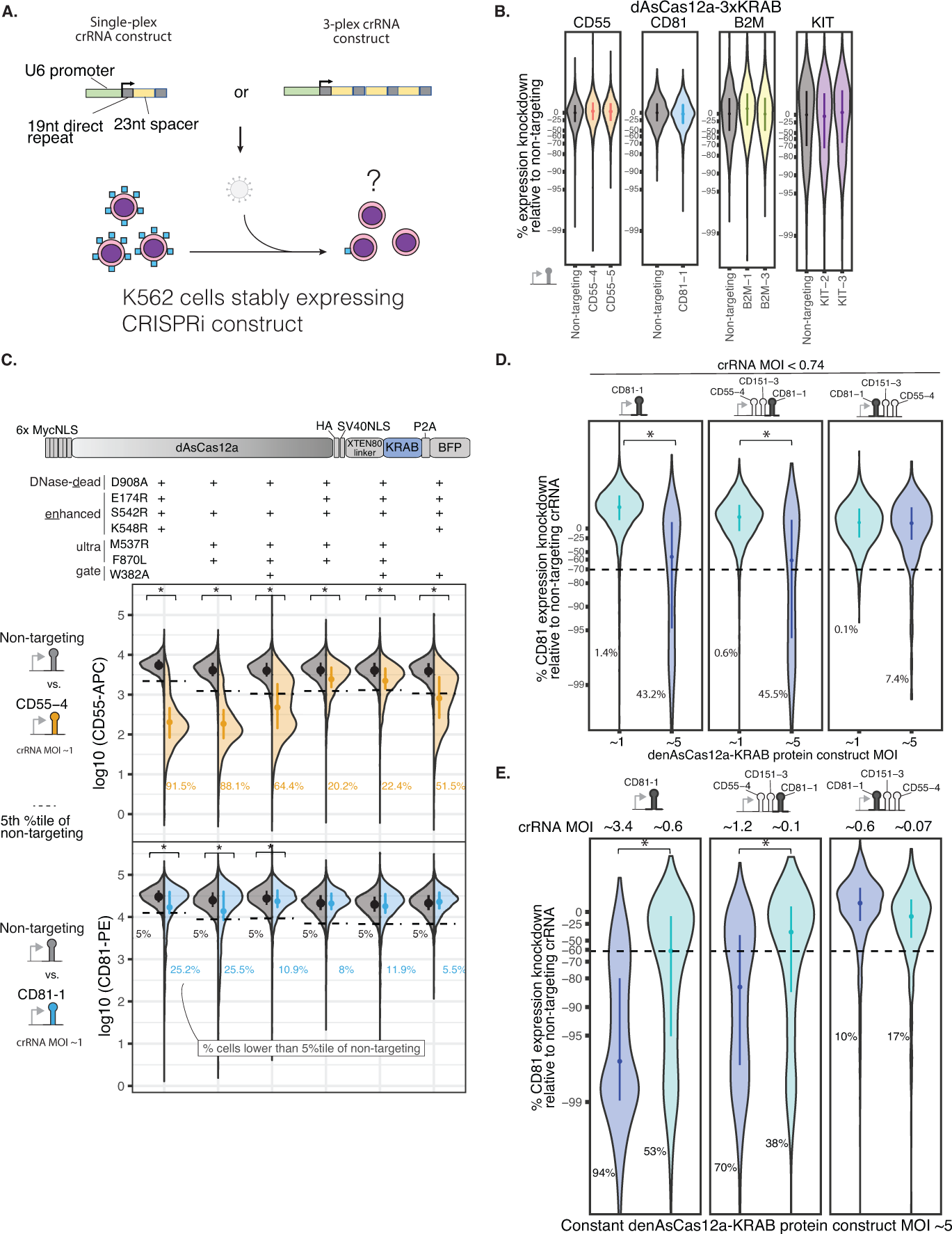
– dAsCas12a-KRAB variants are dose-limited and weak in CRISPRi activity when delivered lentivirally, despite incorporating state-of-the-art optimizations. **A)** Schematic for assaying CRISPRi activity of Cas12a constructs using lentivirally transduced single-plex or 3-plex crRNAs targeting cell surface marker genes assayed by antibody staining and flow cytometry. **B)** K562 cells constitutively expressing dAsCas12a-KRABx3 (Campa et al., 2019) were lentivirally transduced with single crRNAs targeting CD55, CD81, B2M, KIT, or a non-targeting crRNA, and assayed by flow cytometry 6 days after crRNA transduction. **C)** A panel of Cas12a variants harboring combinations of mutations are tested using crCD55-4 and crCD81-1 using the fusion protein domain architecture shown. Both Cas12a fusion protein and crRNA constructs are delivered by lentiviral transduction. D908A is a mutation in the RuvC catalytic triad that renders Cas12a DNase-inactive (Yamano et al., 2016; Zetsche et al., 2015). Other mutations are described in detail in the main text. Shown are single-cell distributions of target gene expression assayed by flow cytometry 6 days after crRNA transduction for one of 3 independent replicates. One-sided Wilcoxon rank-sum test was performed comparing the single-cell distributions of the non-targeting control vs. the corresponding targeting crRNA; asterisk indicates p*<*0.01. Additional replicates and results for additional crRNA constructs (up to 3-plex crRNA constructs) are summarized in Fig. S6. **D)** Analysis of CD81 knockdown in cells lentivirally transduced with denAsCas12a-KRAB protein construct at MOI ∼1 vs. MOI ∼5, while maintaining constant crRNA MOI (*<*0.74) for each crRNA construct. CD81 expression was assayed by flow cytometry 6 days after crRNA transduction. Shown are single-cell distributions of target gene expression knockdown as a percentage of non-targeting control for one of 3-6 biological replicates for each crRNA construct. Median and interquartile range are shown for each distribution. Percentage of cells below the 5th percentile (dashed line) of non-targeting crRNA are shown. One-sided Wilcoxon rank-sum test was performed on single-cell distributions for this replicate; asterisk indicates p*<*0.01. Summaries of all replicates shown in Fig. S7A. **E)** Similar to D, but maintaining constant denAsCas12a-KRAB protein construct MOI at ∼5, while crRNA MOI is changed from high to low as indicated. CD81 expression was assayed by flow cytometry 10 days after crRNA transduction. Shown are single-cell distributions of CD81 knockdown for one of two biological replicates. One-sided Wilcoxon rank-sum test was performed on single-cell distributions for each replicate; asterisk indicates p*<*0.01. Additional replicate shown in Fig. S7B.

In an attempt to overcome this lack of CRISPRi activity, we tested combinations of several Cas12a mutations representing state-of-the-art optimizations of Cas12a. These include: 1) E174R/S542R/K548R (enhanced AsCas12a, <u>en</u>AsCas12a)^31^; 2) M537R/F870L (AsCas12a ultra)^60^; 3) W382A, a mutation that reduces R-loop dissociation in vitro for an orthologous enzyme (*Lachnospiraceae* Cas12a W355A)^61^, but which has not yet been tested in cells. We generated six dAsCas12a variants that each harbor the DNase-inactivating D908A mutation, plus a select combination of the aforementioned mutations. We delivered these variants in K562 cells by stable lentiviral expression, followed by lentiviral transduction of crRNA construct targeting the TSS of CD55 or CD81 (Fig. 1C and Fig. S6). Among this panel, denAsCas12a-KRAB (E174R/S542R/K548R, plus D908A DNase-dead mutation) performed the best and demonstrated strong repression of CD55. However, even for this best construct, we observed weak repression of CD81, indicating inconsistent performance across crRNAs (Fig. 1C and Fig. S6).

Dose-response and construct potency are key considerations for multiplexed applications, as increased multiplexing effectively reduces the Cas protein available to bind each individual crRNA. Focusing on denAsCas12a-KRAB as the top variant, we tested the effect of separately altering the dosage of Cas12a protein and crRNAs. We found that increasing the MOI of the denAsCas12a-KRAB construct from ∼1 to ∼5 can improve CRISPRi knockdown by some crRNA constructs (Fig. 1D and Fig. S7A). However, CRISPRi activity of denAsCas12a-KRAB is lost when the crRNA MOI is reduced to <1 to mimic the low MOI required for pooled screens (Fig. 1E and Fig. S7B). More problematically, CD81 knockdown by a 3-plex crRNA (CD81-1_CD151-3_CD55-4) is extremely weak (∼0%-25% median expression knockdown relative to non-targeting control) across all doses of protein (Fig. 1D and Fig. S7A) and crRNA (Fig. 1E and Fig. S7B) tested.

Given the inconsistent and deficient performance of denAsCas12a-KRAB, we tested an alternative CRISPRi approach without mutating the RuvC DNase active site. In the setting of transient plasmid transfection delivery in HEK 293T cells, wild-type AsCas12a has been used for transcriptional control with truncated (15nt) crRNA spacers, which enable DNA binding but not cleavage ^26,27^. We tested this approach by fusing KRAB or KRABx3 to opAsCas12a, a fully DNase-active Cas12a optimized for pooled screens ^29^. We confirmed that 15nt spacers do not support DNA cleavage, while 23nt spacers do (Fig. S8). However, using 15nt spacers, we observed weak or no CRISPRi activity in two cell lines (Fig. S8). In total, we demonstrated 3 separate approaches to abolish the DNase activity of AsCas12a (E993A in Fig. 1B and Fig. S4; D908A in Fig. 1C-E and Fig. S7B; and truncated spacers in Fig. S8) result in poor CRISPRi activity using lentivirally transduced crRNA constructs.

### multiAsCas12a-KRAB (R1226A/E174R/S542R/K548R), a variant that favors a nicked DNA intermediate, substantially improves lentivirally delivered CRISPRi

The mediocre performance of dAsCas12a for CRISPRi surprised us given the success of AsCas12a in DNA-cutting pooled screens ^28,29,35,36^. We wondered whether full inactivation of DNA cutting in dAsCas12a may preclude strong CRISPRi activity by reducing DNA affinity. Previous studies indicate that DNA cleavage strengthens the Cas12a:DNA interaction ^54,62,63^. In the Cas12a DNA cleavage process, the RuvC active site first cuts the non-target strand, then the target strand ^64^. While double-strand breaks are undesired for CRISPRi applications, we wondered whether favoring the nicked DNA intermediate might reduce the R-loop dissociation rate (Fig. 2A, see Discussion). In support of this possibility, in vitro studies showed that dCas12a:DNA complexes are 20-fold more stable when the non-target strand is pre-cleaved ^54^, and that non-target strand nicking biases Cas12a:DNA complexes away from dissociation-prone conformations^65,66^.

**Figure 2.**
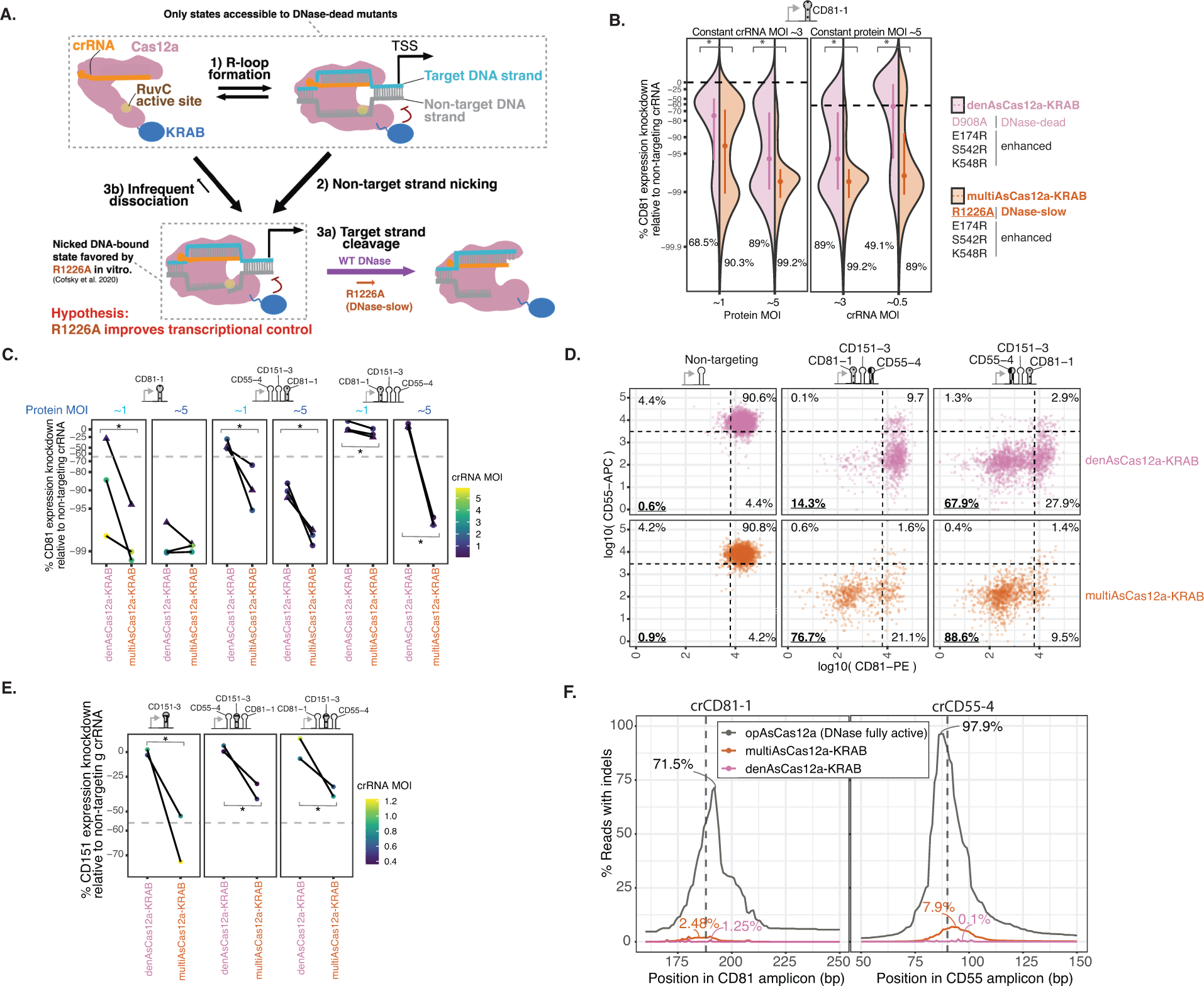
– multiAsCas12a-KRAB (R1226A/E174R/S542R/K548R), an engineered variant that favors a nicked DNA intermediate, substantially improves lentivirally delivered CRISPRi activity. **A)** Model of Cas12a DNA binding and cleavage states for wildtype DNase vs. R1226A mutant based on prior in vitro studies as detailed in main text. Sizes of arrows qualitatively reflect reaction rates. **B)** Comparison of denCas12a-KRAB (D908A/E174R/S542R/K548R) vs. multiAsCas12a-KRAB (R1226A/E174R/S542R/K548R) in CRISPRi knockdown of CD81 using crCD81-1. CD81 expression assayed by flow cytometry 10 days after crRNA transduction. Left panel: Holding crRNA MOI constant at ∼3 while testing protein MOI ∼1 vs. ∼5. Right panel: Holding protein MOI constant at ∼5 while testing crRNA MOI at ∼3 vs. ∼0.5. Asterisks indicate p *<*0.01 for one-sided Wilcoxon rank-sum test of single-cell distributions. Percentage of cells below the 5th percentile of non-targeting crRNA control (dashed line) is shown for each condition. One biological replicate is shown for each condition; additional replicates shown in Fig. S9. **C)** Comparison of CD81 knockdown by lentivirally delivered denAsCas12a-KRAB vs. multiAsCas12a-KRAB at protein MOI ∼1 vs. ∼5 across a panel of single and 3-plex crRNA constructs, while holding constant crRNA MOI for each paired fusion protein comparison for each crRNA construct. Dashed gray line indicates 5th percentile of non-targeting crRNA control. crRNA MOI indicated by color scale. Lines connect paired replicates. One-sided Wilcoxon rank-sum tests were performed on single-cell distributions for each replicate, and asterisk denotes p*<*0.01 for all paired replicates within each condition. Dots indicate flow cytometry measurement 10 days after crRNA transduction; triangles indicate flow cytometry measurement 16 days after crRNA transduction. **D)** Same as C but showing scatter plot of CD55-APC and CD81-PE antibody co-staining signals on flow cytometry performed 16 days after transduction of the indicated crRNA constructs in K562 cells lentivirally transduced with denAsCas12a-KRAB vs. multiAsCas12a-KRAB at protein MOI ∼5. Quadrants drawn based on the 5th percentile of non-targeting controls and the percentage of cells in each quadrant denoted. **E)** K562 cells piggyBac-engineered to constitutively express denAsCas12a-KRAB or multiAsCas12a-KRAB were transduced with the in-dicated crRNA constructs, followed by measurement of CD151 expression by antibody staining and flow cytometry 13 days after crRNA transduction. Median CD151 expression knockdown relative to non-targeting control is shown for each individual replicate. Dashed gray line indicates 5th percentile of non-targeting crRNA control. crRNA MOI indicated by color scale. One-sided Wilcoxon rank-sum test was performed for single-cell expression distributions comparing denAsCas12a-KRAB vs. multiAsCas12a-KRAB for each replicate; asterisk indicates p*<*0.001 for all replicate-level comparisons. **F)** Indel quantification from PCR amplicons surrounding target sites of crCD81-1 and crCD55-4 in cells lentivirally transduced at protein MOI ∼5 for denAsCas12a-KRAB and multiAsCas12a-KRAB. Cells lentivirally transduced with opAsCas12a (DNase fully active) are shown for comparison. Percent of reads containing indels at each base position within the amplicon is plotted, with labels indicating maximum indel

To engineer nicking-induced stabilization of AsCas12a binding to DNA for CRISPRi applications, we incorporated R1226A, a mutation that has not been tested in the context of transcriptional control. Relative to wild-type AsCas12a, the R1226A mutant, originally described as a nickase ^55^, is ∼100-1,000 fold slower in cleaving the non-target DNA strand and ∼10,000-fold slower in cleaving the target DNA strand in vitro ^54^. Consistent with nicking-induced stabilization, AsCas12a R1226A indeed binds DNA more strongly in vitro than the fully DNase-inactivated D908A variant ^54^. We expect the R1226A mutation to both disfavor R-loop reversal and slow progression to double-stranded breaks (Fig 2A; see Discussion). We hypothesized that, by trapping the ribonucleoprotein:DNA in a nicked DNA intermediate, the R1226A mutation would prolong chromatin occupancy and thus the time available for the KRAB domain to recruit transcriptional repressive complexes.

To test the impact of R1226A on CRISPRi activity, we replaced the DNase-inactivating D908A in denAsCas12a-KRAB with R1226A, and hereafter refer to this Cas12a variant as multiAsCas12a (<u>mul</u>tiplexed transcriptional interference, i.e. R1226A/E174R/S542R/K548R). To compare dose-sensitivity of their CRISPRi activities, we stably expressed denAsCas12a-KRAB and multiAsCas12a-KRAB protein and crRNA constructs by lentiviral transduction at high vs. low MOI’s for protein and crRNA constructs. Across a panel of single and 3-plex crRNA constructs, multiAsCas12a-KRAB consistently exhibits robust CRISPRi with less sensitivity to low MOI of protein or crRNA constructs (Fig. 2B-C, Fig. S9 and Fig. S10). Notably, multiAsCas12a-KRAB substantially rescues the activities of several crRNA constructs that are virtually inactive for denAsCas12a-KRAB even at high protein dose (Fig. 2C), including a non-canonical GTTC PAM target (crCD151-3, Fig. 2E). Targeting by multiAsCas12a-KRAB results in low indel frequencies at crCD81-1 (2.48%) and crCD55-4 (7.9%) target sites (Fig. 2F). Simulations accounting for DNA copy number indicate that any possible gene expression impact from these indels are far lower than the observed target gene knockdown (Fig. S14A-B). multiAsCas12a-KRAB CRISPRi activity shows generally minimal or no off-target transcriptomic effects as evaluated by bulk RNA-seq (Fig. S11).

### multiAsCas12a-KRAB enables multi-gene transcriptional repression using higher-order arrayed crRNA lentiviral constructs

We next tested performance of multiAsCas12a-KRAB in targeting >3 genomic sites per cell for CRISPRi using lentiviral crRNA arrays ^29^. To minimize the possibility of lentiviral recombination, the expression construct (Fig. 3A) uses a unique direct repeat variant at each position of the array, selected from a set of previously tested direct repeat variants^28^. We assembled a panel of 13 distinct crRNA constructs (7 single-plex, two 3-plex, two 4-plex, two 5-plex, and two 6-plex) from individually active TSS-targeting spacers (Fig. 3B-C and Fig. S12). For these 13 crRNA constructs, we compared the CRISPRi activities of denAsCas12a-KRAB, multiAsCas12a-KRAB, and multiAsCas12a (no KRAB). For a subset of crRNA constructs we also added enAsCas12a-KRAB (DNase fully active) as comparison. For these experiments and the remainder of this study we use piggyBac transposition to constitutively express all fusion proteins at very similar levels (Fig. S3 and Fig. S13B), which yields results similar to that obtained from high MOI (∼5) lentiviral fusion protein delivery and avoids day-to-day variations in lentiviral titers. Across the entire crRNA panel, multiAsCas12a-KRAB substantially outperforms denAsCas12a-KRAB in CRISPRi activity for 7 out of 7 constructs targeting CD81 (Fig. 3B); 4 out of 6 constructs targeting B2M (Fig. S12); and 6 out of 6 constructs targeting KIT (Fig. 3C). For CD55 (Fig. S12), multiCas12a-KRAB substantially outperforms denAsCas12a-KRAB using crCD55-5 (weaker spacer), and performs the same as or marginally better than denAsCas12a-KRAB for 7 constructs containing crCD55-4 (stronger spacer). Similarly superior CRISPRi performance by multiAsCas12a-KRAB was observed in C4-2B cells (Fig. S4).

**Figure 3.**
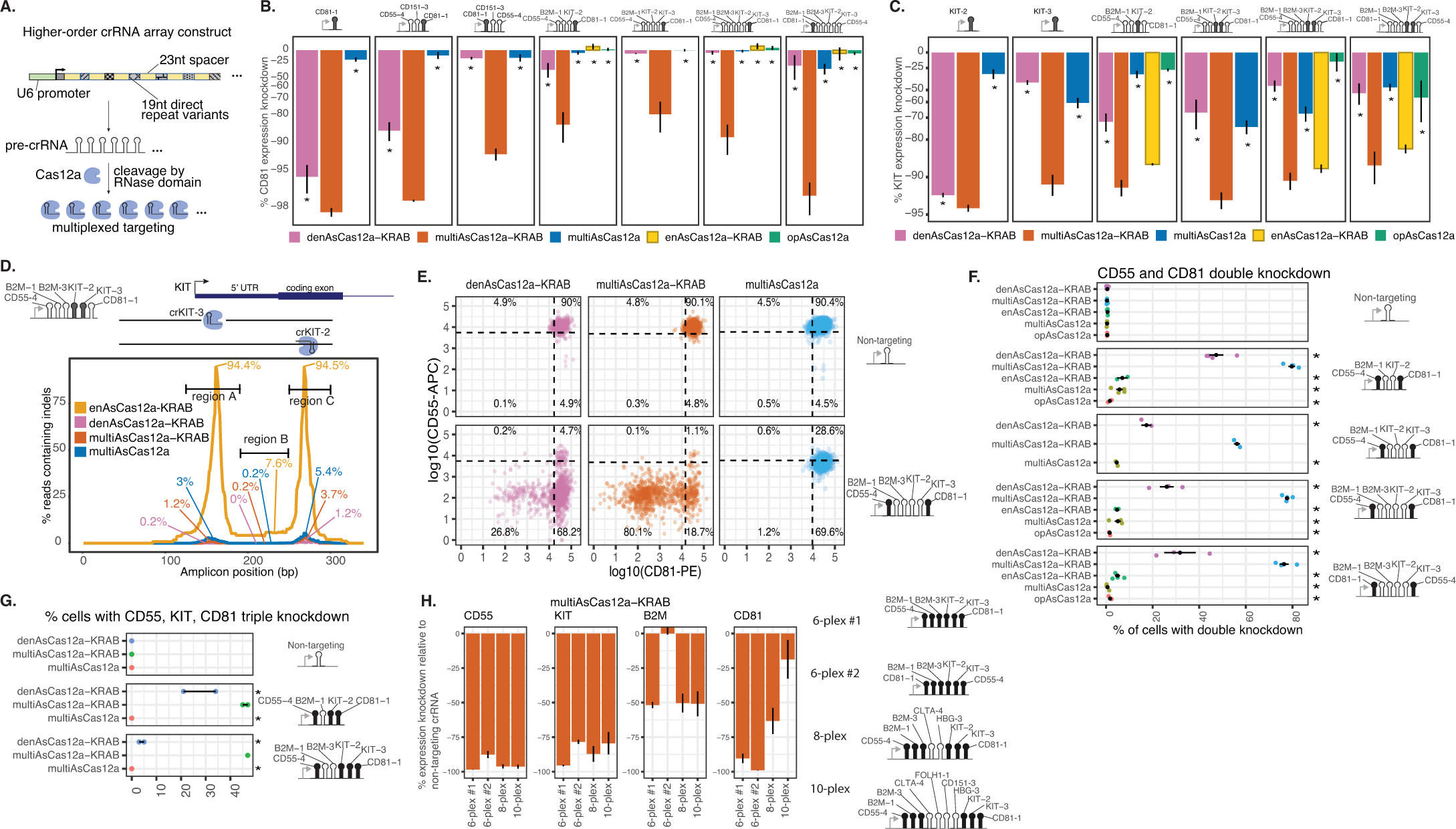
– multiAsCas12a-KRAB enables multi-gene CRISPRi perturbations using higher-order arrayed crRNA lentiviral constructs. **A)** Schematic for higher-order crRNA expression constructs. 23nt spacers are interspersed by 19nt direct repeat variants (DeWeirdt et al., 2020) uniquely assigned to each position within the array. **B)** Flow cytometry analysis of CD81 expression knockdown by antibody staining 6 days after transduction of the indicated lentiviral crRNA constructs in K562 cells engineered to constitutively express the specified fusion protein construct. Shown are averages of median single-cell expression knockdown from 2-5 biological replicates for each crRNA construct, with error bars indicating SEM. One-sided Wilcoxon rank-sum test was performed for differences in single-cell expression distributions for each fusion protein against multiAsCas12a-KRAB for each individual replicate. Asterisk indicates p *<* 0.01 for all replicates for a given pairwise comparison. **C)** Same as B, but shown for KIT expression knockdown. **D)** Indel quantification for the indicated fusion protein constructs using a 6-plex crRNA construct encoding crKIT-2 and crKIT-3 that target opposite strands at sites spaced 112 bp apart near the KIT TSS. Following crRNA transduction, cells were sorted on day 3 for GFP marker on the crRNA construct, and the 340bp genomic region surrounding both crRNA binding sites was PCR amplified from cell lysates harvested 15 days after crRNA transduction. The maximum percentages of reads containing indels overlapping any base position within each of the demarcated regions (region A, region B, region C) are shown. **E)** Comparison of the indicated fusion protein constructs in dual CD55 and CD81 CRISPRi knockdown 10 days after lentiviral transduction of a 6-plex crRNA construct by flow cytometry. Shown are log10 fluorescence intensity for each antibody stain and the percentages of cells in each quadrant, defined by the 5th percentile of non-targeting crRNA for each fluorescence signal, are indicated. **F)** Summary of the same experiment in E for a larger panel of crRNA constructs, showing the percentage of cells with successful double-knockdown of CD55 and CD81 (e.g. same gating strategy as bottom left quadrant in E). 2-6 biological replicates are shown as individual data points and summarized by the mean and SEM as error bars. Two-sample chi-square test was used compare the proportion of cells with double-knockdown between multiAsCas12a-KRAB and each of the other fusion protein constructs; asterisk indicates p*<*0.01 for all replicates. **G)** Analogous to F, except triple knockdown of CD55, KIT, and CD81 was quantified by the percentage of cells that are below the 5th percentile along all 3 dimensions on day 33 after transduction of crRNA constructs. 2 biological replicates are shown as individual data points. Two-sample chi-square test was used compare the proportion of cells with double-knockdown between multiAsCas12a-KRAB and each of the other fusion protein constructs; asterisk indicates p*<*0.01 for all replicates. **H)** Gene expression knockdown by multiAsCas12a-KRAB using 6-plex, 8-plex and 10-plex crRNA array constructs was measured by flow cytometry 10-11 days after lentiviral transduction of crRNA constructs. Shown are median gene expression knockdown averaged from 2-4 biological replicates, with error bars denoting SEM. CRISPRi activities of crFOLH1-1, crCD151-3 and crHBG-3 were not assayed in this experiment, but these spacers are active when encoded as individual crRNAs as shown in Fig. S4B, Fig. 2E, and Fig. 5A, respectively. All protein constructs shown in A-H were delivered by piggyBac transposition into K562 cells and sorted for the same expression level of the P2A-BFP marker (except opAsCas12a was delivered by lentiviral transduction and selected for by puromycin-resistance marker).

For all crRNA constructs tested, multiAsCas12a alone shows much lower impact on target gene expression than multiAsCas12a-KRAB (e.g. Fig. 3B for CD81), demonstrating strong dependence of gene knockdown on the KRAB domain. For some target genes, such as KIT, partial knockdown can be observed for multiAsCas12a alone (Fig. 3C). Such gene knockdown may be due to 1) non-genetic perturbation of transcription, or 2) alteration of DNA sequences crucial for transcription due to residual DNA cutting. To distinguish these possibilities, we used short-read Illumina sequencing of PCR amplicons (Fig. 3D, Fig. S14C and Fig. S15A), as well as long-read Nanopore sequencing of native genomic DNA up to tens of kilobases in length (Fig. S16C), to quantify indels generated by the panel of fusion proteins using single-site or dual-site targeting in the KIT locus. For multiAsCas12a-KRAB we observed a maximum indel frequency of 5% anywhere in the 340 bp PCR amplicon when simultaneously targeting two sites spaced ∼112 bp apart in the KIT locus (Fig. 3D). Based on this observed indel frequency, we calculated an upper estimate of expected 1.3% median KIT expression knockdown driven solely by indels, far less than the observed 90.4% median expression knockdown, which is 44.4% in excess of the observed for denAsCas12a-KRAB (Fig. S15B). Similar conclusions are supported by additional measurements at this and other loci obtained for multiAsCas12a-KRAB and/or multiAsCas12a using short-read PCR amplicon sequencing (Fig. S14) and/or long-read Nanopore sequencing (Fig. S16). Altogether, our analyses of indel frequencies demonstrate that target gene knockdown by multiAsCas12a-KRAB is largely attributable to non-genetic perturbation of transcription via a combination of direct obstruction of transcription by the Cas protein (as was observed for dCas9^43,44,67^) and KRAB-mediated repression.

At the single cell level, multiAsCas12a-KRAB consistently outperforms denAsCas12a-KRAB in the fraction of cells with successful double knockdown (Fig. 3E-F and Fig. S17) and triple knockdown (Fig. 3G and Fig. S17) of target genes using higher-order crRNA arrays. To test the upper limit of multiplexing, we constructed 8-plex and 10-plex constructs assembled using individually active spacers. In these 8-plex and 10-plex arrays, spacers encoded in various positions within the array maintain robust CRISPRi activity (i.e. for CD55, KIT and B2M, Fig 3H). However, for crCD81-1 encoded at the 3’ most position shows progressive diminishment in CRISPRi activity with further multiplexing at 8-plex and 10-plex (Fig. 3H). This pattern suggests an intrinsic deficiency of crCD81-1 that is unmasked by further multiplexing, perhaps related to the dose sensitivity of this spacer (Fig. 2B-C). Nevertheless, these results indicate that 8-plex and 10-plex crRNA arrays can support robust CRISPRi activity for most spacers within these arrays. We also observed that a specific 6-plex crRNA construct (crCD81-1_crB2M-1_crB2M-3_crKIT-2_crKIT-3_crCD55-4, 6-plex #2 in Fig. 3H) fails to knockdown B2M, despite robust CRISPRi of the other target genes. However, the same combination of spacers in a slightly different 6-plex arrangement (crCD55-4_crB2M-1_crB2M-3_crKIT-2_crKIT-3_crCD81-1) and also in 8-plex and 10-plex embodiments achieve ∼50% B2M knockdown (Fig. 3H). These results indicate the existence of still unpredictable pre-crRNA sequence context influences on CRISPRi activity of specific spacers, unrelated to genomic distance from the U6 promoter.

### multiAsCas12a-KRAB outperforms denCas12a-KRAB and performs similarly to dCas9-KRAB in pooled single-guide CRISPRi screens

Given the success of multiAsCas12a-KRAB in individual well-based assays using lentivirally delivered crRNAs, we next evaluated its performance in the context of high-throughput pooled screens. We designed a library, referred to as Library 1 (summarized in Fig. S18), aimed at extracting patterns for Cas12a CRISPRi activity with respect to genomic position relative to the TSS using cell fitness as a readout. Library 1 contains 77,387 single crRNA lentiviral constructs tiling all predicted canonical TTTV PAM sites and non-canonical PAM’s (recognizable by enAsCas12a ^31^) in the −50bp to +300bp region around the TSS’s of 559 common essential genes with K562 cell fitness defects in prior genome-wide dCas9-KRAB screens ^68^.

Using K562 cells piggyBac-engineered to constitutively express multiAsCas12a-KRAB or denAsCas12a-KRAB, we conducted a pooled cell fitness screen using this TSS tiling crRNA library (MOI = 0.15). In this assay, CRISPRi knockdown of target essential genes results in the relative depletion of cells harboring the corresponding crRNA over time, quantified as a cell fitness score (Fig. 4A). Concordance between cell fitness scores of screen replicates is high for multiAsCas12a-KRAB (R = 0.71) and much lower for denAsCas12a-KRAB (R = 0.32), the latter due to much lower signal-to-background ratio (Fig. S19). The cell fitness score distributions are virtually indistinguishable between the intergenic targeting negative controls and the non-targeting negative controls (Fig. S20), indicating no appreciable non-specific genotoxicity from multiAsCas12a-KRAB single-site targeting. Among the 3,326 crRNA’s targeting canonical TTTV PAM’s, 24.5% vs. 17.5% showed a fitness defect in multiAsCas12a-KRAB vs. denAsCas12a-KRAB, respectively (using the 5th percentile of intergenic negative controls as a threshold), with the magnitude of effect for each crRNA overall stronger for multiAsCas12a-KRAB (Fig. 4B).

**Figure 4.**
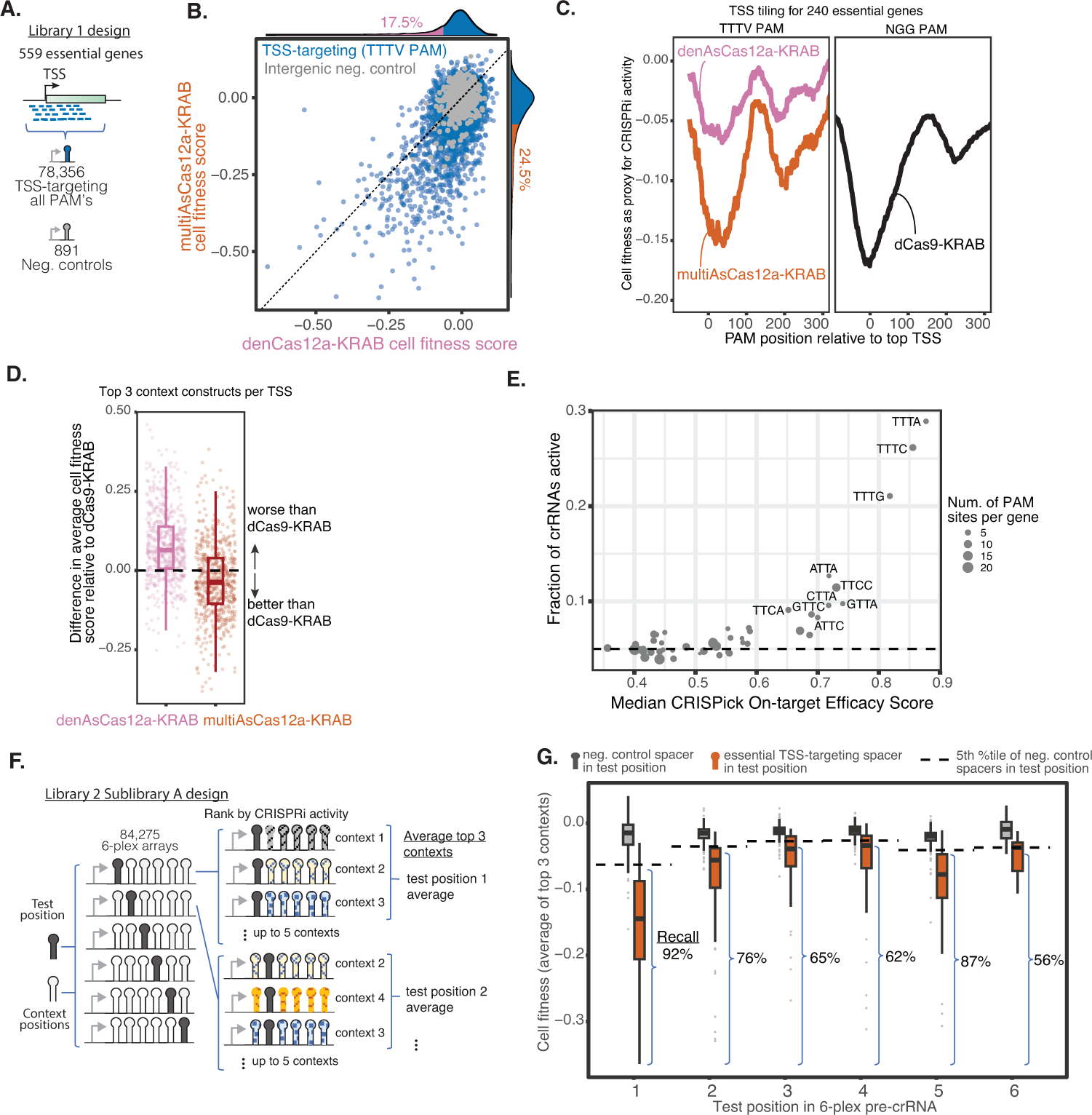
– multiAsCas12a-KRAB enables TSS-targeting pooled CRISPRi screens, including with 6-plex crRNA arrays. **A)** Design of Library 1 consisting of single crRNAs tiling TSS-proximal regions of essential genes. **B)** Library 1: Scatter plot of cell fitness scores in K562 cells for multiAsCas12a-KRAB vs. denCas12a-KRAB for 3,334 single crRNA constructs with sufficient read coverage for analysis and targeting canonical TTTV PAMs within −50bp to +300bp window of 584 essential gene TSS’s. Marginal histograms show percentage of crRNA constructs with cell fitness scores exceeding the 5th percentile of negative control crRNAs. y = x line is shown. **C)** Library 1: Moving average cell fitness score across all TTTV PAM-targeting crRNAs at each PAM position relative to the TSS (left), shown for the 240 essential gene TSS’s for which analogous dCas9-KRAB NGG PAM tiling screen data (Nuñez et al., 2021) is available in K562 cells (right). **D)** Library 1: Boxplots of average cell fitness scores of top 3 crRNAs for each essential TSS for multiAsCas12a-KRAB or denCas12a-KRAB, subtracted by the average cell fitness scores from top 3 sgRNAs for the same TSS for dCas9-KRAB (Nuñez et al., 2021). The top 3 sgRNA’s per TSS for dCas9-KRAB are taken from a prior genome-wide screen that used 10 sgRNAs per TSS, which were pre-selected based on bioinformatic prediction of strong sgRNA activity (Horlbeck et al. 2016a). Boxplots show median, interquartile range, whiskers indicating 1.5x interquartile range, and are overlaid with individual data points. **E)** Library 1: Bubble plot of the fraction of crRNAs for each PAM that exceed the 5th percentile of negative controls, versus the median CRISPick on-target efficacy score for that PAM. Size of the bubble indicates number of PAM sites per gene. **F)** Design of Library 2 Sublibrary A, aimed at evaluating CRISPRi activity each position in the 6-plex array. For each 6-plex array, a specific position is defined as the test position (which can encode either a TSS-targeting spacer or a negative control spacer), and the remaining positions are referred to as context positions encoding one of 5 sets of negative control spacers designated only for context positions. **G)** Library 2 Sublibrary A: Analysis of 2,987 6-plex constructs with sufficient read coverage and encodes in the test position one of 123 spacers that scored as strong hits as single crRNAs in the Library 1 screen, compared to constructs that encode an intergenic negative control spacer in the test position. Boxplots show cell fitness scores averaged from the top 3 context constructs of each test position spacer in the 6-plex array, grouped by negative control spacers vs. essential TSS-targeting spacers in each test position. Recall is calculated as the percentage of essential TSS-targeting spacers (that were empirically active in the single crRNA Library 1 screen) recovered by the Library 2 6-plex crRNA array screen for a given test position, using the 5th percentile of constructs containing negative control spacer in the same test position as a threshold for calling hits. Boxplots display median, interquartile range, whiskers indicating 1.5x interquartile range, and outliers. One-sided Wilcoxon rank-sum test was performed for the difference in the distributions of negative control spacers vs. TSS-targeting spacers at each position, with asterisks indicating p*<*0.0001.

Previous studies using dCas9-KRAB have identified a strong association between CRISPRi activity and genomic proximity of the crRNA binding site to the TSS ^44,52^. We found that multiAsCas12a-KRAB generates a metagene profile of average CRISPRi activity around the TSS that is remarkably similar in magnitude and bimodal genomic distribution of CRISPRi activity obtained by dCas9-KRAB targeting NGG PAM’s (Fig. 4C), consistent with nucleosomal hindrance near the +150bp region ^52,69^. denCas12a-KRAB is substantially weaker than both at all positions relative to the TSS (Fig. 4C). multiAsCas12a-KRAB also outperforms denAsCas12a-KRAB in the average CRISPRi activity of the top 3 best performing crRNAs/sgRNA for each TSS, and is similar to dCas9-KRAB (Fig. 4D). Compared to the canonical TTTV PAM’s, a smaller proportion of crRNAs targeting non-canonical PAM’s are active, in agreement with lower median CRISPick on-target activity predictions (Fig. 4E) ^28,70^. Within each PAM sequence, individual crRNAs show significant variations in activity not accounted for by CRISPick predictions (Fig. S21).

### multiAsCas12a-KRAB enables pooled sequencing screens using 6-plex crRNA arrays

To evaluate the performance of multiAsCas12a-KRAB in pooled sequencing screens using multiplexed crRNA constructs, we constructed a library consisting of 6-plex crRNAs. We refer to this 6-plex library as Library 2 (summarized in Fig. S18), which includes Sublibrary A (described in this section) and Sublibrary B (described in the next section). Sublibrary A was designed to contain 84,275 6-plex constructs for evaluating CRISPRi activity at each of the 6 positions in the array in a K562 cell fitness screen (Fig. 4F). Each 6-plex construct has one of the 6 positions designated as the “test” position, which can encode either 1) a spacer targeting one of the top 50 essential gene TSS’s (ranked based on prior dCas9-KRAB screen data ^52^), or 2) an intergenic negative control (Fig. 4F). The remaining 5 positions in the array are designated as “context” positions that encode negative control spacers drawn from a separate set of 30 negative control spacers (Fig. 4F). The motivation for this library design was to enable sampling multiple sets of context spacers for a given test position.

The entirety of Library 2 was used in a cell fitness screen in K562 cells piggyBac-engineered to stably express multiAsCas12a-KRAB. For a given test position spacer, we calculated the average cell fitness scores from the top 3 context constructs ranked by cell fitness score (Fig. 4F). The 123 spacers that previously showed strong cell fitness scores as single crRNAs in the Library 1 screen also showed cell fitness score distributions (average of top 3 contexts) that are clearly distinct from the corresponding negative control distributions for the same test position (Fig. 4G). Each TSS-targeting spacer encoded in the test position shows weaker cell fitness score than the same spacer encoded singly in the Library 1 screen (Fig. S22). The recall of empirically active single crRNA spacers from the Library 1 screen by the 6-plex crRNA constructs in this Library 2 Sublibrary A screen ranges from 56%-92% across test positions (Fig. 4G). As each position in the array is assigned a unique direct repeat variant held constant across all constructs in this analysis, these apparent positional biases may reflect contributions from differences among direct repeat variants. Together, these results systematically demonstrate that the majority of individually active crRNAs retain measurable activities when embedded within 6-plex crRNA arrays in the setting of pooled screens using multiAsCas12a-KRAB.

### multiAsCas12a-KRAB enables discovery and higher-order combinatorial perturbations of cis-regulatory elements

The human genome contains ∼500,000 predicted enhancers ^71^, a small minority of which have been functionally tested by perturbations. To our knowledge, no study has reported enhancer perturbation by CRISPRi using Cas12a. We confirmed that, similar to dCas9-KRAB ^72^, multiAsCas12a-KRAB targeting using single crRNAs can effectively perturb a known enhancer of the HBG gene, HS2 (Fig. 5A and Fig. S11). We next used multiAsCas12a-KRAB to discover previously uncharacterized enhancers using the CD55 locus in K562 cells as a myeloid cell model. CD55 encodes for decay-accelerating factor, a cell surface protein that inhibits the activation of complement and is expressed in most human cell types ^73^. CD55 function in the myeloid lineage is relevant in multiple disease states, including paroxysmal nocturnal hemoglobinuria ^74^ and malaria ^75,76^. To our knowledge, no known enhancers in myeloid cells have been identified for CD55. In K562 cells, several DNase hypersensitive sites (DHS) bearing histone 3 lysine 27 (H3K27Ac), a modification associated with active enhancers ^71,77^, reside near CD55 (Fig. 5B). To conduct a well-based flow cytometry screen of the DHSs within this general region for enhancers that regulate CD55, we designed 21 4-plex crRNAs (encompassing 88 unique spacers) targeting 11 manually selected regions (R1-R11) bearing varying levels of DNase hypersensitivity and H3K27Ac, plus a negative control region (R12) devoid of DHS and H3K27Ac. R1-R4 are predicted by the activity-by-contact (ABC) model ^78^ as candidate enhancers. Each region is independently targeted by two completely distinct 4-plex crRNAs (except R10 and R12 are each targeted by one 4-plex crRNA). We observed ∼50%-75% reduction in CD55 expression upon multiAsCas12a CRISPRi targeting of the ABC-predicted R1-R4, whereas no decrease in CD55 expression is observed for R5-R12. For each region, the two distinct 4-plex crRNA arrays show quantitatively similar levels of CD55 knockdown (Fig. 5B), indicating each array contains some 4-plex or lower-order combination of active spacers. This consistency in the magnitude of CD55 expression knockdown likely reflects the magnitude of true enhancer impact on gene transcription, rather than technical peculiarities of individual spacer activities, which might be more unpredictably variable and labor-intensive to test if encoded as single-plex perturbations. To our knowledge, R1-R4 are the first functionally demonstrated enhancers for CD55 in a myeloid cell type, in addition to another enhancer recently reported in a B-cell model ^79^. Using opAsCas12a to target R1-R4 for DNA cutting using the same 4-plex crRNAs elicits very little or no CD55 expression knockdown, despite potent knockdown by a positive control crRNA targeting a coding exon (Fig. 5C).

**Figure 5.**
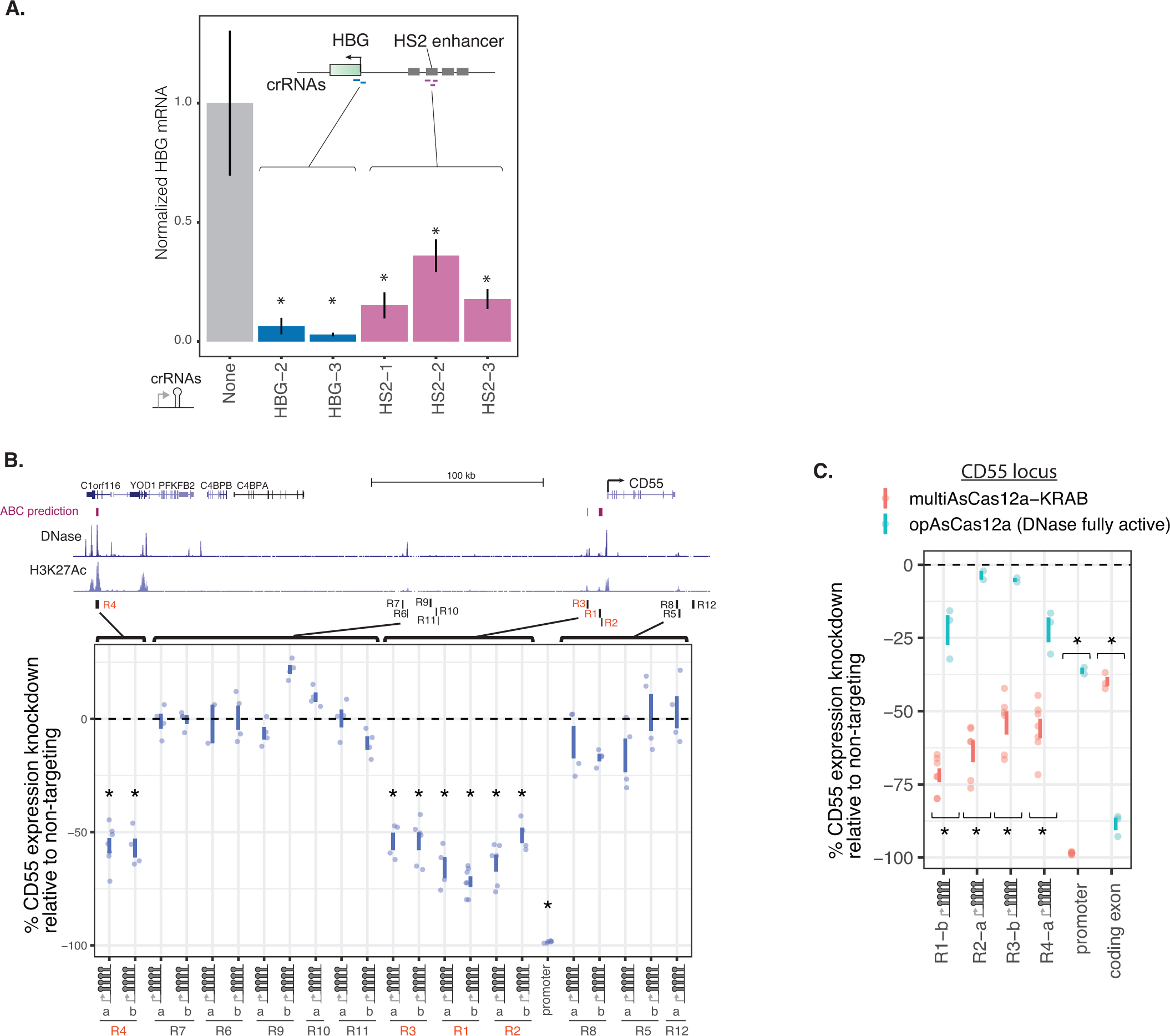
– multiAsCas12a-KRAB CRISPRi enables enhancer perturbation and discovery. **A)** K562 cells constitutively expressing multiAsCas12a-KRAB are lentivirally transduced with single crRNAs targeting the HBG TSS or its known enhancer, HS2. Shown are HBG mRNA levels measured by RT-qPCR, normalized to GAPDH levels and averaged across 6-7 technical replicates from 2 independent experiments. Error bars denote SEM. One-sided unpaired Student’s t-test was performed to compare untransduced control vs. each individual crRNA; asterisk denotes p*<*0.05. **B)** Genome browser view of the CD55 locus, including predicted enhancers using the activity-by-contact model and DNase-seq and H3K27Ac ChIP-seq tracks from ENCODE. K562 cells piggyBac-engineered to constitutively express multiAsCas12a-KRAB was transduced with 4-plex crRNA constructs targeting regions (R1-R11) in the CD55 locus, and R12 as a negative control region devoid of enhancer features. Each unique 4-plex crRNA construct is labeled as “a” or “b”. For comparison, targeting the CD55 promoter using a 6-plex crRNA array (crCD55-4crB2M-1crKIT-2crKIT-3crCD81-1) is included. CD55 expression was assayed by flow cytometry between 9 and 11 days after crRNA transduction. One-sided Wilcoxon rank-sum test was performed on the median expression knockdown across 4-7 biological replicates for each crRNA construct, compared to the median expression knockdown of R12 (negative control region); asterisk indicates p *<* 0.01. **C)** Comparison of CRISPRi targeting in K562 cells engineered to constitutively express multiAsCas12a-KRAB vs. opAsCas12a using a subset of lentivirally transduced crRNA constructs from B, plus a crRNA construct targeting a coding exon of CD55 as a positive control for knockdown by DNA cutting. CD55 expression was assayed by flow cytometry 11 days after crRNA transduction. One-sided Wilcoxon rank-sum test was performed to compare the median expression knockdown of multiAsCas12a-KRAB vs. opAsCas12a across 2-7 biological replicates; asterisk indicates p*≤*0.05.

To further test the utility of multiAsCas12a-KRAB in studies of enhancer function, we used the MYC locus as a model. Prior studies using CRISPRi pooled screens in K562 cells have shown that MYC expression is proportional to cell fitness ^80^ and is regulated by several enhancers identified by screens using cell fitness^80^ and mRNA expression ^81^ readouts. A recent study found that pairwise dCas9-KRAB perturbations of these enhancers elicit stronger phenotypes than perturbing single enhancers ^82^. Here we use multiasCas12a-KRAB to dissect the phenotypic impact of ≥3-plex perturbations of MYC cis-regulatory elements, which remained unknown. To avoid testing intractably numerous higher-order combinations of crRNA spacers that are largely uninformative due to the inclusion of weak or inactive spacers, we pre-screened for a small group of active 3-plex crRNA combinations that can be subsequently assembled into higher-order combinations. We used multiAsCas12a-KRAB to test four 3-plex crRNA constructs targeting combinations of MYC cis-regulatory elements (3 crRNAs for promoter and 3 crRNAs for each of 3 known enhancers, e1, e2 and e3) in a well-based cell competition assay (Fig. 6A-B). We found that these four 3-plex crRNAs induce varying degrees of cell fitness defect as a proxy of MYC expression knockdown, indicating that each construct contains some spacer combination with CRISPRi activity. For comparison, we included denAsCas12a-KRAB, multiAsCas12a, enAsCas12a-KRAB and opAsCas12a, which showed relative activities that further demonstrate multiAsCas12a-KRAB’s superior gene knockdown is largely attributable to non-genetic transcriptional perturbation (Fig. 6B).

**Figure 6.**
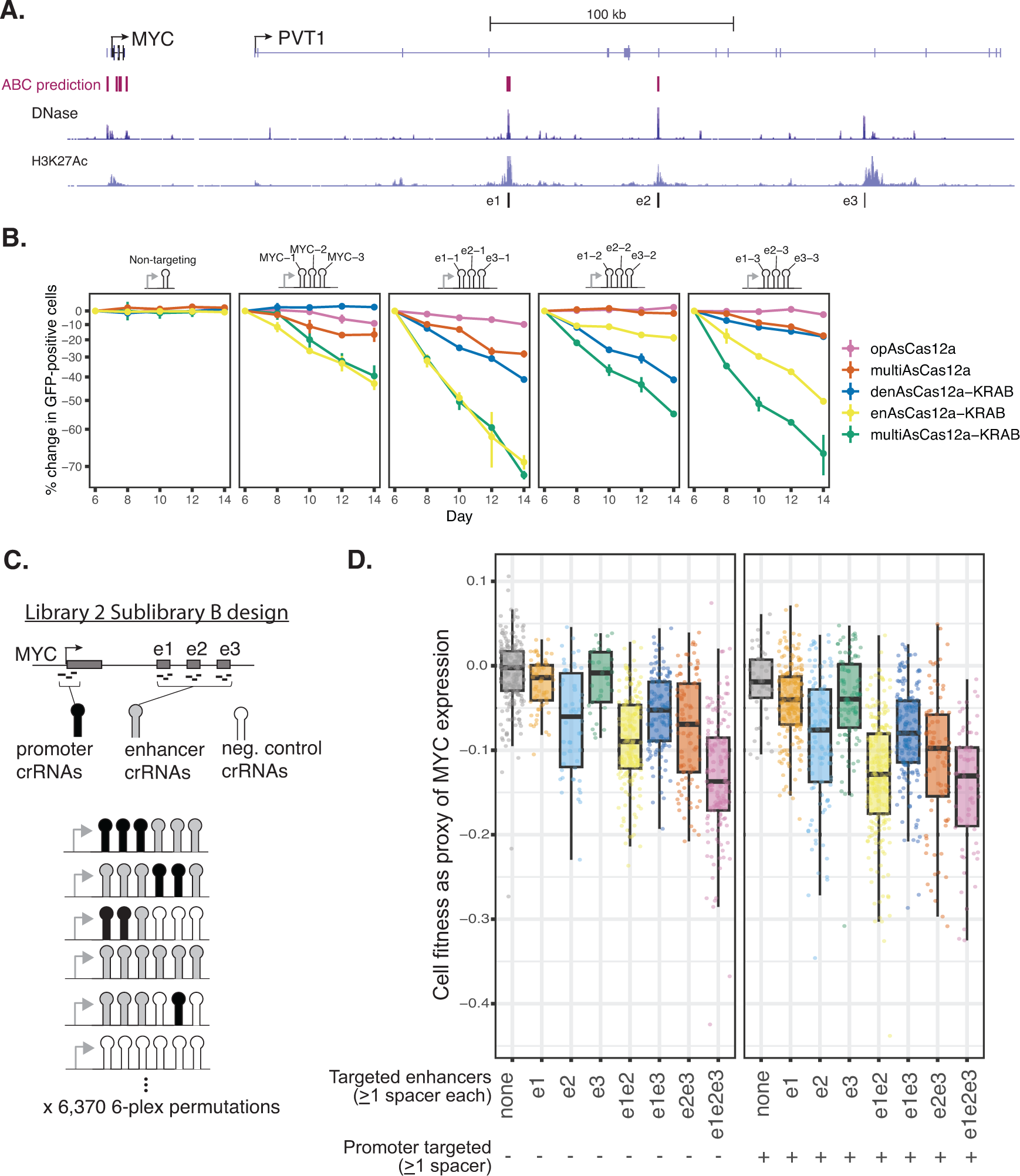
– multiAsCas12a-KRAB enables combinatorial targeting of cis-regulatory elements in pooled CRISPRi screens. **A)** Genome browser view of the MYC locus, including activity-by-contact model predictions, and DNase-seq and H3K27Ac ChIP-seq tracks from ENCODE. 3 of the known MYC enhancers (e1, e2, e3) in the body of the non-coding RNA, PVT1, are shown. **B)** K562 cells piggyBac-engineered to constitutively express the indicated panel of fusion protein constructs were transduced with one of 4 3-plex crRNA constructs targeting the MYC promoter or co-targeting the 3 enhancers using one crRNA per enhancer. Cell fitness as a proxy of MYC expression is measured as log2 fold-change in percentage of cells expressing GFP marker on the crRNA construct, relative to day 6 after crRNA transduction. Shown are the average of 2 biological replicates, with error bars denoting the range. **C)** 6,370 6-plex permutations of the 12 individual spacers from B, together with 3 intergenic negative control spacers, were designed and cloned as 6-plex crRNA arrays used in the design of Library 2 Sublibrary B. **D)** Library 2 Sublibrary B: Analysis of 1,823 constructs with sufficient read coverage, categorized based on whether each contains at least one of 3 crRNAs that target the MYC promoter, and/or at least one crRNA that targets each of the MYC enhancers. Boxplots summarize cell fitness score distributions (as proxy of MYC expression) of all constructs that fall in each category. Boxplots show median, interquartile range, whiskers indicating 1.5x interquartile range, and are overlaid with individual data points each representing a 6-plex construct.

We in silico assembled these 12 nominated spacers and 3 intergenic negative control spacers into Library 2 Sublibrary B, consisting of 6,370 6-plex permutations encoded as 6-plex crRNA arrays (Fig. 6C and summarized in Fig. S18). These 6-plex arrays each target up to 4 cis-regulatory elements (promoter + 3 enhancers) with up to 3 spacers per element. Negative control spacers fill in the remaining array positions not filled by targeting spacers. This Sublibrary B was included as part of the cell fitness screen for the entirety of Library 2. Among 1,823 6-plex arrays with sufficient read coverage for analysis, we grouped them into 16 categories, based on whether it encodes at least 1 spacer targeting the promoter, and/or at least 1 spacer targeting each of the 3 enhancers (Fig. 6D). We found that higher-order targeting of enhancers shows stronger cell fitness defects beyond lower-order enhancer combinations (Fig. 6D, left panel). Co-targeting the promoter together with any combination of enhancers showed greater cell fitness defect over targeting the promoter alone while also exhibiting the cumulative effects of multi-enhancer targeting (Fig. 6D, right panel). These results suggest that when targeting subsets of cis-regulatory elements in a locus by CRISPRi, other cis-regulatory elements can compete with CRISPRi to partially sustain gene transcription. Such effects may reflect how cis-regulatory elements combinatorially respond to endogenous repressive cues in the natural regulation of MYC gene transcription. These results indicate that co-targeting multiple cis-regulatory elements may be more effective for gene knockdown than targeting the promoter alone for some genes.

## Discussion

In this study, we engineered multiAsCas12a-KRAB as a new platform for higher-order combinatorial CRISPRi perturbations of gene transcription and enhancer function. The enhanced CRISPRi potency of multiAsCas12a-KRAB is more robust to lower doses of ribonucleoprotein (Fig. 2B-C, Fig. 2E), critically enabling higher-order multiplexing (Fig. 3) and high-throughput pooled screening applications conducted at single-copy integrations of crRNA (Fig. 4 and Fig. 6C-D). We propose that the improved CRISPRi activity of multiAsCas12a-KRAB emerges from prolonged chromatin occupancy due to DNA nicking (Fig. 2A). This strategy is conceptually distinct from prior protein engineering approaches to improving Cas12a function in mammalian cells, which focused on substituting for positively charged amino acid residues near the protein:DNA interface ^31,50^, using directed evolution to optimize DNA cleavage ^60,83^, or optimizing transcriptional effector domain function ^57^. We propose the following biophysical explanation for improved multiAsCas12a function, grounded in prior in vitro literature. In the absence of nicking, R-loop reversal occurs by invasion of the non-target strand into the crRNA:target strand duplex, displacing the crRNA in a process analogous to toehold-mediated nucleic acid strand displacement ^84^. Severing the non-target strand increases its conformational entropy and effectively destroys the toehold, decreasing the rate at which the non-target strand can invade the crRNA:target strand duplex ^84^. This model can also explain previous observations of cutting-dependent complex stabilization ^54,62,63^ and suggests that engineering a nicking preference may improve the efficacy of other Cas enzymes in chromatin targeting. Other potential explanations for multiAsCas12a’s enhanced CRISPRi activity include protein:DNA contacts formed after non-target-strand nicking (Naqvi et al., 2022) and/or nicking-induced relaxation of DNA supercoiling ^85^. We have demonstrated that the effects of multiAsCas12a-KRAB’s residual DNase activity on DNA sequence contributes minimally to target gene knockdown for typical functional genomics experimental conditions (Fig. 2F, Fig. 3B-G, Fig. 5C, Fig. 6B, Fig. S12, and Fig. S14-S16). Nevertheless, for screens involving strong positive selection, it may be possible for infrequent deletions to exert more appreciable influence on screen outcome.

We propose that multiAsCas12a-KRAB provides new solutions to a major challenge in combinatorial genetics: the infeasibility of surveying potentially enormous combinatorial spaces of ≥3-plex genetic perturbations. Testing a single higher-order N-plex combination also indirectly tests all or many of its constituent lower-order combinations, for up to a total of 2^N^ combinations. Thus, increases in multiplexing capability potentially yield exponential increases in search efficiency using group testing ^86,87^. In group testing (Fig. 7), a primary screen is conducted on grouped subjects (e.g. a multiplexed array of crRNA constructs) to reduce the costs otherwise incurred by individually testing all subjects (e.g. an individual crRNA). Our screen for CD55 enhancers instantiates this approach by testing 22 4-plex crRNA arrays targeting 12 candidate regions, indirectly testing 22 x 2^4^ = 352 crRNA combinations in a cost-effective well-based experiment (Fig. 5B). For this experimental objective, the grouped hits can be biologically interpreted without further testing (Fig. 7). For other objectives, such as the analysis of combinatorial cis-regulation at the MYC locus (Fig. 6), grouped hits can be followed by a focused secondary screen as needed (Fig. 7). For pooled sequencing screens, the ability to deterministically encode specific higher-order crRNA combinations in a single array is crucial for group testing. In contrast, cloning combinatorial guide libraries by a multiplicative and stochastic approach ^17^ requires testing all combinations at the onset, and thus is incompatible with group testing. Group testing can significantly compress the size of crRNA libraries to facilitate screens limited by assayable cell numbers. Group testing may be combined with compressed sensing ^88,89^ to facilitate screens with multidimensional phenotypic readouts ^19,20,59,90–95^.

**Figure 7.**
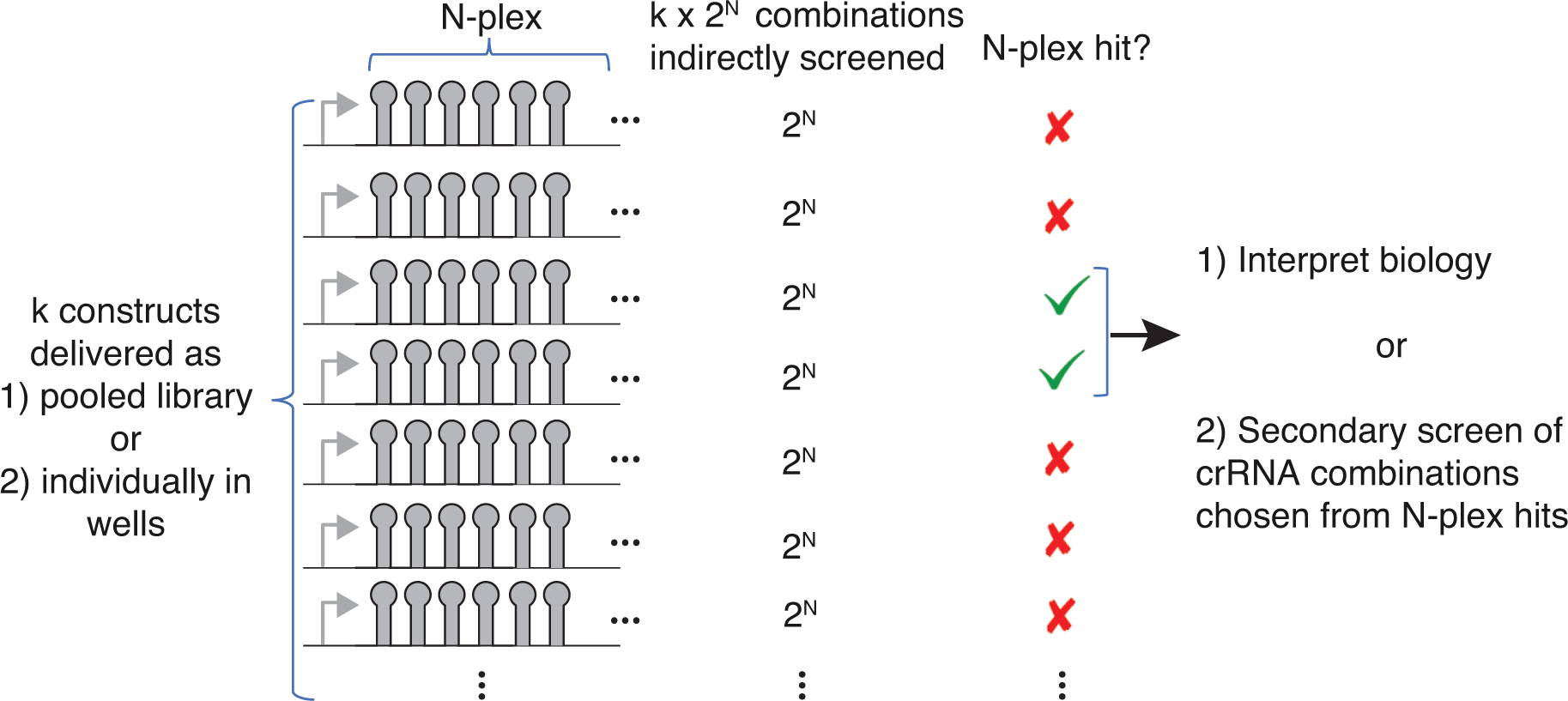
– Group testing framework for efficient exploration of combinatorial CRISPR perturbations.

A key parameter in group testing is the extent of potential signal dilution and/or interference relative to individual testing. Signal dilution can arise from limiting doses of ribonucleoprotein due to delivery format or reduction in the protein available to bind each individual crRNA due to increased crRNA multiplexing. Signal interference can arise if certain sequence features in one part of the crRNA array masks the activity of individual constituent crRNAs in a dominant fashion. Despite some evidence of signal dilution and/or interference for multiplexed crRNA arrays (Fig. 3H, Fig. 4G, and Fig. S22), multiAsCas12a-KRAB demonstrates sufficient robustness for yielding new biological insights into combinatorial cis-regulation at the MYC locus using 6-plex crRNAs in high-throughput pooled screens (Fig. 6C-D). While we have focused our optimizations to meet the stringent requirements of pooled screening formats, multiAsCas12a also significantly lowers technical barriers to higher-order combinatorial perturbations in array-based screening (Fig. 5B), which has recently improved in throughput ^96^. The assay format will likely influence the deliverable dose of synthetic components and thus the upper limit of multiplexing for effective CRISPRi using multiAsCas12a-KRAB. Improved prediction of crRNA array activities will likely further support highly multiplexed and/or dose-limited applications, thus extending the scalability of combinatorial genetic screens by group testing. A fully active 10-plex crRNA array could indirectly screen up to 2^10^ = 1,024 crRNA combinations. Another area for improvement is the observation that a low proportion of crRNAs targeting non-canonical PAMs show CRISPRi activity when targeted by multiAsCas12a-KRAB (Fig. 4E). Increasing the fraction of active crRNAs, especially for non-canonical PAM’s, would enable more reliable targeting with fewer crRNAs, especially in GC-rich TSS-proximal regions.

While we have focused on CRISPRi applications using the KRAB domain, the discovery and engineering of effector domains for chromatin perturbations by CRISPR-Cas is rapidly evolving. Recent advances include new repressive effectors ^97–100^, activation effectors ^57,98,100,101^, and combination effectors for epigenetic memory ^52,102–105^. We expect that multiAsCas12a can be flexibly combined with these and other effector domains to support group testing for many chromatin perturbation objectives. We envision multiAsCas12a and the group testing framework will enable elucidating and engineering combinatorial genetic processes underlying broad areas of biology at previously intractable scales.

## Author Contributions

C.C.H. and L.A.G. conceived of and led the design of the overall study. C.C.H. wrote the paper incorporating input from L.A.G., C.M.W., J.C.C., J.S., and agreement from all authors. C.C.H. conceived of testing the R1226A mutation for CRISPRi applications, proposed the conceptual link to group testing, and led the design, execution, and computational analysis of all experiments, with contributions from others noted below. L.A.G. proposed the dose of lentivirally delivered CRISPR components as a key parameter for evaluation, interpreted results, and guided and obtained funding for the overall study. C.M.W. designed, executed, and analyzed quantification of indel frequencies by short-read sequencing, and contributed to cloning and flow cytometry-based CRISPRi experiments. N.A.S. contributed to the execution of flow cytometry-based CRISPRi experiments, including all experiments in C4-2B cells, testing of higher-order crRNAs arrays, replicates of dose-response CRISPRi experiments, and enhancer CRISPRi experiments at the CD55 and MYC loci. R.D. contributed to cell line engineering and CRISPRi analysis of the DNase-dead mutant panel. Q.C. designed, executed, and analyzed all experiments testing truncated crRNAs, with guidance and funding obtained by J.S.. J.C.C. provided feedback on interpretations of the biophysics literature and proposed the entropic barrier to R-loop reversal upon severance of the non-target strand. S.M. contributed to crRNA cloning and cell culture. T.O. prepared 3’ RNA-seq Illumina sequencing libraries. A.A. contributed to screen data processing and analysis for 3’ RNA-seq. N.T. wrote scripts in Rust for screen read mapping and counting.

## Acknowledgements

We thank Brian Yu and Khushali Patel for assistance in Illumina and Nanopore sequencing; Michael Herschl for sharing a modified target-enrichment Nanopore sequencing protocol and Vince Tran for helpful discussions; Eric Chow for suggestions on Illumina sequencing of screen libraries; April Pawluk for feedback on manuscript; Gavin Knott, Ashir Borah, Garrett Wong, James Nunez, Jonathan Weissman, Howard Chang, Patrick Hsu, Silvana Konermann, and the Gilbert Lab for helpful discussions.

## Funding

C.C.H. is supported by the Physician Scientist Incubator, Dept. of Pathology, Stanford University School of Medicine and NIH NHGRI (K01HG012789). Q.C. and J.S. were supported by the Mark Foundation for Cancer Research. J.C.C. is a fellow of the Helen Hay Whitney Foundation. This work was funded in part by NIH (R01HG012227) and in part by GSK through an award to the UCSF Laboratory for Genomics Research to L.A.G.. L.A.G. is funded by the Arc Institute, NIH (DP2CA239597, UM1HG012660), CRUK/NIH (OT2CA278665 and CGCATF-2021/100006), a Pew-Stewart Scholars for Cancer Research award and the Goldberg-Benioff Endowed Professorship in Prostate Cancer Translational Biology. Sequencing of CRISPR screens and 3’ RNA-seq libraries was performed at the UCSF CAT, supported by UCSF PBBR, RRP IMIA, and NIH 1S10OD028511-01 grants.

## Declaration of Interests

C.C.H., C.M.W., R.D. and L.A.G. have filed patent applications related to multiAsCas12a. J.S. is a scientific consultant for Treeline Biosciences. L.A.G has filed patents on CRISPRoff/on, CRISPR functional genomics and is a co-founder of Chroma Medicine.

## Methods

### Plasmid Design and Construction

A detailed table of constructs generated in this study will be provided as a Supplemental File with all sequences. Constructs will be made available on Addgene. Cloning was performed by Gibson Assembly of PCR amplified or commercially synthesized gene fragments (from Integrated DNA Technologies or Twist Bioscience) using NEBuilder Hifi Master Mix (NEB Cat# E262), and final plasmids sequence-verified by Sanger sequencing of the open reading frame and/or commercial whole-plasmid sequencing service provided by Primordium.

### Protein constructs components

Fusion protein sequence maps are provided in Supplemental Files. To summarize, denAsCas12a-KRAB, multiAsCas12a-KRAB, multiAsCas12a, and enAsCas12a-KRAB open reading frames were embedded in the same fusion protein architecture consisting of an N-terminal 6x Myc-NLS (Gier et al. 2020) and C-terminal XTEN80-KRAB-P2A-BFP (Replogle et al. 2022). The denAsCas12a open reading frame was PCR amplified from pCAG-denAsCas12a(E174R/S542R/K548R/D908A)-NLS(nuc)-3xHA-VPR (RTW776) (Addgene plasmid # 107943, from ^31^). AsCas12a variants described were generated by using the denAsCas12a open reading frame as starting template and introducing the specific mutations encoded in overhangs on PCR primers that serve as junctions of Gibson assembly reactions. opAsCas12a^29^ is available as Addgene plasmid # 149723, pRG232. 6xMyc-NLS was PCR amplified from pRG232. KRAB domain sequence from KOX1 was previously reported in ^43^. The lentiviral backbone for expressing Cas12a fusion protein constructs are expressed from an SFFV promoter adjacent to UCOE and is a gift from Marco Jost and Jonathan Weissman, derived from a plasmid available as Addgene 188765. XTEN80 linker sequence was taken from ^52^ and was originally from ^106^. For constructs used in piggyBac transposition, the open reading frame was cloned into a piggyBac vector backbone (Addgene #133568) and expressed from a CAG promoter. Super PiggyBac Transposase (PB210PA-1) was purchased from System Biosciences.

dAsCas12a-KRABx3 open reading frame sequence is from ^27^, in that study encoded within a construct referred to as SiT-ddCas12a-[Repr]. We generated SiT-ddCas12a-[Repr] by introducing the DNase-inactivating E993A by PCR-based mutagenesis using SiT-Cas12a-[Repr] (Addgene #133568) as template. Using Gibson Assembly of PCR products, we inserted the resulting ddCas12a-[Repr] open reading frame in-frame with P2A-BFP in a piggyBac vector (Addgene #133568) to enable direct comparison with other fusion protein constructs cloned in the same vector backbone (crRNA’s are encoded on separate plasmids as described below).

Fusion protein constructs described in Fig. S8 were assembled by subcloning the protein-coding sequences of AsCas12a and KRAB into a lentiviral expression vector using the In-Fusion HD Cloning system (TBUSA). AsCas12a mutants were cloned by mutagenesis PCR on the complete wildtype AsCas12a vector to generate the final lentiviral expression vector.

### crRNA expression constructs

Unless otherwise specified, individual single and 3-plex crRNA constructs were cloned into the human U6 promoter-driven expression vector pRG212 (Addgene 149722, originally from ^29^). Library1, Library2, some 3-plex and all 4-plex, 5-plex, and 6-plex *As.* crRNA constructs were cloned into pCH67, which is derived from pRG212 by replacing the 3’ DR with the variant DR8 ^28^. For constructs cloned into pCH67, the specific *As.* DR variants were assigned to each position of the array as follows, in 5’ to 3’ order:

3-plex: WT DR, DR1, DR3, DR8
4-plex: WT DR, DR1, DR10, DR3, DR8
5-plex: WT DR, DR1, DR16, DR10, DR3, DR8
6-plex: WT DR, DR1, DR16, DR18, DR10, DR3, DR8
8-plex: WT DR, DR1, DR16, DR_NS1, DR17, DR18, DR10, DR3, DR8
10-plex: WT DR, DR1, DR16, DR_NS1, DR4, DR_NS2, DR17, DR18, DR10, DR3, DR8

Where the sequences of DR_NS1 and 2 were based on combining hits from the variant DR screen from DeWeirdt et al., 2020. The sequences are DR_NS1: aattcctcctcttggaggt, and DR_NS2: aattcctcctataggaggt.

1-plex,3-plex, 8-plex, and 10-plex crRNA constructs were cloned by annealing complementary oligos, phosphorylation by T4 polynucleotide kinase (NEB M0201S), and ligated with T4 DNA ligase (NEB M0202) into BsmbI site of vector backbones. 4-plex, 5-plex and 6-plex crRNA arrays were ordered as double-stranded gene fragments and cloned into the BsmbI site of vector backbones by Gibson Assembly.

### Design of individual crRNAs

For cloning individual crRNA constructs targeting TSS’s, CRISPick was used in the enAsCas12a CRISPRi mode to design spacers targeting PAM’s located within −50bp to +300bp region around the targeted TSS. We manually selected spacers from the CRISPick output by picking TTTV PAM-targeting spacers (except for crCD151-3, which targets a non-canonical GTTC PAM) with the highest On-Target Efficacy Scores and generally excluded any spacers with high off-target predictions. The same non-targeting spacer was used throughout the individual well-based experiments and was randomly generated and checked for absence of alignment to the human genome by BLAT ^107^. The hg19 genomic coordinates for MYC enhancers are: e1 chr8:128910869-128911521, e2 chr8:128972341-128973219, and e3 chr8:129057272-129057795. DNA sequences from those regions were downloaded from the UCSC Genome Browser and submitted to CRISPick. The top 3 spacers targeting TTTV PAM’s for each enhancer were picked based on CRISPick On-target Efficacy Score, having no Tier I or Tier II Bin I predicted off-target sites, and proximity to the zenith of the ENCODE DNase hypersensitivity signal in K562 cells.

### Cell culture, lentiviral production, lentiviral transduction

All cell lines were cultured at 37deg. C with 5% CO2 in tissue culture incubators. K562 and C4-2B cells were maintained in RPMI-1640 (Gibco cat# 22400121) containing 25 mM HEPES, 2mM L-glutamine, and supplemented with 10% FBS (VWR), 100 units/mL streptomycin, and 100 mg/mL penicillin. For pooled screens using K562 cells cultured in flasks in a shaking incubator, the culture media was supplemented with 0.1% Pluronic F-127 (Thermo Fisher P6866).

HEK 293T cells were cultured in media consisting of DMEM, high glucose (Gibco 11965084, containing 4.5g/mL glucose and 4mM L-glutamine) supplemented with 10% FBS (VWR) and 100units/mL streptomycin, 100mg/mL penicillin. Adherent cells are routinely passaged and harvested by incubation with 0.25% Trypsin-EDTA (Thermo Fisher 25200056) at 37deg. C for 5-10min, followed by neutralization with media containing 10% FBS.

Unless otherwise specified below, lentiviral particles were produced by transfecting standard packaging vectors into HEK293T using TransIT-LT1 Transfection Reagent (Mirus, MIR2306). At <24 hours post-transfection culture media with exchanged with fresh media supplemented with ViralBoost (Alstem Bio, cat# VB100) at 1:500 dilution. Viral supernatants were harvested ∼48-72 hours after transfection and filtered through a 0.45 mm PVDF syringe filter and either stored in 4deg. C for use within <2 weeks or stored in −80deg. C until use. Lentiviral infections included polybrene (8 mg/ml). MOI was estimated from the fraction of transduced cells (based on fluorescence marker positivity) by the following equation ^108,109^: MOI = -ln(1-fraction of cells transduced).

For experiments described in Fig. S8, lentivirus was produced by transfecting HEK293T cells with lentiviral vector, VSVG and psPAX2 helper plasmids using polyethylenimine. Media was changed ∼6–8 h post transfection. Viral supernatant was collected every 12 h for 5 times and passed through 0.45 µm PVDF filters. Lentivirus was added to target cell lines with 8 µg/mL Polybrene and centrifuged at 650 × g for 25 min at room temperature. Media was replaced 15 h post infection. Antibiotics (1 µg/mL puromycin) was added 48 h post infection.

### Antibody staining and flow cytometry

Antibodies used: CD55-APC (Biolegend 311312), CD81-PE (Biolegend 349506), B2M-APC (Biolegend 316311), KIT-PE (Biolegend 313204), FOLH1-APC (Biolegend 342508), CD56(NCAM1)-APC (Invitrogen 17-0567-42). Cells were stained with antibodies in 96-well plates, using 500g 5min at 4deg. C for centrifugation steps and decanting in between each step. Cells were washed once with 200ul with FACS Buffer (PBS with 1% BSA), then resuspended in 50ul of antibodies diluted at 1:100 in FACS Buffer for 30min at 4deg. C. Then 150ul of FACS Buffer was added, followed by centrifugation and supernatant, then washed one more time with 200ul FACS buffer, followed by final resuspension in 200ul FACS Buffer for flow cytometry. For CRISPRi experiments, all data points shown in figures are events first gated for single cells based on FSC/SSC, then gated on GFP-positivity as a marker for cells successfully transduced with crRNA construct. Flow cytometry was performed on the Attune NxT instrument unless otherwise specified.

For cell fitness competition assays, the percentage of cells expressing the GFP marker encoded on the crRNA expression vector is quantified by flow cytometry. log2 fold change of % GFP-positive cells was calculated relative to day 2 (for experiments targeting the Rpa3 locus in Fig. S8) or day 6 (for experiments targeting the MYC locus in Fig. 6B). For experiments targeting the Rpa3 locus, flow cytometry was performed on the Guava Easycyte 10 HT instrument.

### Indel analysis by Illumina short-read sequencing

200,000 cells were collected on day 14 after crRNA transduction and genomic DNA was isolated using NucleoSpin Blood (Macherey-Nagel, Catalog no. 740951.50). Briefly, PCRs for loci of interest were run using Amplicon-EZ (Genewiz) partial IlluminaÒ adapters and amplicons were processed using NucleoSpin Gel and PCR Clean-up Kit (Macherey-Nagel, Catalog no. 740609.250). Paired end (2 x 250 bp) sequencing was completed at GENEWIZ (Azenta Life Sciences). Raw fastq files were obtained from GENEWIZ and aligned to reference sequences using *CRISPResso2* ^110^ with the following modifications:

--quantification_window_size 12

--quantification_window_center −3

CRISPResso --fastq_r1 R1.fastq.gz --fastq_r2 R2.fastq.gz --amplicon_seq

acccgtcttgtttgtcccacccttggtgacgcagagccccagcccagaccccgcccaaagcactcatttaactggtattgcggagccacgaggcttctgcttactgcaactcgctccggccgctgggcgtagctgcgactcggcggagtcccggcggcgcgtccttgttctaacccggcgcgccatgaccgtcgcgcggccgagcgtgcccgcggcgctgcccctcctcggggagctgccccggctgctgctgctggtgctgttgtgcctgccggccgtgtggggtgagtaggggcccggcggccggggaagcccctgggctgggtgggaggtccaagtcggtctctgaga -g actggtattgcggagccacgagg -wc −3 -w 12

For crRNA constructs in which the PAM is found on the opposite strand with respect to the amplicon sequence (in this case, CD81) the following modifications were included:

--quantification_window_size 20

--quantification_window_center −18

CRISPResso --fastq_r1 pCH45H-CD81-array_S5_L001_R1_001.fastq.gz --fastq_r2 pCH45H-CD81-array_S5_L001_R2_001.fastq.gz --amplicon_seq

ctgcttcgcggggacgaggggggggctcgcgggcgggactcctggcgccccgcccccatgagctcatcaagagccgccgcccctggatggtggggcgggggcgcacactttgccggaggttgggggcgatccgcctcactctttccccagcccagctcactctccaatctgcggtcaccacccgagaccttcctgggggtcgcgcctaaaaggagcgcagactcccgccgggatggcccagaagctggggtgcgcgcaccctggccgtccctgcctgggagccgatctccctctcctcacccagacacgttccagcggaggcctcctcccagaagggctctggaggcctcgcaggagtggggatcccgcggttctgagttgg -p 3 -g gagaccttcctgggggtcgcgcc -wc −18 -w 20

Quantification diagrams were generated in R.

For analysis of dual cutting at the *KIT* TSS, briefly, DNA was isolated using QuickExtract DNA Solution (Lucigen) and amplicons were generated using 15 cycles of PCR to introduce Illumina sequencing primer binding sites and 0-8 staggered bases to ensure library diversity. After reaction clean-up using ExoSAP-IT kit (Thermo Fisher 78201), an additional 15 cycles of PCR was used to introduce unique dual indices and Illumina P5 and P7 adaptors. Libraries were pooled and purified by SPRIselect magnetic beads before paired-end sequencing using an Illumina MiSeq at the Arc Institute Multi-omics Technology Center. Sequencing primer binding sites, unique dual indices (from Illumina TruSeq kits), P5 and P7 adaptor sequences are from Illumina Adaptor Sequences Document # 1000000002694 v16.

Reads were analyzed using CRISPRessoBatch from CRISPResso2 ^110^ with the following modifications: wc −4 -w 15

CRISPRessoBatch --batch_settings batch2.batch --amplicon_seq

aagagcaggggccagacgCCGCCGGGAAGAAGCGAGACCCGGGCGGGCGCGAGGGAGGGGAGGCGAGGAGGGGCGTGGCCGGCGCGCAGAGGGAGGGCGCTGGGAGGAGGGGCTGCTGCTCGCCGCTCGCGGCTCTGGGGGCTCGGCTTTGCCGCGCTCGCTGCACTTGGGCGAGAGCTGGAACGTGGACCAGAGCTCGGATCCCATCGCAGCTACCGCGATGAGAGGCGCTCGCGGCGCCTGGGATTTTCTCTGCGTTCTGCTCCTACTGCTTCGCGTCCAGACAGGTGGGACACCGCGGCTGGCACCCCGACCGTGcgactactcggcgaagcc tgtg -p 3 -g TCTGCGTTCTGCTCCTACTGCTT -wc −4 -w 15

For dual gRNA cutting, both guides were included in the batch analysis.The total number of insertions and deletions at each amplicon position were calculated and displayed using the effect_vector_combined.txt output.

Frequencies of nucleotide substitutions with multiAsCas12a-KRAB targeting are negligible and indistinguishable from sequencing error (<4.5%) observed in unmodified K562 cells.

### Simulations of indel impacts on gene expression

The fraction of reads containing indels within each specified region was subjected to technical noise background subtraction by the fraction of reads containing indels observed in K562 parental cells. This background-subtracted indel allelic frequency was used to calculate per-copy deletion probabilities in Fig. S14 and Fig. S15B. For denAsCas12a-KRAB this background-subtraction can result in a small negative value and in those cases 0% is reported as per-copy deletion probability.

We simulated the impact of indels on gene expression under the assumption that gene expression changes are entirely driven by indels generated by the given AsCas12a fusion protein at the crRNA target site. A prior study reported at one Cas9 target site that the frequency of larger (>250bp) indels is ∼20% relative to the smaller (<250bp) indels. This 20:80 ratio of unobserved-to-observed indels is very likely a high overestimate in our case because our PCR amplicons are 340bp-382bp and thus are expected to capture a large fraction of even the >250bp indels. Nevertheless we added an additional 20% to the observed indel frequencies to arrive at our final estimates of probability of the occurrence of any ≥1bp indels per DNA copy for a single target site. Based on this indel probability we calculated the proportion of cell population expected to harbor a given number of DNA copies with indels assuming indels occur independently among DNA copies and using previously measured DNA copy number for each genomic locus ^111^. For dual targeting of the KIT locus by crKIT-2 and crKIT-3 we assume that the occurrence of a large (>250bp) unobserved deletion at one target site precludes a deletion at the other target site. Starting from the single-cell expression distributions obtained by flow cytometry from the non-targeting crRNA control, we simulated the expected change in single-cell expression distribution under the assumption that any ≥1bp indel in the PCR amplicon would generate a null allele completely abolishing expression of the target gene in cis (i.e. indel in 1 out of 3 copies would reduce expression by 33%). We refer to this as the “expected null” expression change, which is expected to be a high overestimate of impact on expression. To better more accurately estimate the true hypomorphic effects of indels we calculated a “hypomorphic coefficient” defined as the ratio of the observed median expression change vs. the expected null expression change for opAsCas12a. We multiply the expected null expression change for all other fusion proteins by this hypomorphic coefficient to derive an “expected hypomorph” expression change for each fusion protein and crRNA construct combination.

### Nanopore long-read sequencing analysis of deletion frequencies

Genomic DNA was harvested from 20 million cells using the Qiagen Genomic Tips Kit (cat #10243). We used a custom protocol adapted from the Nanopore Cas9 Sequencing Kit user’s manual (SQK-CS9109, though this kit was not actually used) to enrich for genomic DNA surrounding crRNA target sites for Nanopore sequencing using Kit 14 chemistry. Except where specified below, reagents are from the Nanopore Ligation Sequencing Kit V14 (SQK-LSK114). To summarize in brief, synthetic spCas9 Alt-R crRNAs targeting ∼20kb regions surrounding target sites of crCD55-4, crCD81-1, crKIT-2, crKIT-3, crB2M-1, and crB2M3 were designed according to instructions for the Nanopore Cas9 Sequencing Kit and ordered from IDT. At least two guides were used to target the (+) strand upstream of the Cas12a crRNA target sites, and at least two guides were used to target the (-) strand downstream of the target sites. Synthetic spCas9 guides were pooled and annealed with spCas9 tracrRNA (IDT Cat # 1072532) by heating to 95deg. C for 5min and cooling on the bench at room temperature, followed by assembly into ribonucleoprotein complexes by incubating the following mixture at room temperature for 30min: 10ul annealed crRNA:tracrRNA pool (100uM), 10ul 10x NEB rCutSmart Buffer (NEB B6004S), 79.2ul nuclease-free water, and 0.8ul HiFi Cas9 (62uM, S. pyogenes HiFi Cas9 nuclease V3 IDT Cat#1081060). 5ug of genomic DNA was dephosphorylated in a 30ul reaction volume using 15U of Quick calf intestinal phosphatase (NEB cat #M0525) and 3ul of 10x rCutsmart Buffer (NEB B6004S), incubated for 37deg. C for 10min, 80deg. C for 2min, and hold at 20deg. C. The entire volume of dephosphorylated genomic DNA is then mixed with 10ul of Cas9 ribonucleoprotein, 1ul 10mM dATP, and 1ul Taq polymerase for a total of 42ul, incubated at 37deg. C for 20min, 72 deg. C for 5min, and held at 4deg. C. The reaction is then cleaned up using 21ul (0.5X) AMPure XP using two washes with 200ul 80% ethanol, and eluted in 61ul of water. Adapters ligation was set up as follows: 60ul DNA eluate, 25ul Ligation Buffer, 10ul NEBNExt Quick T4 DNA Ligase (NEB E6056S), and 5ul Ligation Adapter, and incubated for 40min at room temperature. The entire ligation reaction was subjected to AMPure XP bead clean up using 40ul (0.4X) of AMPure beads, two washes with Long Fragment Buffer, and eluted in 15ul Elution Buffer. Size distribution and concentration of library was checked by Agilent Tapestation Genomic DNA ScreenTape Analysis. ∼75ng-560ng of each library per flow cell was each run on 1 or 2 flow cells on a Promethion P2 Solo, yielding ∼100k reads per flow cell run over 40h-72h.

fastq files generated by the MiniKNOW software were aligned to the ∼20kb regions (defined by the outermost Cas9 sgRNA protospacer sites flanking each targeted locus) surrounding each crRNA target site in MiniKNOW to generate bam files. Bam files for each sample were merged using *samtool merge* (samtools v1.6). Merged bam files were filtered for alignments that overlap the start and end coordinates of the protospacer region of the Cas12a crRNA using *bamtools filter -region* (bamtools v2.5.1). Filtered bam files were loaded into the Integrative Genomics Viewer 2.17.0 for visualization of individual read alignments. *pysamstats –fasta –type variation* (pysamstats v1.1.2) was used to extract per base total read coverage and deletion counts. The fraction of aligned reads harboring a deletion at each base was plotted using custom scripts in R.

### Pooled crRNA library design

For all crRNAs in Library 1 and Library 2: we excluded in the analysis spacers with the following off-target prediction criteria using CRISPick run in the CRISPRi setting: 1) off-target match = ‘MAX’ for any tier or bin, or 2) # Off-Target Tier I Match Bin I Matches > 1). The only crRNAs for which this filter was not applied are the non-targeting negative control spacers, which do not have an associated CRISPick output. All crRNA sequences were also filtered to exclude BsmbI sites used for cloning and >3 consecutive T’s, which mimic RNA Pol III termination signal.

#### Library 1 (single crRNA’s)

To design crRNA spacers targeting gene TSS’s for Library 1, we used the −50bp to +300bp regions of TSS annotations derived from capped analysis of gene expression data and can include multiple TSS’s per gene ^68^. We targeted the TSS’s of 559 common essential genes from DepMap with the strongest cell fitness defects in K562 cells based on prior dCas9-KRAB CRISPRi screen ^68^. We used CRISPick with enAsCas12a settings to target all possible PAM’s (TTTV and non-canonical) in these TSS-proximal regions. Except for the criteria mentioned in the previous paragraph, no other exclusion criteria were applied. For the TSS-level analyses shown in Fig. 4D-E, each gene was assigned to a single TSS targeted by the crRNA with the strongest fitness score for that gene.

Negative controls in Library 1 fall into two categories: 1) 524 intergenic negative controls, and 2) 445 non-targeting negative controls that do not map to the human genome. Target sites for intergenic negative controls were picked by removing all regions in the hg19 genome that are within 10kb of annotated ensembl genes (retrieved from biomaRt from https://grch37.ensembl.org) or within 3kb of any ENCODE DNase hypersensitive site (wgEncodeRegDnaseClusteredV3.bed from http://hgdownload.cse.ucsc.edu/goldenpath/hg19/encodeDCC/wgEncodeRegDnaseClustered/). The remaining regions were divided into 1kb fragments. 90 such 1kb fragments were sampled from each chromosome. Fragments containing >=20 consecutive N’s were removed. The remaining sequences were submitted to CRISPick run under CRISPRi settings. The CRISPick output was further filtered for spacers that meet these criteria: 1) off-target prediction criteria described in the beginning of this section, and 2) On-target Efficacy Score >= 0.5 (the rationale is to maximize representation by likely active crRNAs to bias for revealing any potential cell fitness effects from non-specific genotoxicity due to residual DNA cutting by multiCas12a-KRAB), 3) mapping uniquely to the hg19 genome by Bowtie ^112^ using ‘-m 1’ and otherwise default parameters, 3) filtered once more against those whose uniquely mapped site falls within 10kb of annotated ensembl genes or any ENCODE DNase hypersensitive site.

Non-targeting negative control spacers were generated by combining 1) non-targeting negative controls in the Humagne C and D libraries, 2) taking 20nt non-targeting spacers from the dCas9-KRAB CRISPRi_v2 genome-wide library ^68^, removing the G in the 1st position, and appending random 4-mers to the 3’ end. This set of spacers were then filtered for those that do not map to the hg19 genome using Bowtie with default settings.

#### Library 2 (6-plex cRNA’s)

Sublibrary A (84,275 constructs): Test position spacers were encoded at each position of the 6-plex array, with remaining positions referred to as context positions and filled with negative control spacers. Test positions encodes one of 506 intergenic negative control spacers and 2,303 essential TSS-targeting spacers. The essential TSS-targeting spacers were selected from among all spacers targeting PAM’s within −50bp to +300bp TSS-proximal regions of 50 common essential genes with the strongest K562 cell fitness defect in prior dCas9-KRAB CRISPRi screen ^68^, and must have ≥0.7 CRISPick On-target Efficacy Score. Negative control context spacers consist of 5 6-plex combinations, 3 of these combinations consist entirely of non-targeting negative controls and 2 of the combinations consist entirely of intergenic negative controls.

Sublibrary B (6,370 constructs): crRNA combinations targeting cis-regulatory elements at the MYC locus were assembled from a subset of combinations possible from 15 starting spacers (3 targeting MYC TSS, 3 targeting each of 3 enhancers, and 3 intergenic negative control spacers). The 3 enhancer elements are described in the subsection “Design of individual crRNAs.” These 15 starting spacers were grouped into 5 3-plex combinations, each 3-plex combination exclusively targeting one of the 4 cis-regulatory elements, or consisting entirely of intergenic negative controls. Each 3-plex was then encoded in positions 1-3 of 6-plex arrays, and positions 4-6 were filled with all possible 3-plex combinations chosen from the starting 15 spacers. All 6-plex combinations were also encoded in the reverse order in the array.

All-negative control constructs (2000 constructs): 1500 6-plex combinations were randomly sampled from the intergenic negative control spacers described for Library 1. 500 6-plex combinations were randomly sampled from non-targeting negative control spacers described for Library 1.

Intergenic negative controls and non-targeting negative controls are defined the same as in Library 1.

As Library 2 was designed and cloned prior to the completion of the Library 1 screen, the majority of Library 2 contains constructs encoding for spacers in the test position that in hindsight do not produce strong phenotypes as single crRNAs in the Library 1 screen.

### crRNA library construction

For Library 1, ∼140 fmol of pooled oligo libraries from Twist were subjected to 10 cycles of PCR amplification using primers specific to adaptor sequences flanking the oligos and containing BsmbI sites. The PCR amplicons were cloned into a crRNA expression backbone (pCH67) by Golden Gate Assembly with ∼1:1 insert:backbone ratio using ∼500 fmol each. Golden Gate Assembly reaction was carried out in a 100ul reaction containing 2.5U Esp3I (Thermo ER0452) and 1000U T4 DNA Ligase in T4 DNA Ligase reaction buffer (NEB M0202L). The reaction mix was incubated for 31 cycles alternating between 37deg. C and 16deg. C for 20min at each temperature, then heat-inactivated at 65deg. C for 5min. Assembly reactions were column purified with Zymo DNA clean and concentrator-5 (Zymo D4004), eluted in 12ul of water and <7ul added to 70ul of MegaX DH10B T1R Electrocomp Cells (C640003) for electroporation using BioRad Gene Pulser Xcell Electroporator with settings 2.0kV, 200ohms, 25μF. Cells were recovered at 37deg. C for rotating for 1h in ∼5ml recovery media from the MegaX DH10B T1R Electrocomp Cells kit and small volumes plated onto bacterial LB plates containing carbenicillin for quantification of colony forming units. The remaining recovery culture was inoculated directly into 200ml liquid LB media with carbenicillin and incubated in 37deg. C shaker for 12h-16h prior to harvesting for plasmid purification using ZymoPURE II Plasmid Midiprep kit (Zymo D4200). Based on the colony forming units from the small volumes in the bacterial plates, the estimated coverage of the library is 778x. 24 individual colonies were verified by Sanger sequencing and the library subjected to deep sequencing as described in Illumina sequencing library preparation. For Library 2, 915 fmol of pooled oligo libraries from Twist was subjected to 18 cycles of PCR amplification and agarose gel purification of the correctly sized band before proceeding similarly with the remainder of the protocol as described above. The estimated coverage of the library from colony forming units is ∼60x.

### Illumina sequencing library preparation

crRNA inserts were amplified from genomic DNA isolated from screens using 16 cycles of first round PCR using pooled 0-8nt staggered forward and reverse primers, treated with ExoSAP-IT (Thermo Fisher 78201.1.ML), followed by second round of PCR to introduce Illumina unique dual indices and adaptors. Sequencing primer binding sites, unique dual indices, P5 and P7 adaptor sequences are from Illumina Adaptor Sequences Document # 1000000002694 v16. PCR amplicons were subject to size selection by magnetic beads (SPRIselect, Beckman B23318) prior to sequencing on an Illumina NovaSeq6000 using SP100 kit for Library 1 or SP500 kit for Library 2. Sequencing of plasmid libraries were performed similarly, except 7 cycles of amplification were each used for Round 1 and Round 2 PCR. The size distribution of the final library was measured on an Agilent TapeStation system. We noted that even after magnetic bead selection of Round 2 PCR-amplified Library 2 plasmid library (colonies from which were Sanger sequencing verified) and genomic DNA from screens, smaller sized fragments from non-specific PCR amplification during Illumina sequencing library preparation persisted. This might contribute to the fraction of reads that could not be mapped to our reference 6-plex array. Thus, these unmapped reads do not necessarily reflect recombination of the crRNA library constructs, though the latter could contribute as well.

### Cell fitness screens

Library 1 screen: K562 cells engineered by piggyBac transposition to constitutively express denAsCas12a-KRAB or multiAsCas12a-KRAB were transduced with lentivirally packaged Library 1 constructs at MOI = ∼0.15. Transduced cells were then selected using 1ug/ml puromycin for 2 days, followed by washout of puromycin. On Day 6 after transduction, initial (T0) time point was harvested, and the culture was split into 2 replicates that are separately cultured henceforth. 10 days later (T10), the final time point was harvested (8.6 total doublings for multiAsCas12a-KRAB cells, 9.15 total doublings for denasCas12a-KRAB cells). A cell coverage of >500x was maintained throughout the screen. Library 2 screen: K562 cells engineered by piggyBac transposition to constitutively express multiAsCas12a-KRAB were transduced with lentivirally packaged Library 2 constructs at MOI = ∼0.15. The screen was carried out similarly as described for Library 1 screen, except the screen was carried out for 14 days (T14) or 13.5 total doublings and maintained at a cell coverage of >2000x throughout. Genomic DNA was isolated using the NucleoSpin Blood XL Maxi kit (Machery-Nagel 740950.50).

### Screen data processing and analysis

Summary of library contents are in Fig. S18.

Library 1: Reads were mapped to crRNA constructs using sgcount (https://noamteyssier.github.io/sgcount/), requiring perfect match to the reference sequence.

Library 2: First, reference construct sequences were created by interspersing provided spacer and constant regions. Each construct is then given a unique construct id (CID). Each CID is then split into R1 and R2 reference sequences, which are constructed by taking the first three and last three spacer-construct pairs of the reference sequence respectively. The R2 sequence is then reverse complemented for matching against the R2 sequencing reads. Next, two hashmaps are created for the R1 and R2 spacer-construct pairs respectively, which map the R1/R2 sequences to a set of corresponding CIDs. Finally, for each R1/R2 sequencing pair, each k-mer (k = length of R1/R2 respective construct sequence) in the sequence is mapped against their respective R1/R2 hashmap. If both sequencing pairs are able to be mapped to a CID set, then the intersection of their sets is their original construct, and the total count of that CID is incremented. We implemented the above algorithm as the ‘casmap constructs’ command in a package written in Rust, available at https://github.com/noamteyssier/casmap.

Starting from read counts, the remainder of analyses were performed using custom scripts in R. Constructs that contained <1 read per million reads (RPM) aligned to the reference library in either replicates at T0 were removed from analysis. From the constructs that meet this read coverage threshold, a pseudocount of 1 was added for each construct and the RPM re-calculated and used to obtain a fitness score ^113^ that can be interpreted as the fractional defect in cell fitness per cell population doubling:

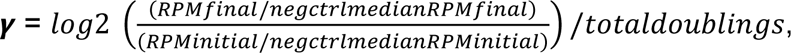

where RPM = read count per million reads mapped to reference (initial = at T0, final = at end of screen), negctrlmedian = median of RPM of intergenic negative control constructs, totaldoublings = total cell population doublings in the screen. For Library 1, data from a single T0 sample was used to calculate the fitness score for both replicates due to an unexpected global loss of sequencing read counts for one of two originally intended T0 replicate samples.

### 3’ RNA-seq experiment and data analysis

#### Experimental procedure

3’ RNA-seq was performed as part of a batch processed using a QuantSeq-Pool Sample-Barcoded 3′ mRNA-Seq Library Prep Kit for Illumina (Lexogen cat#139) in accordance with the manufacturer’s instructions. Briefly, 10 ng of each purified input RNA was used for first strand cDNA synthesis with an oligo(dT) primer containing a sample barcode and a unique molecular identifier. Subsequently, barcoded samples were pooled and used for second strand synthesis and library amplification. Amplified libraries were sequenced on an Illumina HiSeq4000 with 100 bp paired-end reads. The QuantSeq Pool data was demultiplexed and preprocessed using an implementation of pipeline originally provided by Lexogen (https://github.com/Lexogen-Tools/quantseqpool_analysis). The final outputs of this step are gene level counts for all samples (including samples from multiple projects multiplexed together).

#### Gene level and differential expression analysis

For generating scatter plots, normTransform function from DESeq2 ^114^ used to normalize the raw counts and then a pseudocount of 1 added, and log2-transformed. The output was plotted in R as scatter plots.

For differential expression analysis, DESeq2 (version 1.34) default Wald-test was used to compare each targeting construct (one replicate) with non-targeting samples (two replicates). We calculated log2 transformed TPM counts and applied the threshold of 6.5 to eliminate genes with low expression. Using ggplot2, volcano plots visualized in R are then displayed for genes with log2FoldChange above or below 2.055 and p-values smaller than 0.01. The log2FoldChange cutoff was based on visually examining the concordance between two replicates of untransduced controls and manually identifying a threshold below which the log2FoldChange are poorly correlated between the replicates of the untransduced control.

#### Off-target analysis of spacers

To evaluate potential off-target effect of spacers, we used the crisprVerse (version 1.0.0) ^115^ and crisprBowtie (version 1.2.0) together with other R packages including GenomicRanges (version 1.50) ^116^ and tidyverse (version 1.3.2) ^117^. First, we defined dictionary of spacers as “TCCTCCAGCATCTTCCACATTCA”:“HBG-2”,

“TTCTTCATCCCTAGCCAGCCGCC”:“HBG-3”,

“CTTAGAAGGTTACACAGAACCAG”:“HS2-1”,

“TGTGTAACCTTCTAAGCAAACCT”:“HS2-2”,

“AGGTGGAGTTTTAGTCAGGTGGT”:“HS2-3”,

“ATTAACTGATGCGTAAGGAGATC”:“NT-3”. Then, ‘runCrisprBowtiè function used with these parameters: ‘crisprNucleasè as ‘enAsCas12à, ‘n_mismatches’ equal 3, ‘canonical’ equal FALSE, and ‘bowtie_index’ as a path to folder including pre-indexed hg38 reference genome. Thus, the results from this step allows us to assess our previously designed spacers and annotate potential off-target loci in the human genome. To annotate results, we used reference annotation GENCODE (version 34) ^118^ and we defined pam_site +/− 2500 bp for each predicted off-target to overlap them with matched transcription start sites (TSS) +/− 1000 bp of all annotated genes. Results shown as annotated tables.

### RT-qPCR

For the CRISPRi experiments targeting the HBG TSS or HS2 enhancer, K562 cells engineered (by lentiviral transduction at MOI ∼5) for constitutive expression of multiAsCas12a-KRAB were transduced with crRNAs and sorted, followed by resuspension of ∼200k to 1 million cells in 300ul RNA Lysis Buffer from the Quick-RNA Miniprep Kit (Zymo R1055) and stored in −70deg. C. RNA isolation was performed following the kit’s protocols, including on-column DNase I digestion. 500ng of RNA was used as input for cDNA synthesis primed by random hexamers using the RevertAid RT Reverse Transcription Kit (Thermo fisher K1691), as per manufacturer’s instructions. cDNA was diluted 1:4 with water and 2ul used as template for qPCR using 250nM primers using the SsoFast EvaGreen Supermix (BioRad 1725200) on an Applied Biosystems ViiA 7 Real Time PCR System. Data was analyzed using the ddCT method, normalized to GAPDH and no crRNA sample as reference.

### Transient transfection experiments

For co-transfection experiments, transfections were performed similar to prior study ^27^. Briefly, the day before transfection, 100,000 HEK293T cells were seeded into wells of a 24 well plate. The following day, we transiently transfected 0.6µg of each protein construct and 0.3µg gRNA construct per well (in duplicate) in Mirus TransIT-LT1 transfection reagent according to manufacturer’s instructions. Mixtures were incubated at room temperature for 30 minutes and then added in dropwise fashion into each well. 24 hours after transfection, cells were replenished with fresh media. 48 hours after transfection, BFP and GFP positive cells (indicative of successful delivery of protein and crRNA constructs) were sorted (BD FACSAria Fusion) and carried out for subsequent flow-cytometry experiments.

## Data access and availability

Tables of all sequence and read counts are included as Supplementary Files. Plasmids will be made available on Addgene. Engineered cell lines will be made available upon request.

## Supplemental Figures

**Figure S1.**
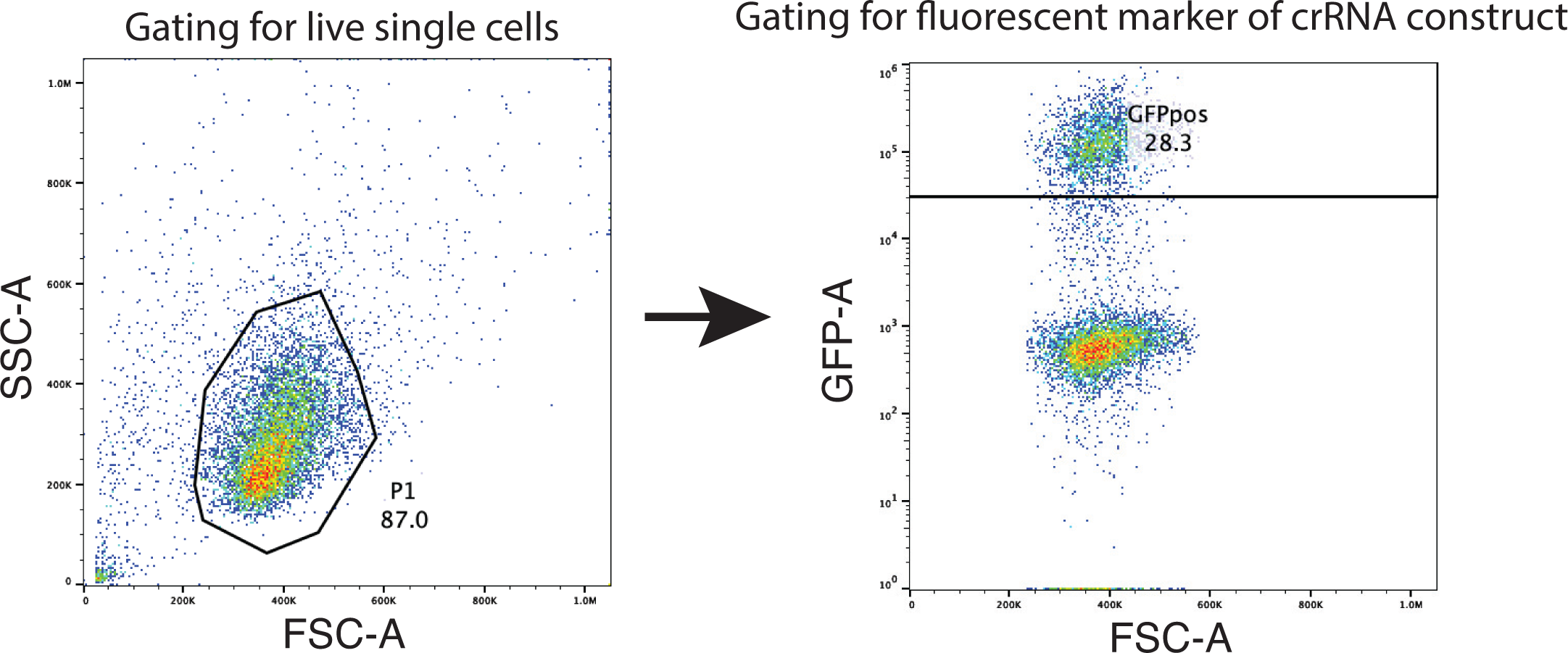
– Example of flow cytometry gating strategy for CRISPRi experiments. Example of the general gating strategy for CRISPRi experiments using flow cytometry readouts, shown for K562 cells as an example. Single live cells are gated by FSC vs. SSC, followed by gating for the fluorescent marker on the crRNA construct (typically GFP), which are subsequently analyzed for target gene expression in the respective fluorescence channels.

**Figure S2.**
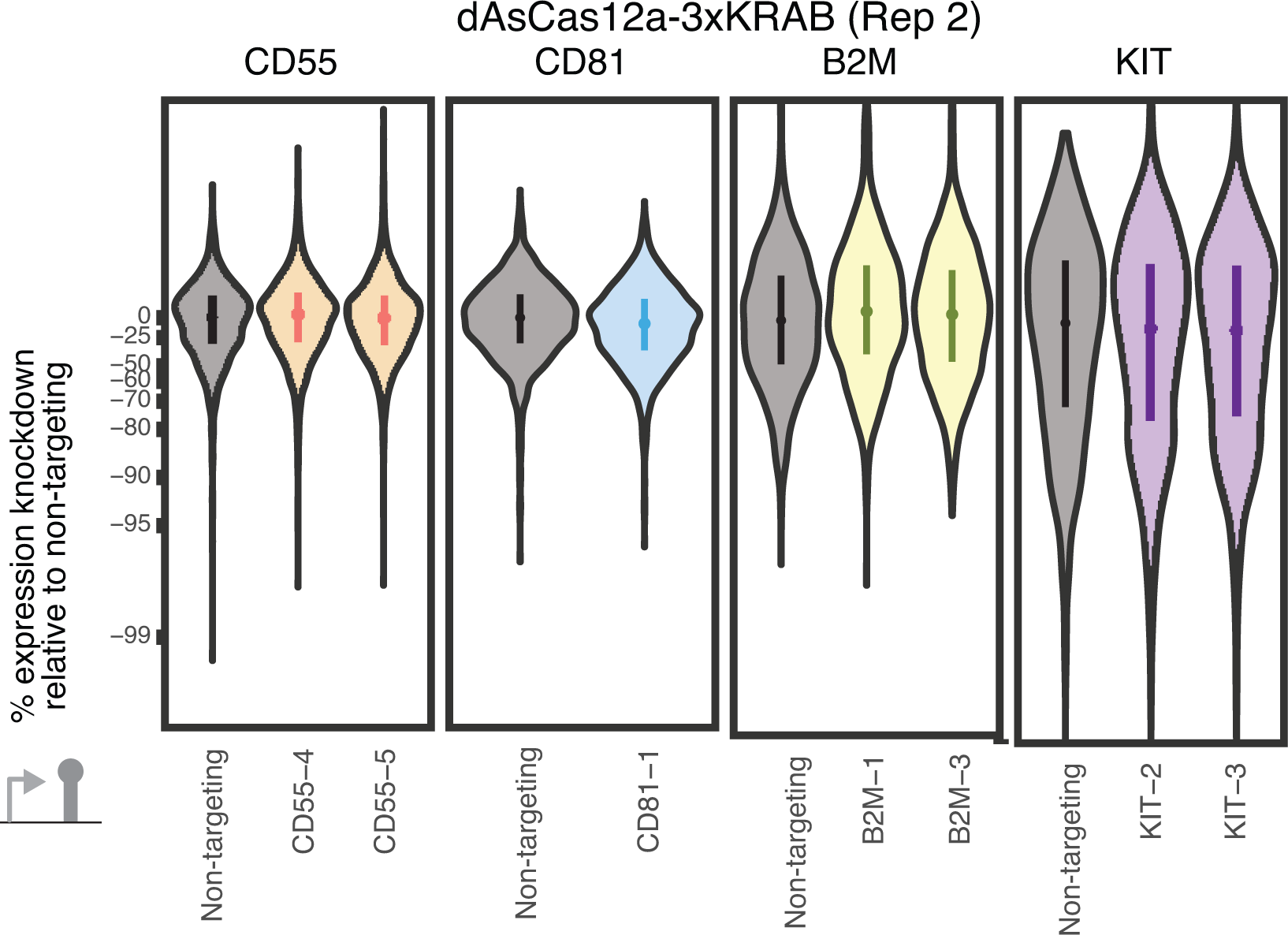
– Additional replicate testing dAsCas12a-3xKRAB CRISPRi activity. Additional replicate for Fig. 1B testing CRISPRi activity of K562 cells piggyBac-engineered to constitutively express dAsCas12a-3xKRAB and using lentivirally delivered crRNA constructs. See Fig. 1B for details.

**Figure S3.**
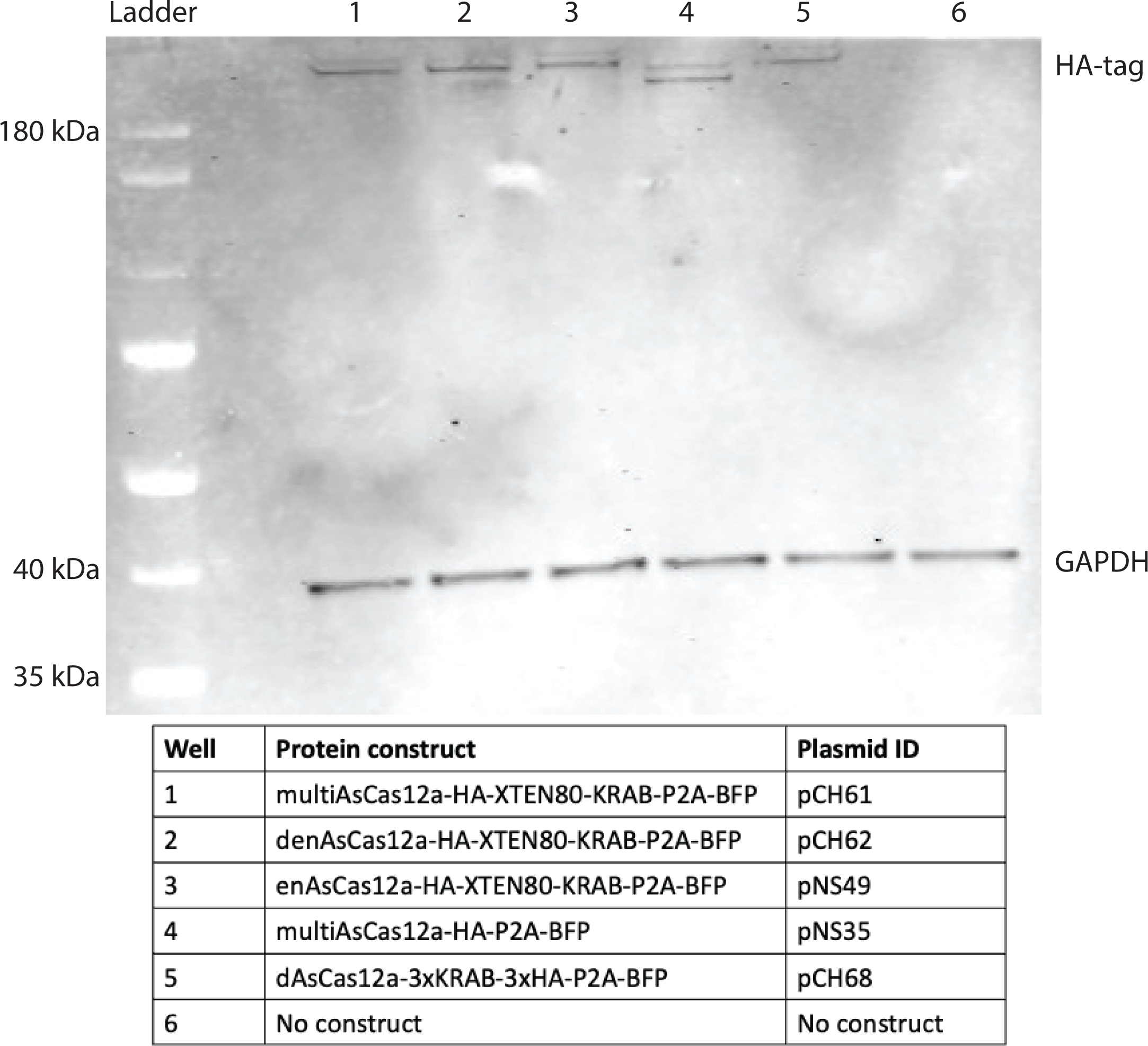
–Western blot of fusion proteins. Western blot of whole-cell lysates prepared from K562 cells piggyBac engineered to constitutively express each of the fusion proteins in the panel. anti-HA tag was used for detection of the fusion protein and anti-GAPDH for detection of GAPDH as loading control.

**Figure S4.**
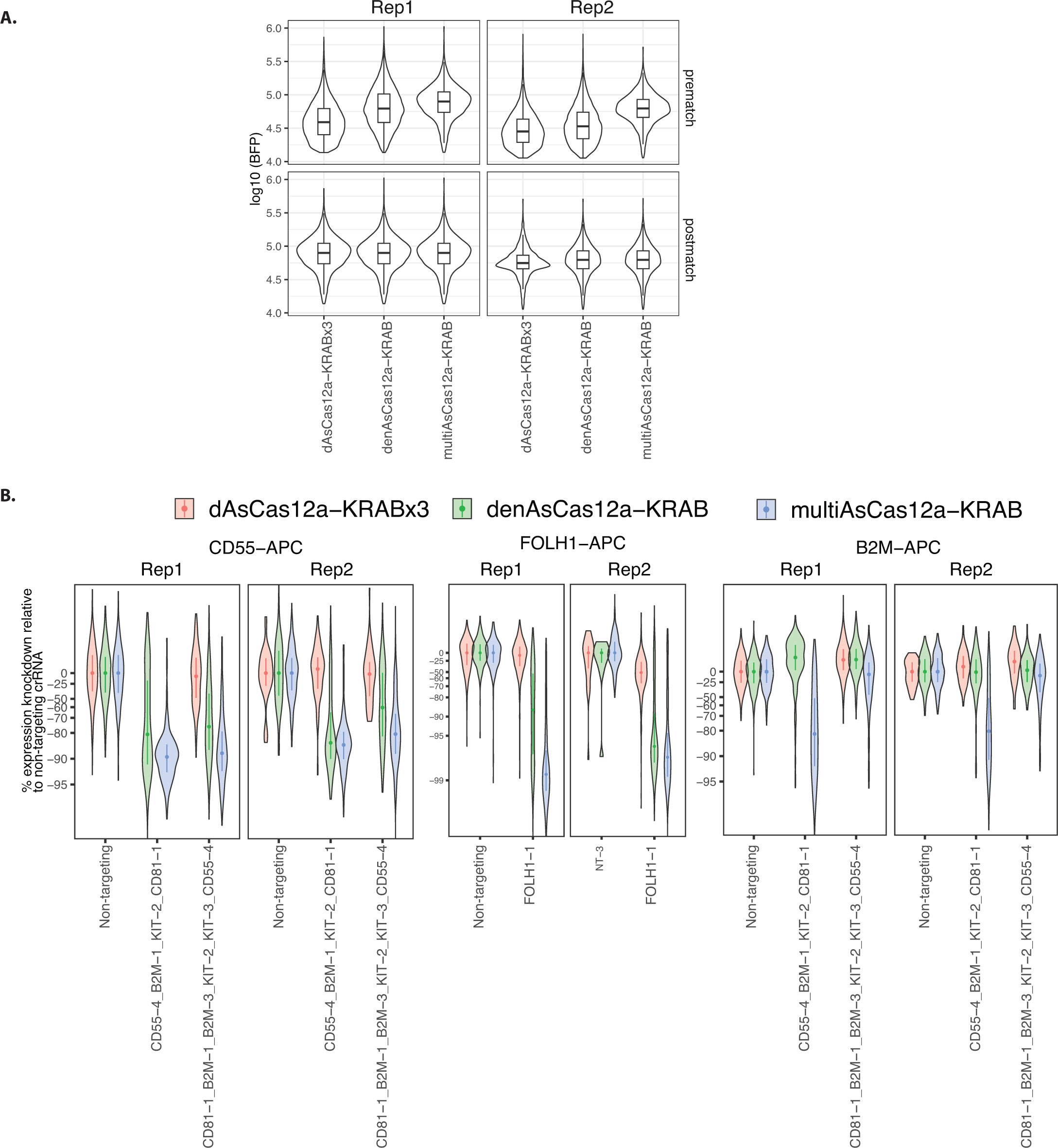
– CRISPRi activity of multiCas12a-KRAB, denAsCas12a-KRAB, and dAsCas12a-KRABx3 in C42B cells. **A)** C4-2B cells piggyBac-engineered to constitutively express each of the fusion protein constructs are lentivirally transduced with the indicated crRNA constructs. Cells were sorted based on P2A-BFP marker signal. Because some of these cell lines showed slightly different levels of BFP signal as a proxy of fusion protein expression, to account for fusion protein expression we performed propensity score matching to subset for populations of cells for each fusion protein construct with the same distributions in BFP signals after data acquisition for flow cytometry in CRISPRi experiments. The BFP signals before and after matching are shown as violin blots overlaid with boxplots showing median and interquartile range. **B)** Target gene expression are measured by cells surface antibody staining and flow cytometry 13-14 days after crRNA transduction. Singlecell distributions of expression knockdown relative to non-targeting crRNA are shown with mean and interquartile range indicated using the cells after propensity score matching for BFP levels as described in A. CRISPRi knockdown results are indistinguishable with and without propensity score matching (not shown).

**Figure S5.**
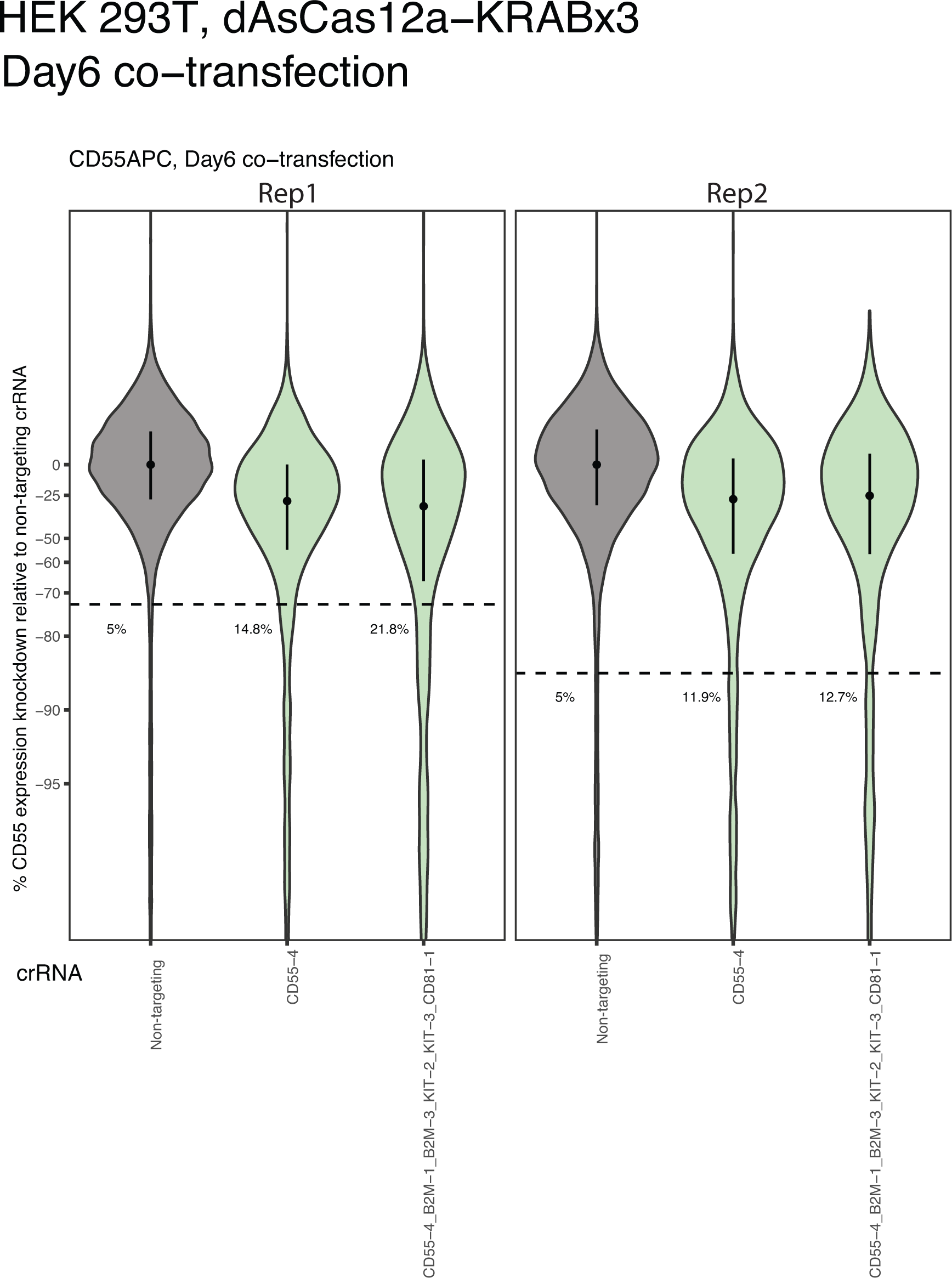
– dAsCas12a-KRABx3 CRISPRi by transient transfection in HEK 293T cells. HEK 293T cells were co-transfected with a plasmid encoding for dAsCas12a-KRABx3 and plasmids encoding for the indicated crRNA constructs targeting CD55. Cells were sorted 2 days after transfection for successful co-transfection based on BFP and GFP markers on the plasmids and CD55 expression was measured by antibody staining on flow cytometry 6 days after transfection. Violin plots of single-cell distributions of CD55 expression knockdown as a percentage of the median of non-targeting control are shown. Median and interquartile range are shown in the plot. The percentage of cells below the 5th percentile of the non-targeting control are also shown.

**Figure S6.**
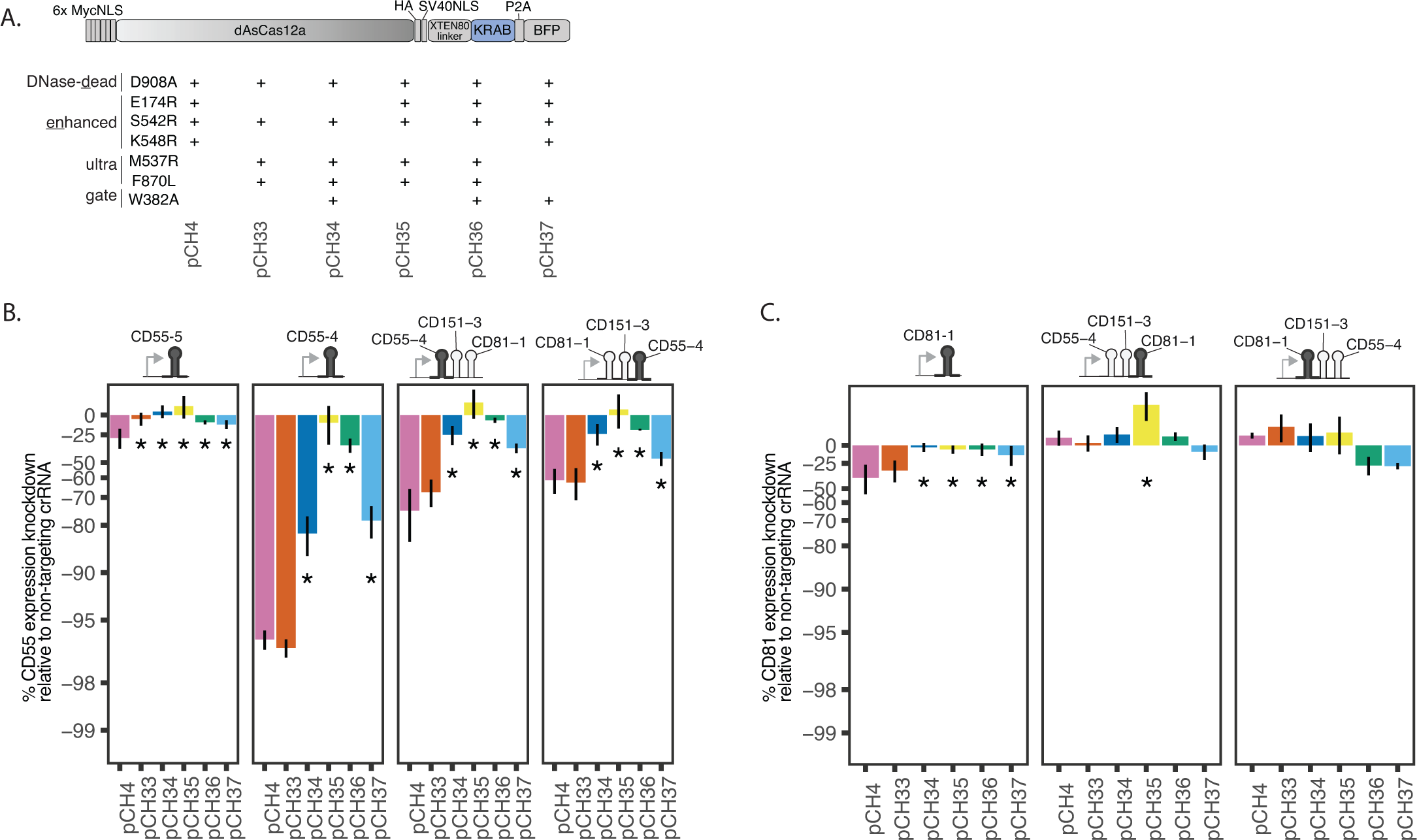
– Comparisons of dAsCas12a variant fusion CRISPRi constructs using up to 3-plex crRNA constructs. **A)** The same fusion protein schematic as shown in Fig. 1C, labeled with construct IDs for ease of reference. **B)** CD55 expression knockdown measured by flow cytometry using the indicated crRNA constructs and the panel of fusion protein constructs in A. Shown are averages of the median single-cell expression knockdown relative to non-targeting crRNA for 3 biological replicates (including the replicate for crCD55-4 shown in Fig. 1C) for all comparisons, except the comparison for crCD81-1 crCD151-3 crCD55-4 contains 2 replicates. One-sided Wilcoxon rank-sum test comparing denAsCas12a-KRAB (pCH4) to each of the other fusion constructs in the panel was performed. Asterisk indicates p*<*0.01 for all replicate-level comparisons for a given construct comparison. **C)** Analogous to B, but for CD81 knockdown. Summaries shown for 3 biological replicates (including the replicate for crCD81-1 shown in Fig. 1C) for all comparisons, except the comparison for crCD81-1 crCD151-3 crCD55-4 contains 2 replicates.

**Figure S7.**
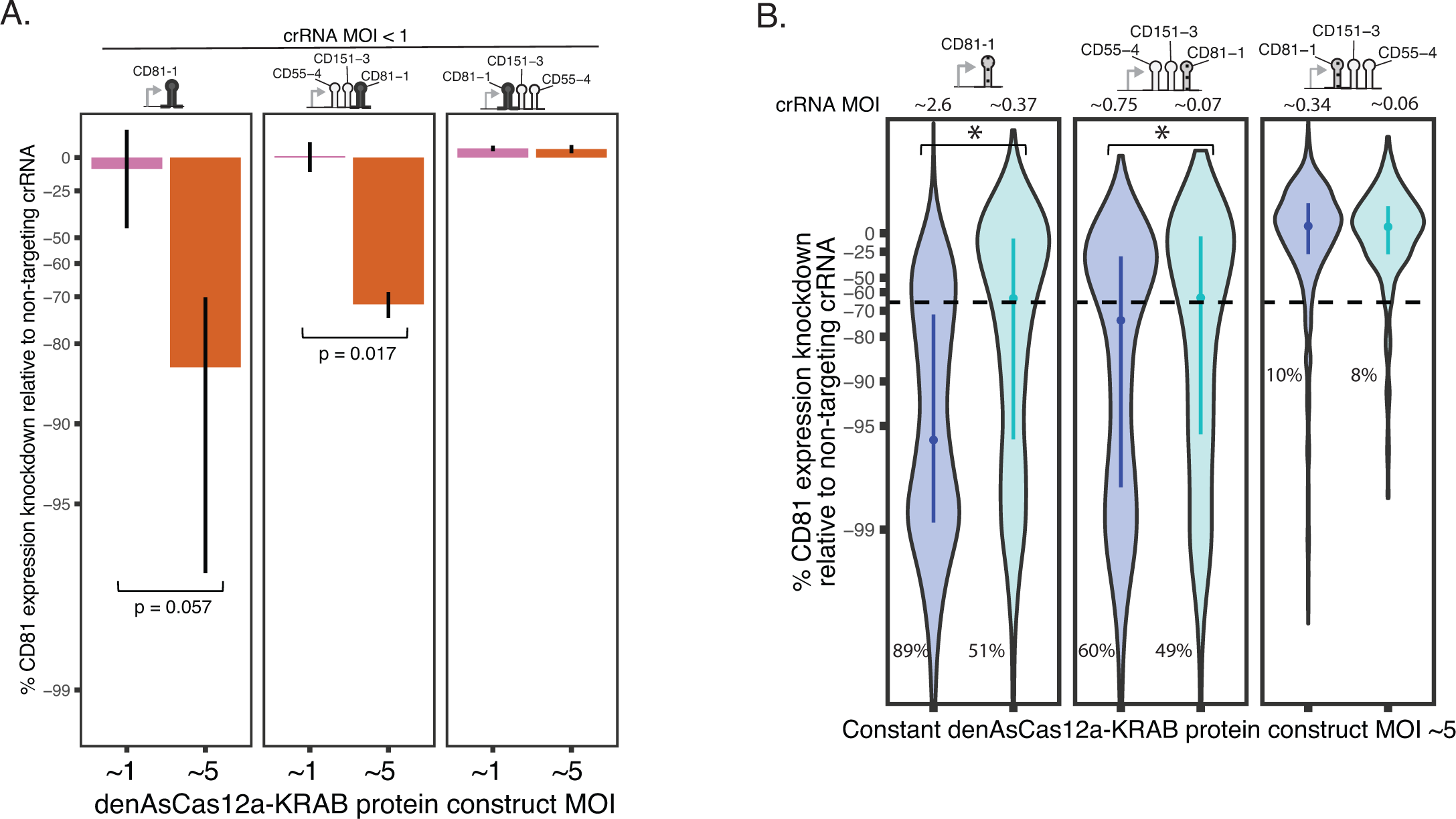
– Additional replicates testing effect of dose on denAsCas12a-KRAB CRISPRi activity. **A)** Summary of all replicates for experiment shown in Fig. 1D: shown are averages of median expression knockdown for each crRNA construct (N = 3-6 biological replicates for each crRNA construct, including the replicate shown in Fig. 1D). Error bars denote SEM. One-sided Wilcoxon rank-sum test was performed on the medians of single-cell expression knockdown of each replicate and p-values indicated where relevant. **B)** Additional biological replicate for Fig. 1E testing denAsCas12a-KRAB CRISPRi activity with varying crRNA MOI while holding protein MOI constant; see Fig. 1E for details.

**Figure S8.**
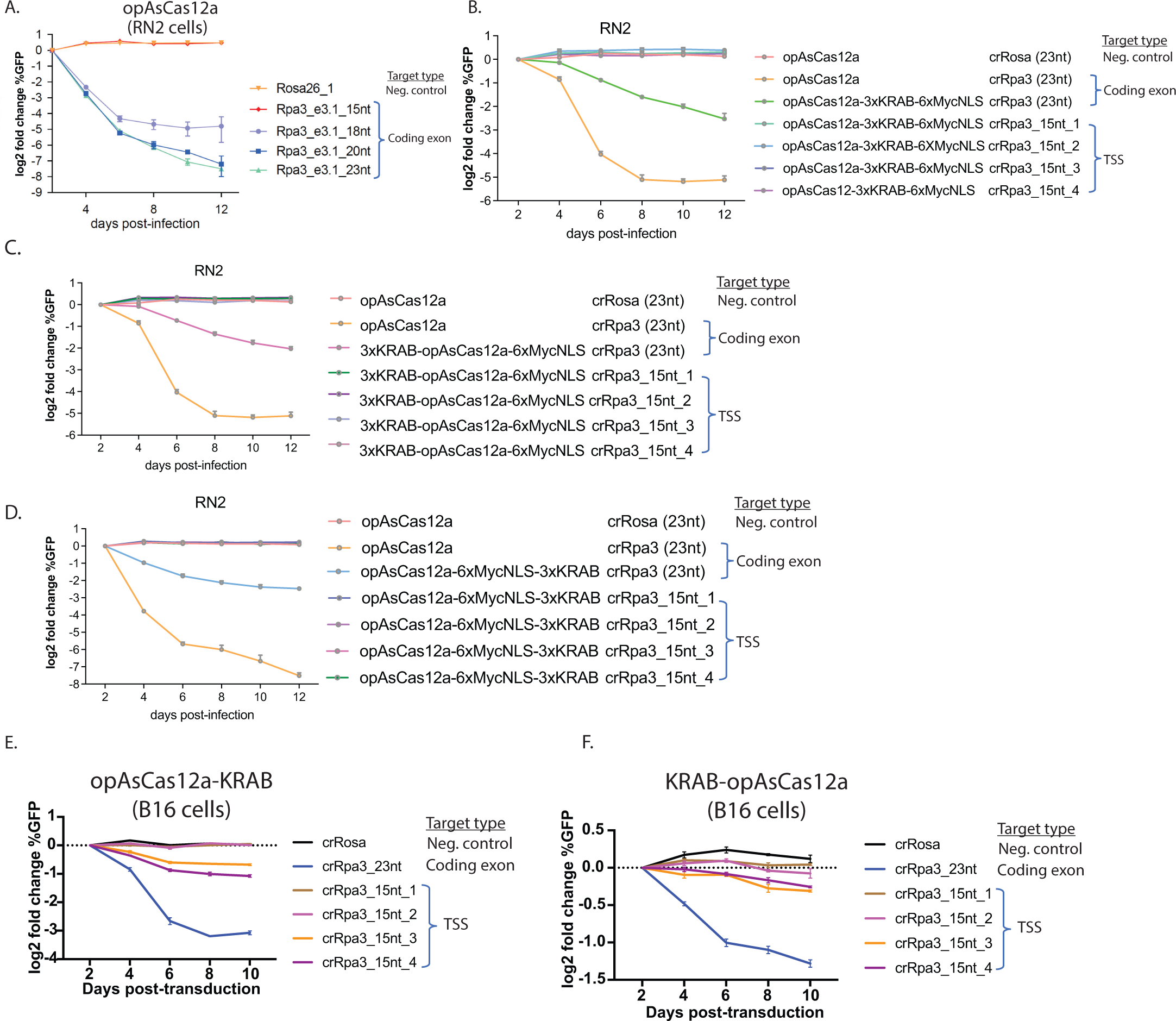
– Testing CRISPRi activity of lentivirally delivered truncated crRNAs. In all panels, the indicated cell line (RN2 or B16) was engineered for constitutive expression of the indicated fusion protein constructs by lentiviral transduction, followed by lentiviral transduction (at MOI between 0.3-0.4) of the indicated single-plex crRNA constructs containing spacers of the indicated lengths targeting Rpa3, an essential gene. The spacers target either the gene’s coding exon, the TSS region, or the Rosa locus (negative control) as indicated in the legends. Cell fitness defect is used as a proxy of Rpa3 gene expression knockdown due to either DNA cutting or CRISPRi activity. Cell fitness is measured in a competition assay by quantifying log2 fold change in percent of cells expressing the GFP marker on the crRNA expression constructs (relative to day 2). Error bars indicate SEM for N = 3 biological replicates for all panels.

**Figure S9.**
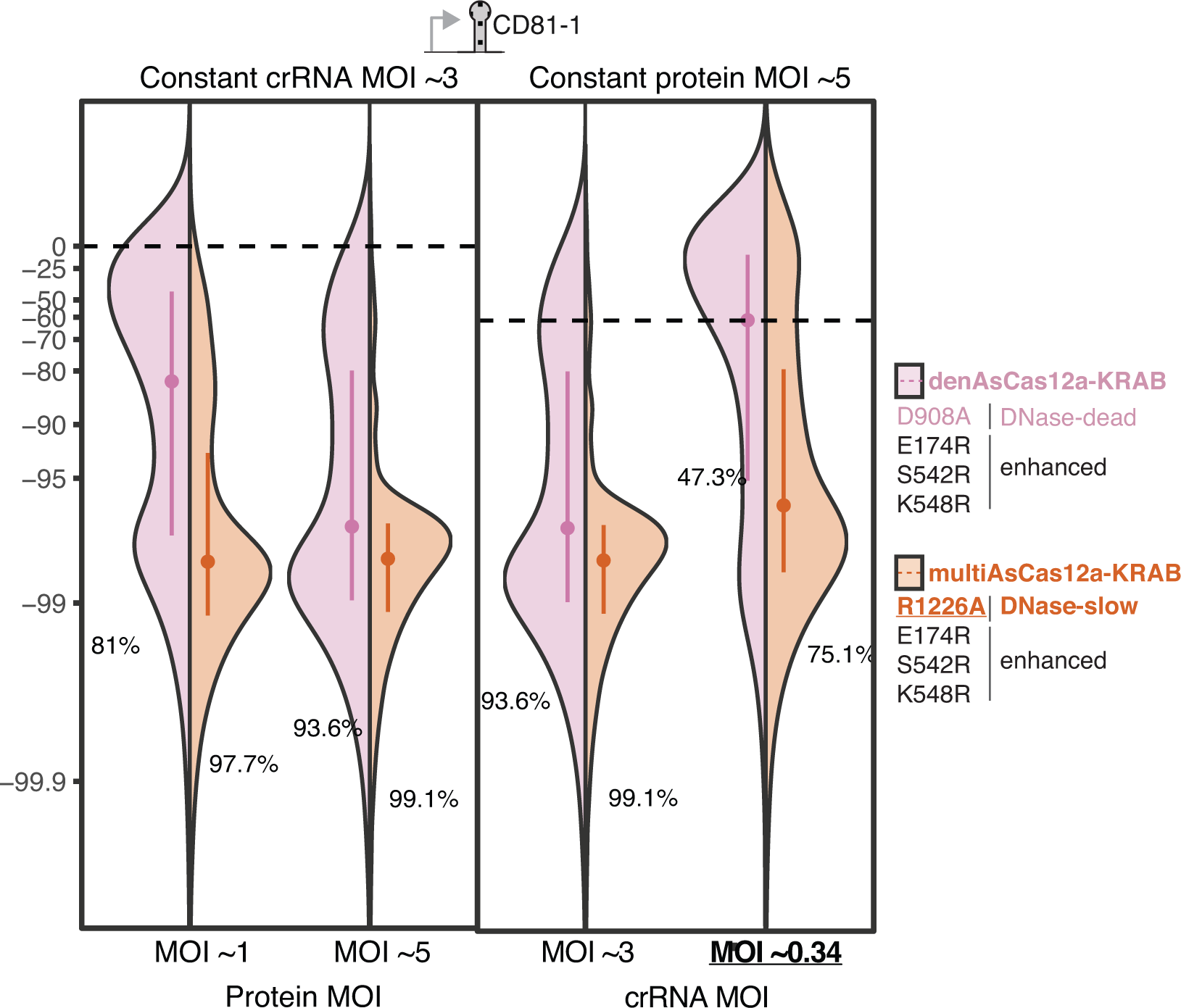
– CD81 knockdown denAsCas12a-KRAB vs. multiAsCas12a-KRAB at different protein and crRNA MOIs. Second biological replicate for testing denAsCas12a-KRAB vs. multiAsCas12a-KRAB CRISPRi activity in the setting of varying protein MOI or crRNA MOI, same as Fig. 2B. See Fig. 2B for further details.

**Figure S10.**
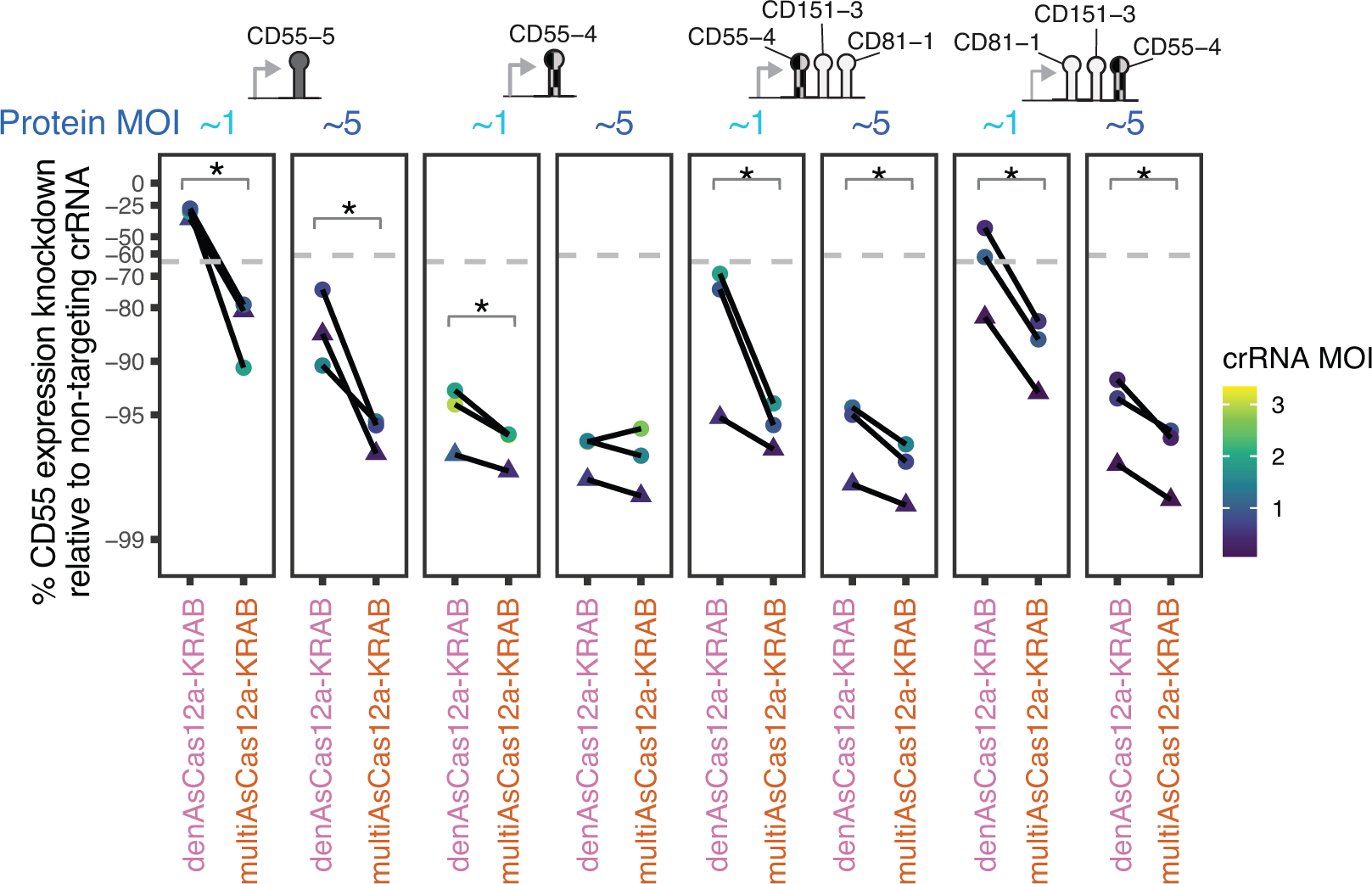
– CD55 knockdown by denAsCas12a-KRAB vs. multiAsCas12a-KRAB at different protein MOIs. Comparison of CD55 knockdown by lentivirally delivered denAsCas12a-KRAB vs. multiAsCas12a-KRAB at protein MOI ∼1 vs. ∼5 across a panel of single and 3-plex crRNA constructs, while holding constant crRNA MOI for each paired fusion protein comparison for each crRNA construct. Dashed gray line indicates 5th percentile of non-targeting crRNA control. crRNA MOI indicated by color scale. Lines connect paired replicates. One-sided Wilcoxon rank-sum tests were performed on single-cell distributions for each replicate, and asterisk denotes p*<*0.01 for all paired replicates within each condition. Dots indicate flow cytometry measurement 10 days after crRNA transduction; triangles indicate flow cytometry measurement 16 days after crRNA transduction.

**Figure S11.**
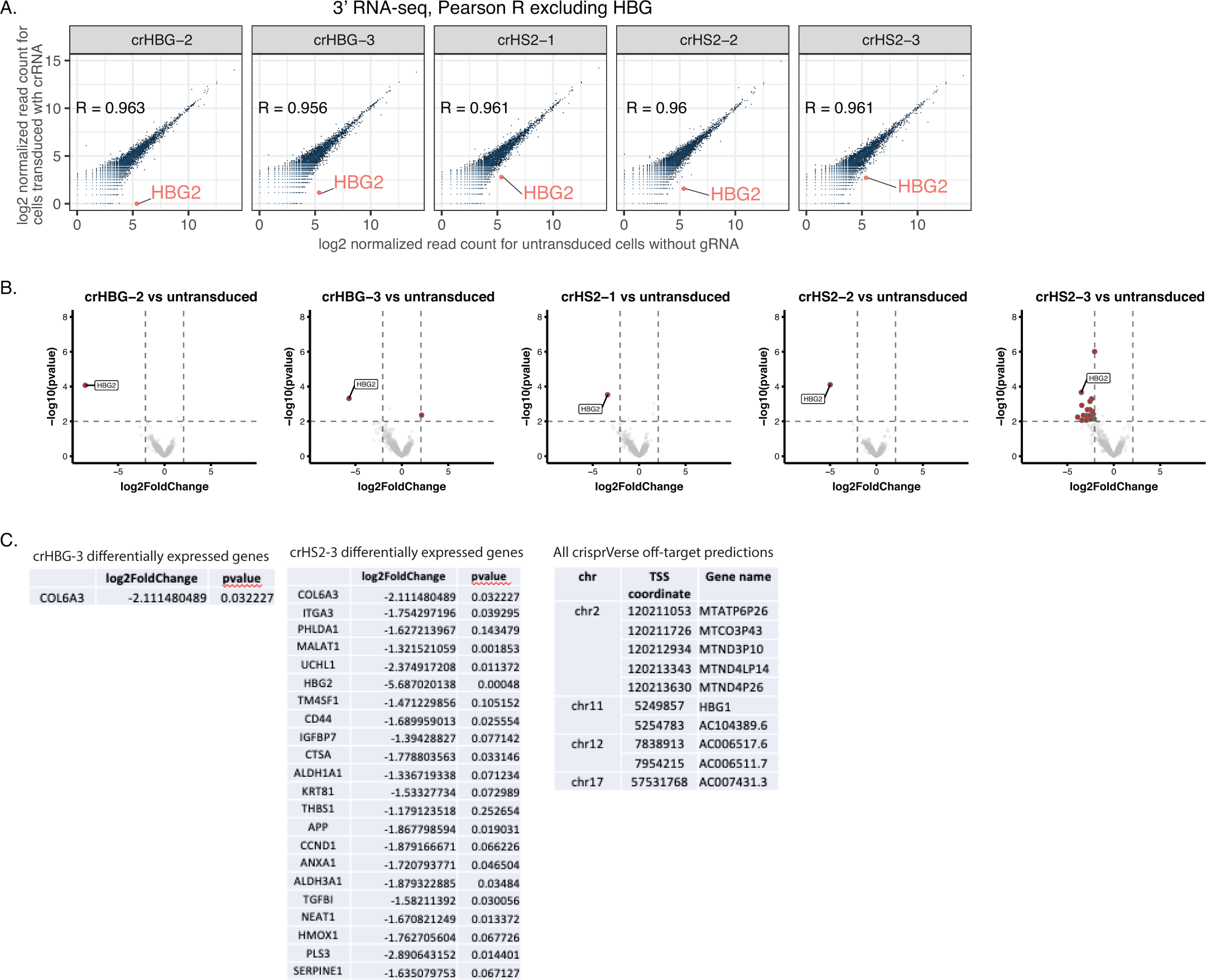
– RNA-seq analysis of crRNA specificity. **A)** K562 cells lentivirally engineered (MOI ∼5) to constitutively express multiAsCas12a-KRAB were either transduced with the indicated crRNA’s at MOI *<*0.3, followed by sorting for crRNA-transduced cells based on GFP marker, or received no crRNAs. RNA was isolated from the sorted cells 32 days of culture after crRNA transduction and subjected to 3’ RNA-seq. Scatter plot of normalized mRNA expression levels for crRNA transduced (1 biological replicate each) vs. cells without crRNA (2 biological replicates), and Pearson correlation coefficient calculated for the transcriptome, excluding HBG. RT-qPCR quantifications are shown in Fig. 5A. **B)** Volcano plots of p-values vs. log2FoldChange from differential expression analysis using DE-seq2 are shown. Genes that fall beyond p-value and log2FoldChange cutoffs (dashed lines) are highlighted. **C)** Lists of differentially expressed genes (other than HBG) from the analysis in B for the crHBG-3 and crHS2-3 transduced cells are shown. For comparison, a list of all off-target predictions generated by crisprVerse are shown for all crRNAs in the panel in A and B.

**Figure S12.**
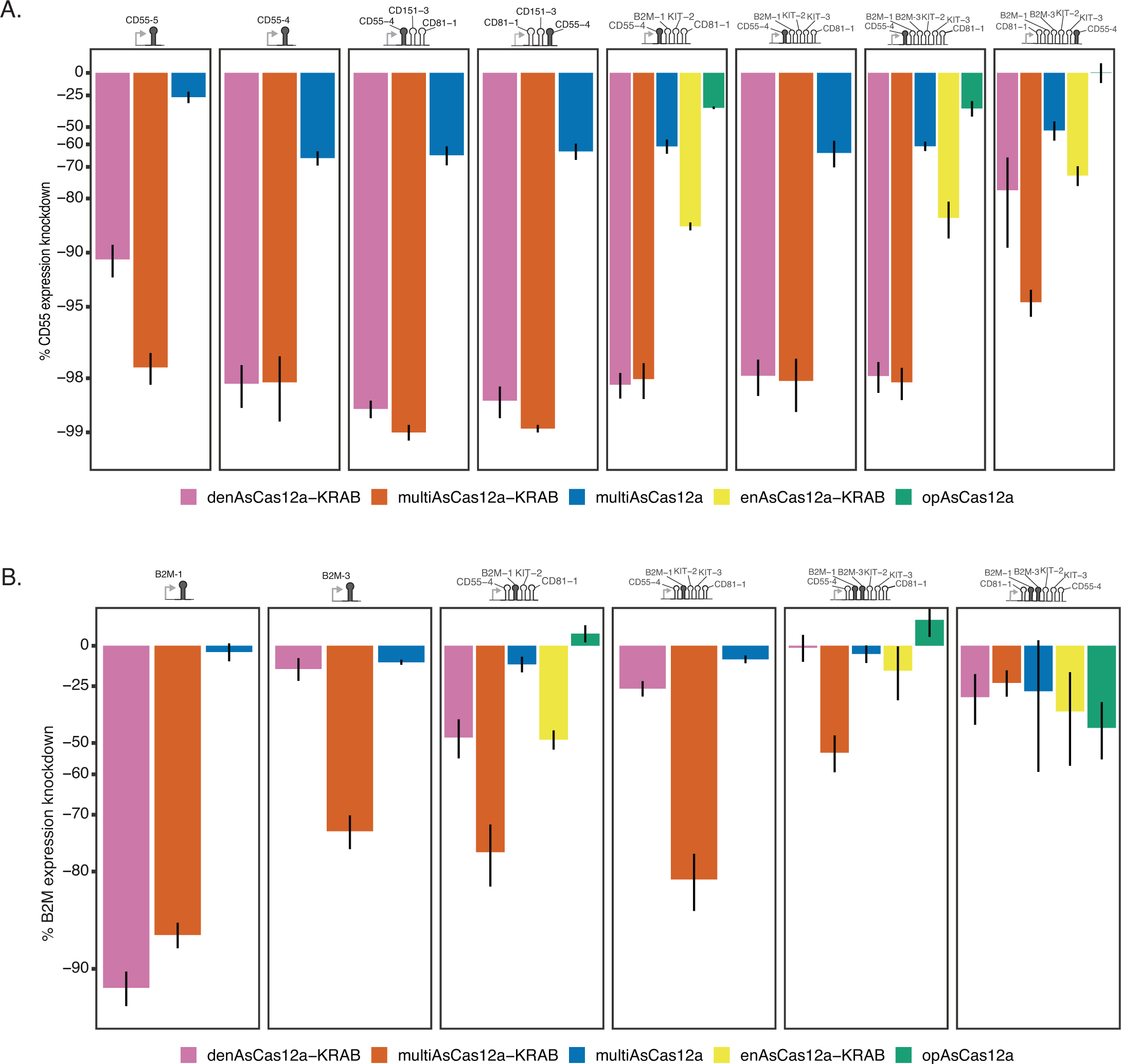
– CRISPRi knockdown of CD55 and B2M using up to 6-plex crRNA arrays. Analogous to Fig. 3B-C, but shown for CD55 and B2M knockdown on day 6 after crRNA transduction, measured by antibody staining of those targets using flow cytometry. Shown are averages of median single-cell expression knockdown from 2-5 biological replicates for each crRNA construct, with error bars indicating SEM.

**Figure S13.**
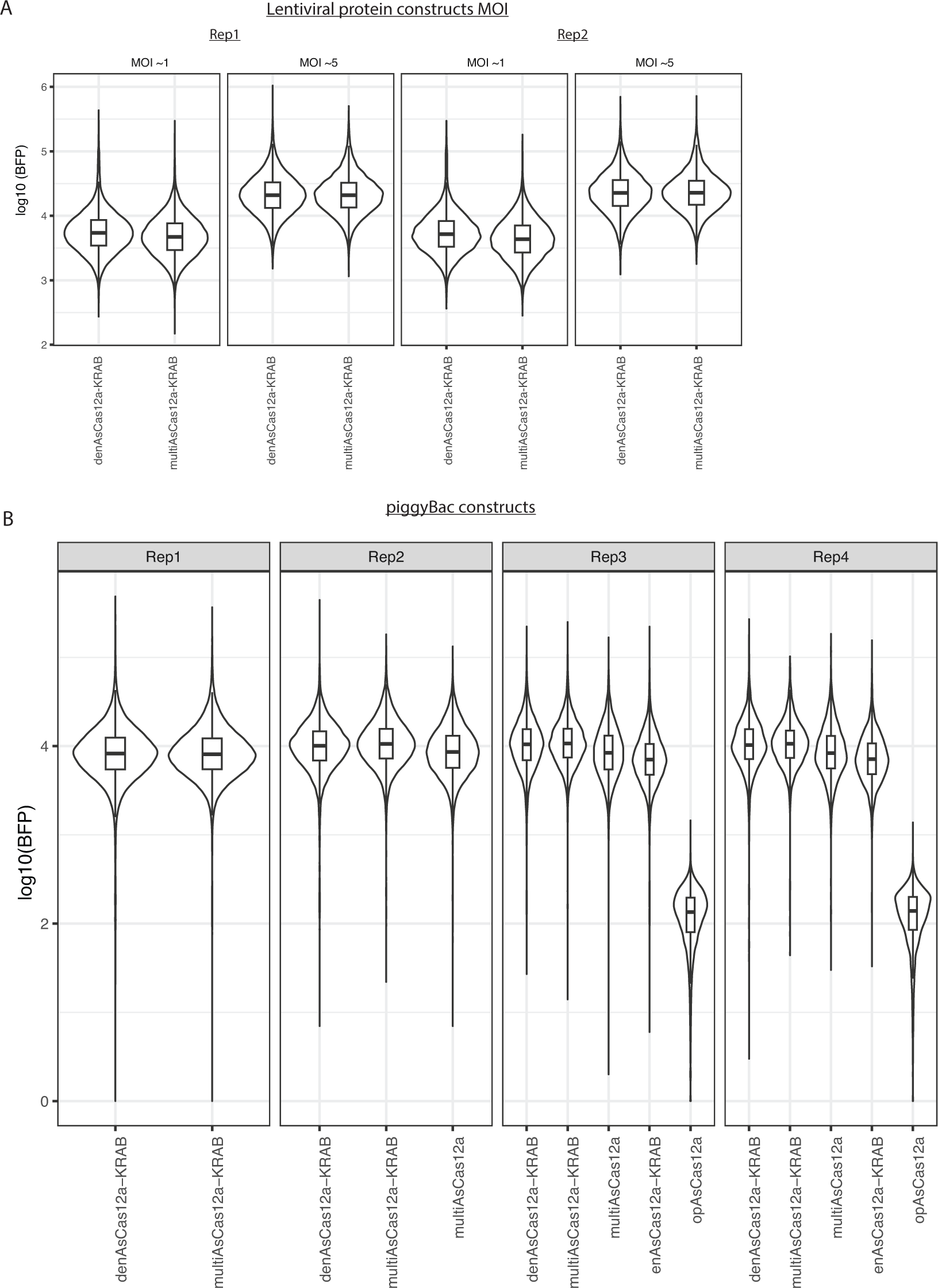
– Monitoring P2A-BFP reporter as proxy of fusion protein expression level. **A)** K562 cells lentivirally engineered (at MOI ∼1 or MOI ∼5) to constitutively express the indicated fusion protein constructs were monitored for P2A-BFP expression levels by flow cytometry. Shown are representative biological replicates from routine monitoring. **B)** Same as A for K562 cells piggyBac-engineered to constitutively express the indicated fusion protein constructs. opAsCas12a does not contain BFP reporter and is shown as fluorescence negative control. Shown are representative biological replicates from routine monitoring.

**Figure S14.**
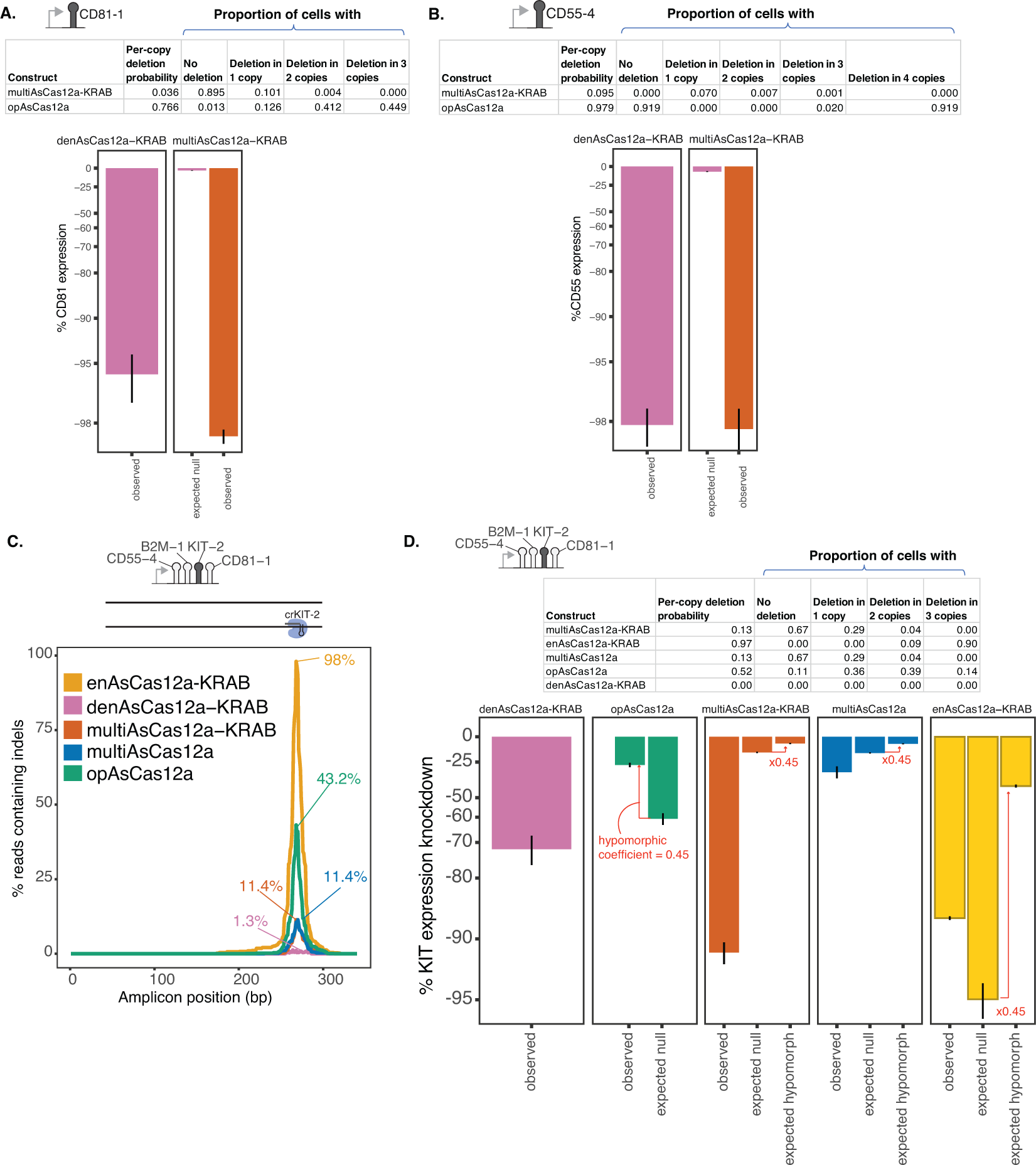
– Indel quantification and gene expression knockdown simulation for single-targeting. **A)** To obtain an upper estimate on the impact on gene expression arising from indels induced by crCD81-1 at the target site, we added an additional 20% unobserved large deletions (upper estimate based on long-read sequencing in Fig. S16A) to the observed indel frequency from short-read sequencing of PCR amplicon (Fig. 2F) to arrive at an estimated per-copy deletion probability. Based on this estimate, the proportions of cells harboring deletions at varying DNA copy numbers are shown assuming independent probability of indel formation across the 3 DNA copies in this region of the K562 genome. Assuming that an indel of any size within the PCR amplicon results in a complete genetic null abolishing CD81 gene expression in cis, we simulate the expected effect of these indels on single-cell distributions of gene expression measured by flow cytometry (”expected null”), compared to the observed gene expression knockdown by crCD81-1 (”observed”). **B)** Analogous to A, but for crCD55-4, which targets a tetraploid region of the K562 genome. **C)** Indel quantification by short-read sequencing of PCR amplicon for a single-site targeting of the KIT TSS region using crKIT-2 encoded within the indicated 4-plex crRNA array. **C)** Analogous to A-B, we simulated expected gene expression knockdown by crKIT-2 single-site targeting based on indel frequencies observed in C. Expected knockdown under this genetic null assumption exceeds that observed for opAsCas12a (fully active DNase), demonstrating the genetic null assumption is an overestimate of gene expression effects of indels in this region. To correct for this overestimate, we use the ratio of observed vs. expected null median expression knockdown by opAsCas12a as an estimate of the hypomorphic effect of deletions in this region (“hypomorphic coefficient”). We multiply the expected null median expression knockdown for all other fusion proteins by this hypomorphic coefficient to obtain an “expected hypormorph” median expression knockdown, which we propose as our final estimate of the effects arising from indels.

**Figure S15.**
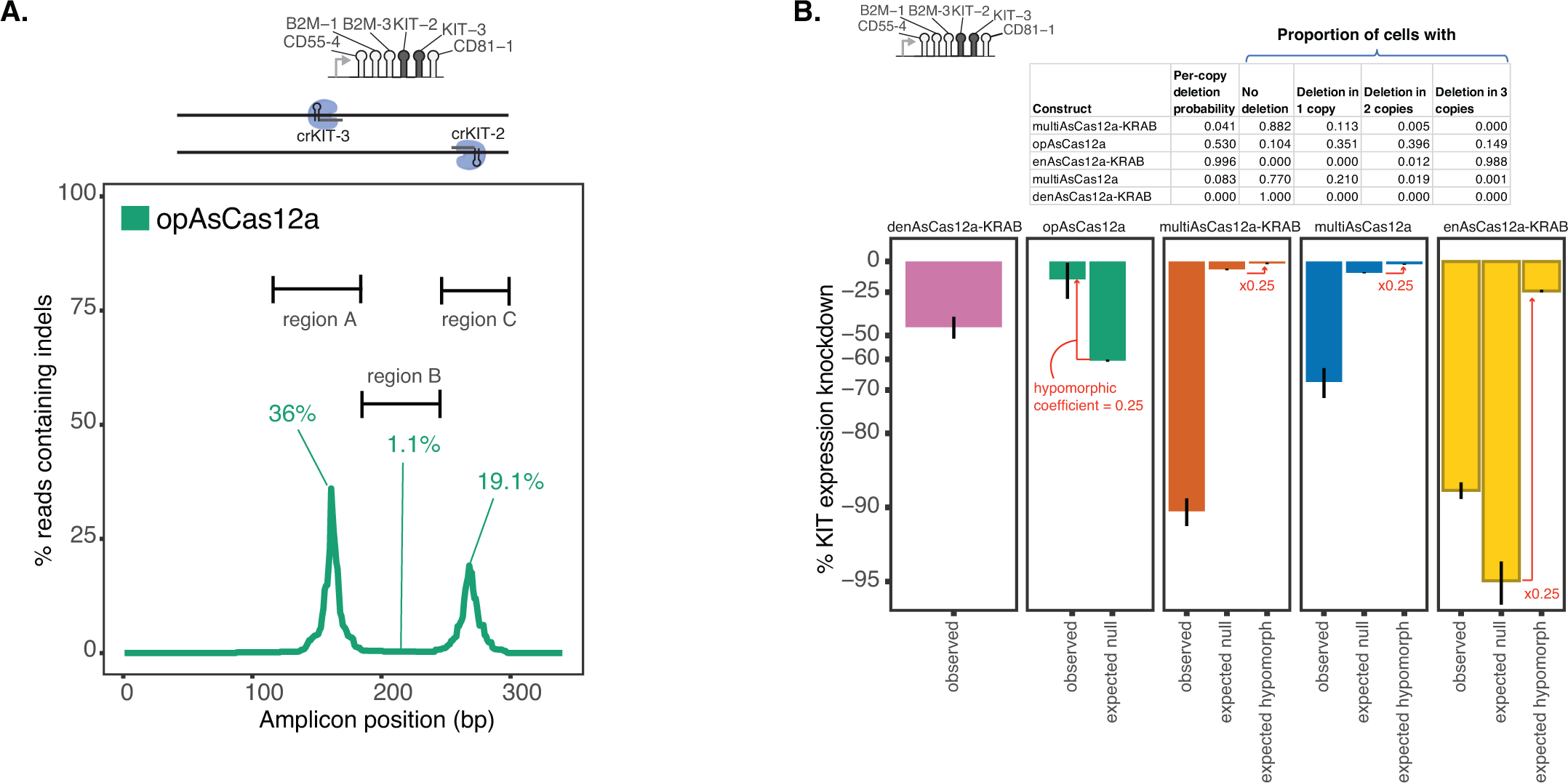
– Indel quantification and gene expression knockdown simulation for dual-targeting of the KIT TSS region. **A)** Related to Fig. 3D. Indel quantification of PCR amplicon near the KIT TSS region in K562 cells lentivirally engineered to constitutively express opAsCas12a 15 days after transduction of the indicated 6-plex crRNA array (sorted for crRNA transduced cells on 2 days after transduction). Note that opAsCas12a is encoded in a different expression backbone using a puromycin selectable marker and thus is not directly matched to other fusion constructs in Fig. 3D in transgenic expression level. **B)** We estimated the per-copy deletion probability occuring any where within the PCR amplicon in the KIT locus in Fig. 3D and Fig. S15A based on the on the observed indel allelic frequencies from short-read sequencing, plus an additional 20% unobserved large deletions (upper estimate based on long-read sequencing in Fig. S16A). A single composite per-copy deletion probability is assigned to the locus assuming independent probabilities of small indels generated separately at the crKIT-2 and crKIT-3 target sites, plus assuming that large deletions at one crRNA target site precludes additional alterations in DNA sequence at the other target site. The proportions of cells that harbor a specified number of DNA copies containing indels of any size in this locus are calculated assuming indels occur independently across DNA copies within each cell. We simulated the expected distribution of single-cell gene expression levels under the assumption that indels of any size in this locus result in a genetic null abolishing KIT expression in cis (”expected null”). The expected knockdown under this genetic null assumption exceeds that observed for opAsCas12a (fully active DNase), demonstrating the genetic null assumption is an overestimate of gene expression effects of indels in this region. To correct for this overestimate, we use the ratio of observed vs. expected null median expression knockdown by opAsCas12a as an estimate of the hypomorphic effect of deletions in this region (“hypomorphic coefficient”). We multiply the expected null median expression knockdown for all other fusion proteins by this hypomorphic coefficient to obtain an “expected hypormorph” median expression knockdown, which we propose as our final estimate of the effects arising from indels.

**Figure S16.**
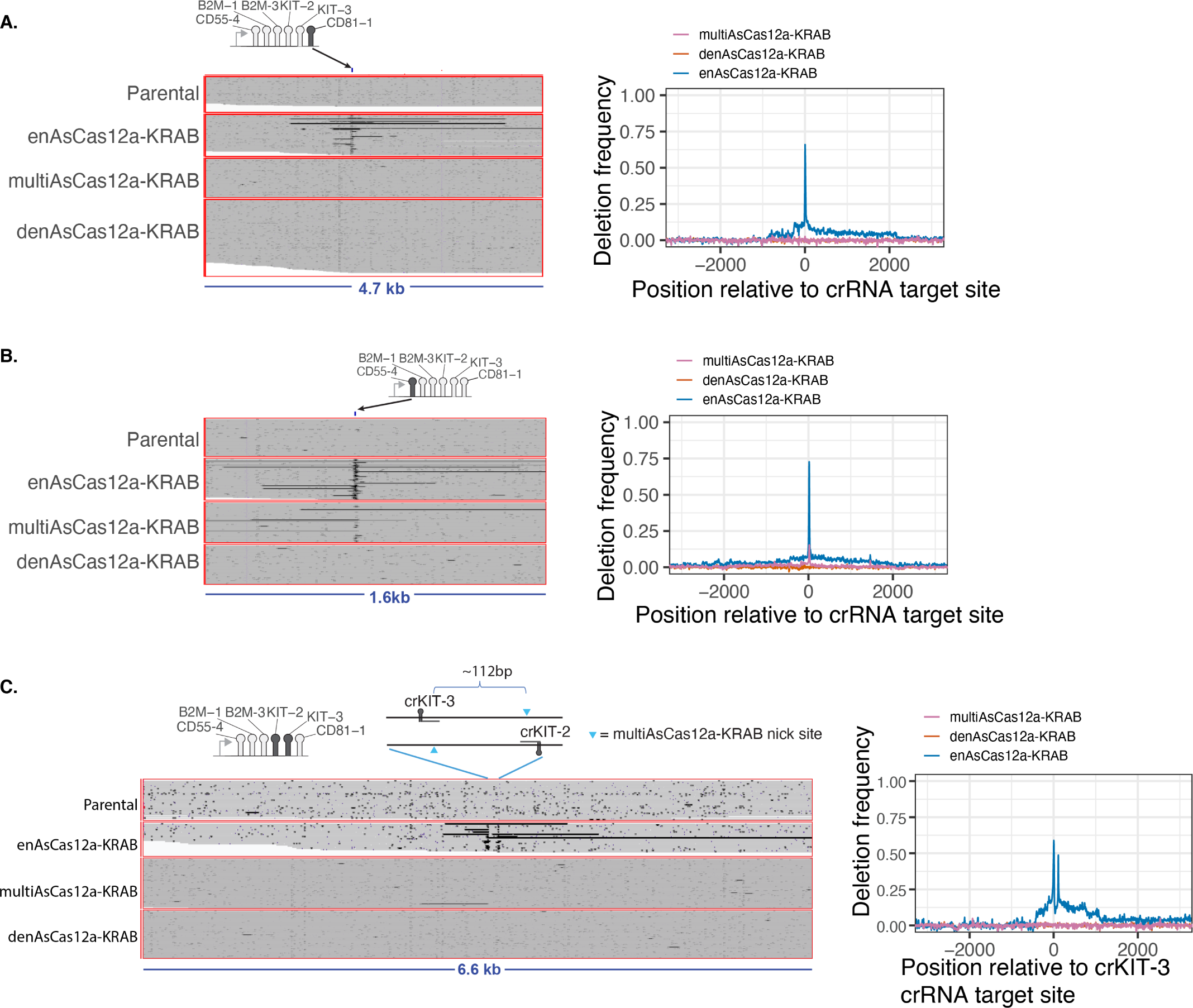
– Long-read sequencing quantification of deletions surrounding single and dual crRNA target sites. **A)** K562 cells piggyBAC-engineered to constitutively express the indicated fusion protein constructs were lentivirally transduced with the indicated 6-plex crRNA constructs at MOI *<* 0.13, sorted for transduced cells based on GFP reporter fluorescence, and the genomic DNA harvested 16 days later and subjected to long-read Nanopore sequencing. Left: Representative view of individual reads aligned to the region surrounding the crCD81-1 target site is shown, with deletions indicated by black horizontal lines. Right: Quantification of deletion frequency across the region surrounding the crCD81-1 target site. **B)** Analogous to A), but shown for region surrounding the crCD55-4 target site. **C** Analogous to A), but shown for region surrounding the crKIT-3 and crKIT-2 target sites located on opposing DNA strands spaced 112bp apart.

**Figure S17.**
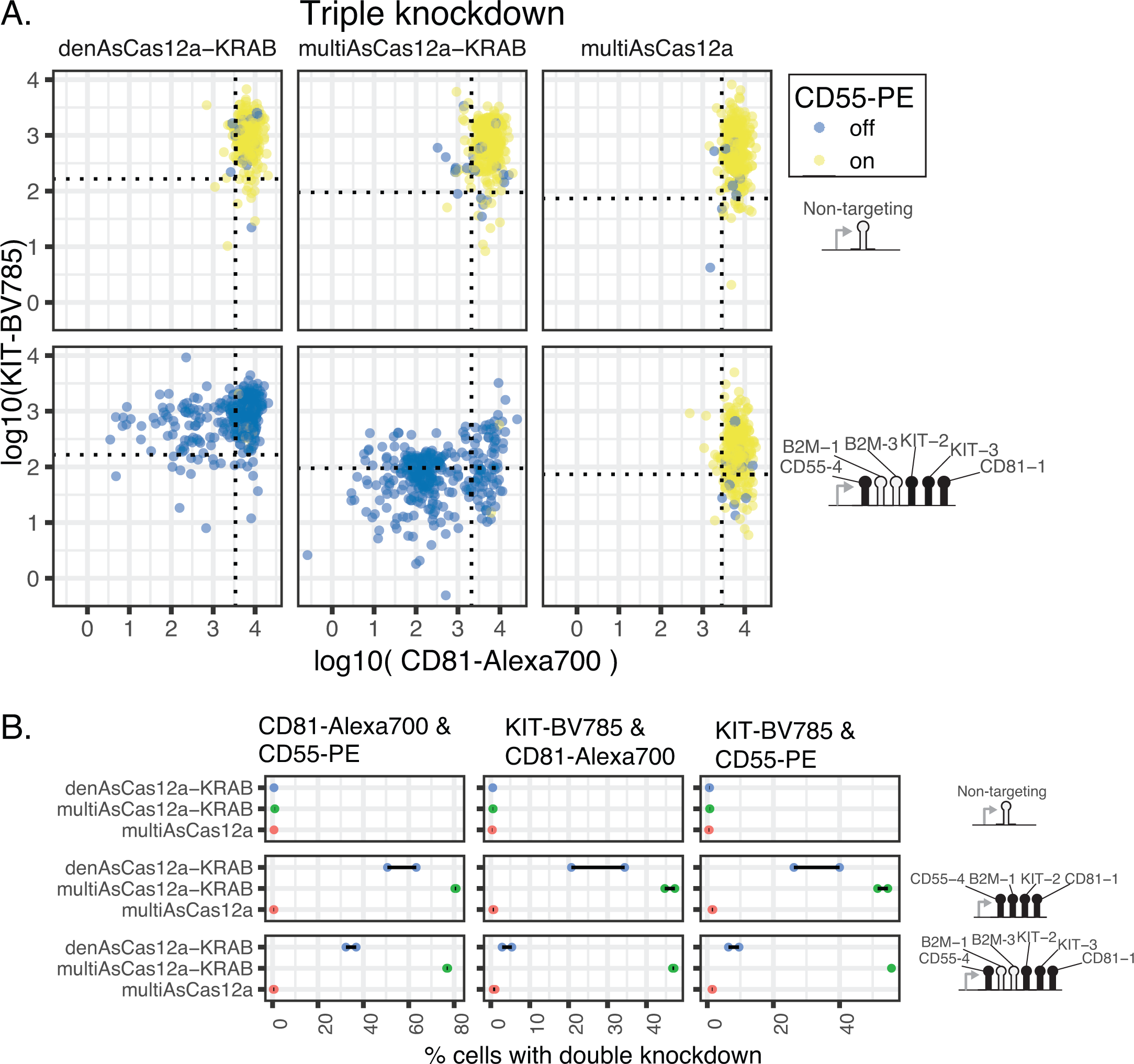
– Double and triple gene knockdown by CRISPRi using higher-order crRNA arrays. **A)** Single-cell view of CD81, KIT, and CD55 3-way knockdown using a 6-plex crRNA construct in K562 cells piggyBac-engineered to constitutively express each of the indicated fusion protein constructs, measured by multiplexed flow cytometry. Summary of percentage of cells with triple knockdown is shown in Fig. 3G. **B)** Quantification of the fraction of cells showing double-knockdown of pairs of target genes in the experiment described in A for 4-plex and 6-plex crRNA arrays. Double knockdown is defined as the fraction of cells with expression below the 5th percentile for the non-targeting crRNA for the expression of a given pair of target genes.

**Figure S18.**
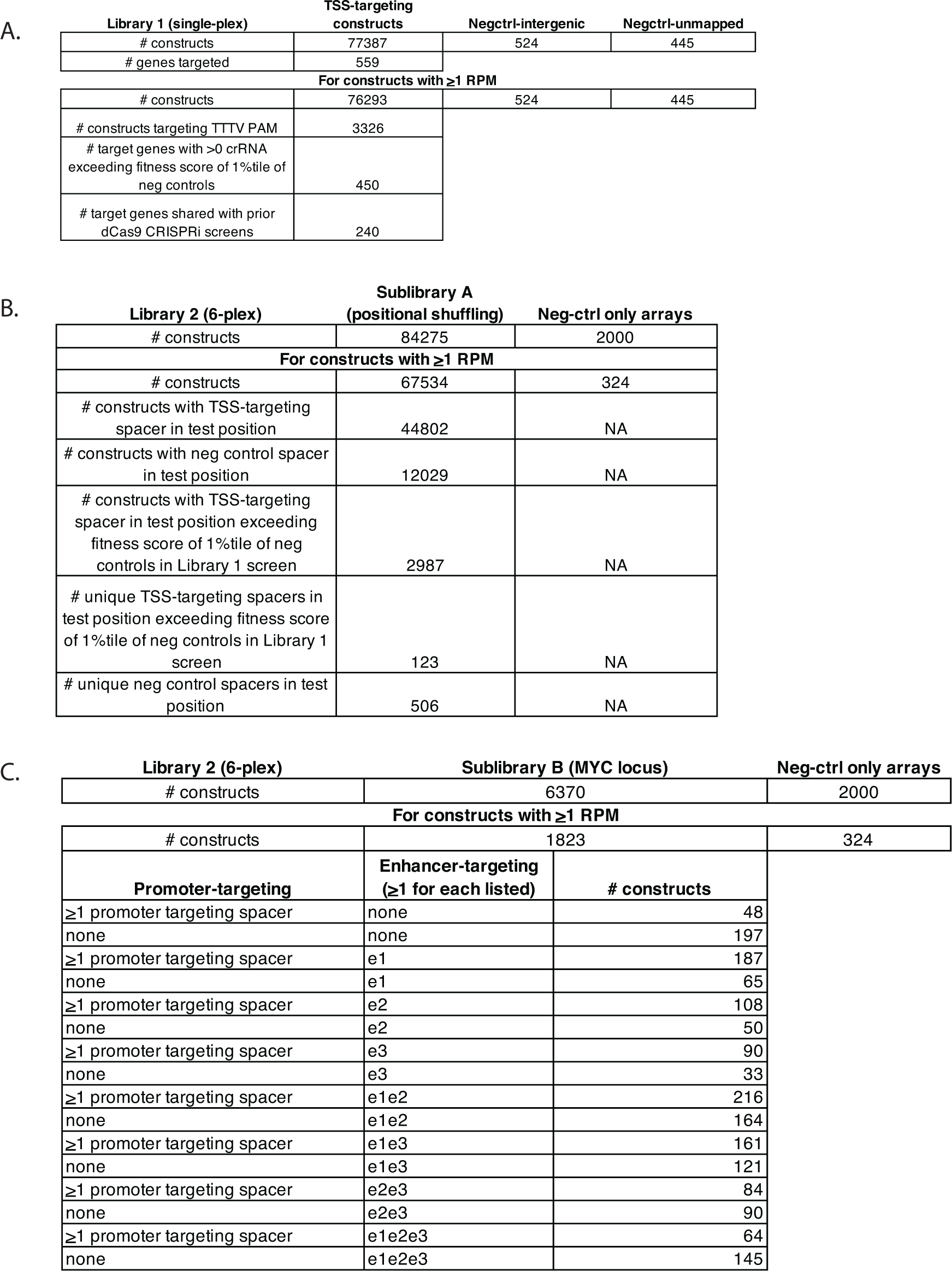
– Summaries of Library 1 and Library 2 screens.

**Figure S19.**
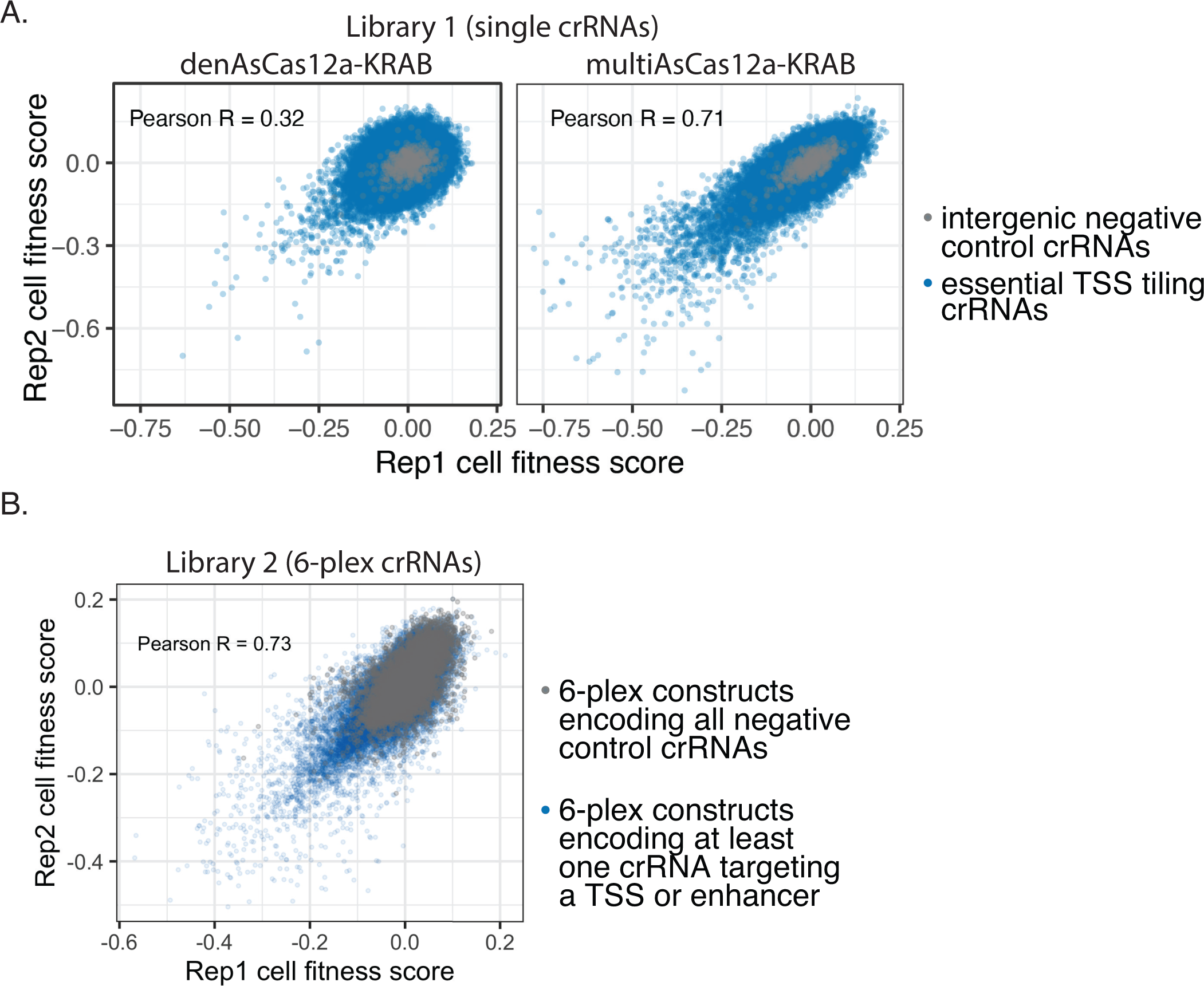
– Screen replicate concordance for Library 1 and Library 2. Shown are 2D density plots of cell fitness scores for individual crRNA constructs in **A)** Library 1 and **B)** Library 2 (Sublibrary A and Sublibrary B), with the Pearson correlation coefficients calculated for all constructs in each library shown.

**Figure S20.**
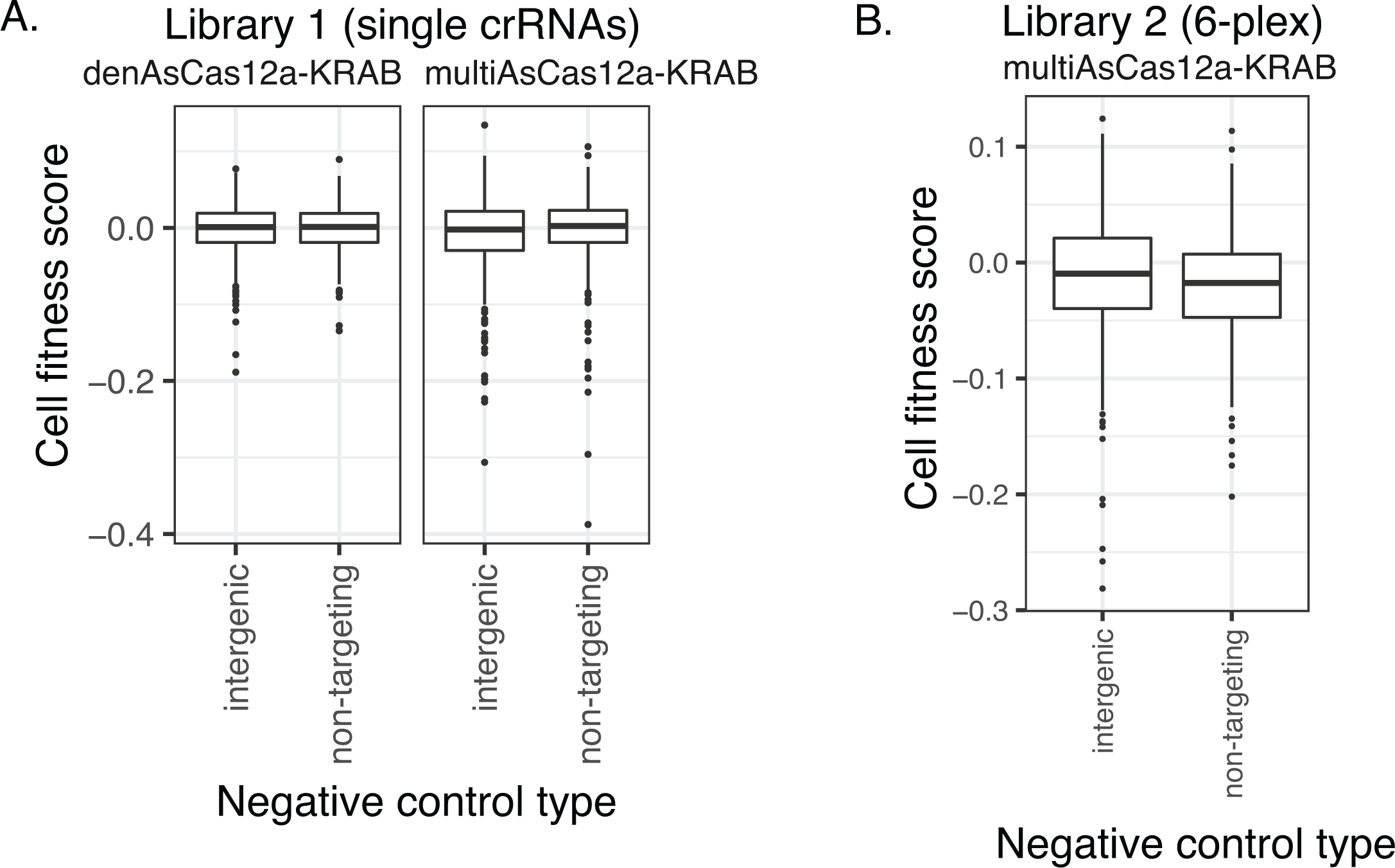
– Cell fitness score distributions of intergenic vs. non-targeting negative control crRNAs. Boxplots of cell fitness scores for **A)** Library 1 single-crRNA constructs, and **B)** Library 2 6-plex constructs, categorized by whether the construct encodes exclusively intergenic vs. non-targeting negative control crRNAs. Boxplots display median, interquartile range, whiskers indicating 1.5x interquartile range, and outliers.

**Figure S21.**
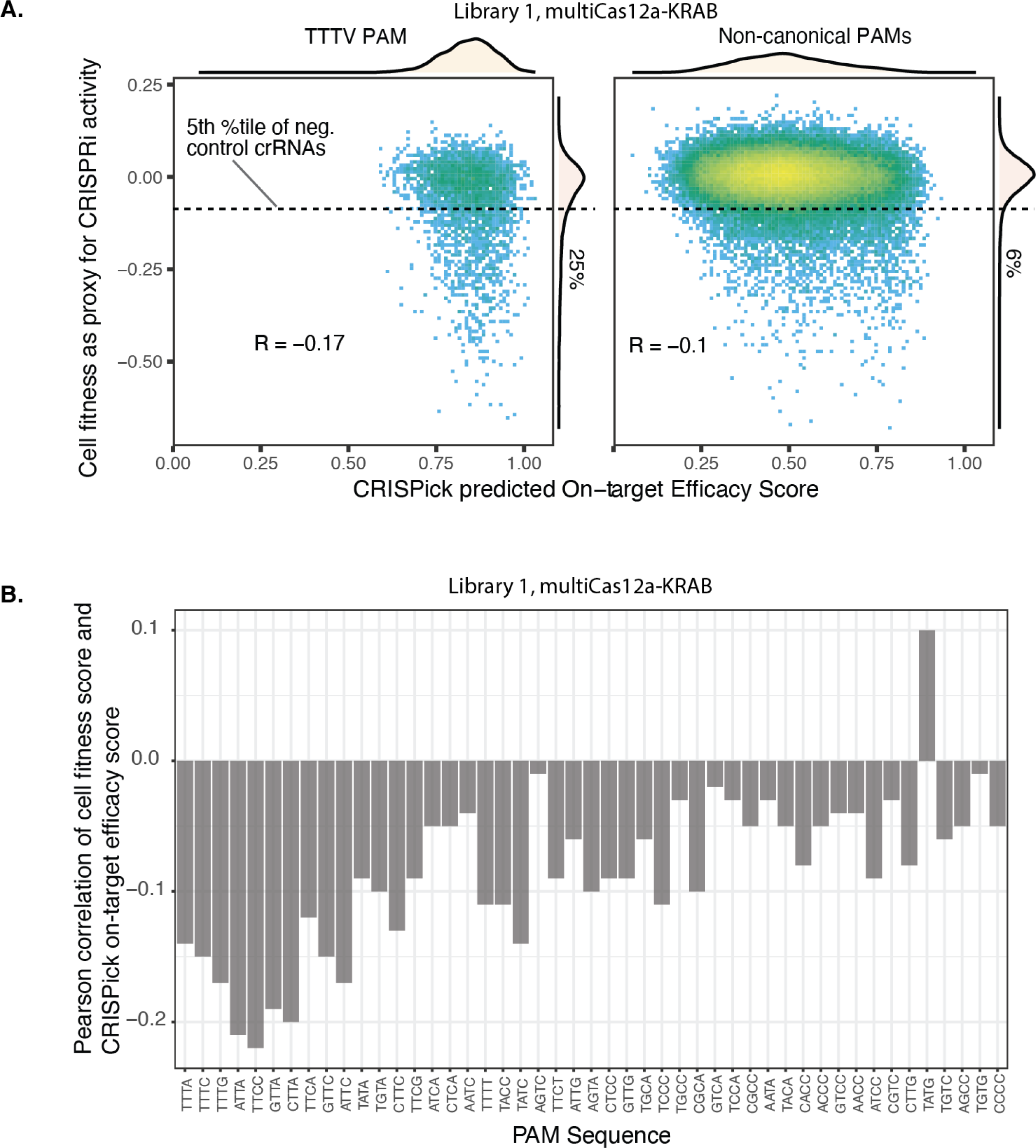
– Relationship between CRISPick predictions and empirical CRISPRi activity. **A)** Library 1: 2D density plots of cell fitness scores of individual crRNAs for multiAsCas12a-KRAB vs. predicted crRNA on-targeting efficacy score from the CRISPick algorithm, grouped by TTTV PAM vs. non-canonical PAM’s. The 5th percentiles of intergenic negative control crRNAs cell fitness scores are shown as a dashed horizontal line and the percentage of crRNAs below that threshold shown in the marginal histogram. Pearson correlation coefficients are shown. **B)** Library 1: Pearson correlation coefficient between the cell fitness score and the CRISPick on-target efficacy score is shown for each individual PAM sequence for multiAsCas12a-KRAB.

**Figure S22.**
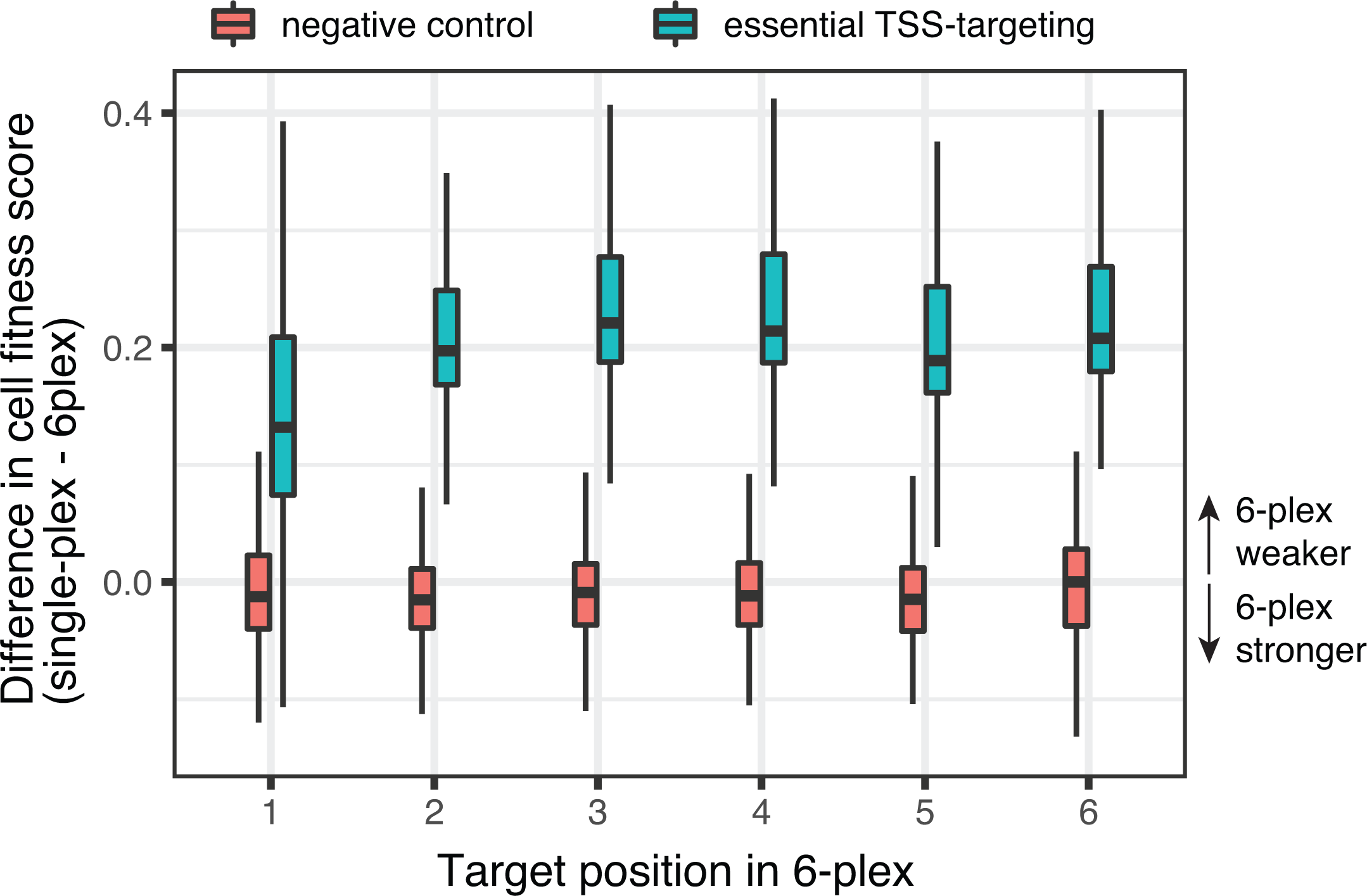
– Difference in cell fitness scores for 6-plex vs. 1-plex crRNA constructs. Library 2 Sublibrary A (6-plex crRNA construct with one test position targeting essential TSS) screen cell fitness scores for a given test position spacer are compared to the cell fitness scores of the same spacer in the Library 1 (single-plex crRNA) screen. Difference in cell fitness scores (6-plex minus single-plex) are shown as boxplots, which display the median, interquartile range, and whiskers indicating 1.5x interquartile range.

## Notes

### Summary of Updates

Text modified for brevity. Supplemental figures updated to include long-read Nanopore sequencing analysis.

